# Intravital single-cell behavior profiling reveals disrupted germinal center B cell motility and interactions by EZH2 gain-of-function mutation

**DOI:** 10.64898/2026.09.02.748910

**Authors:** Chanhong Min, Kibaek Choe, Xi Chen, Ioannis Karagiannidis, Nikita Sivakumar, Chris Xu, Ari M. Melnick, Jude M. Phillip, Wendy Béguelin

## Abstract

Germinal center (GC) B-cells give rise to the majority of non-Hodgkin lymphomas, underscoring the need to pinpoint critical processes that initiate and drive lymphomagenesis. Lymphoma driver mutations can alter GC B cell functions and B cell fate decisions. Here, we studied how EZH2 oncogenic mutation in GC B cells alters cellular motility and interactions with T follicular helper (Tfh) cells and follicular dendritic cells (FDCs) to determine B cell fate. By combining intravital imaging, single-cell behavior analyses, and RNA sequencing, we uncover how lymphoma-associated EZH2 mutations reprogram the behaviors of GC B cells in vivo. We found that EZH2 mutations increased single-cell motility speeds and morphological plasticity of GC B cells, redirecting migration toward the FDC–rich light zone subregions rather than to the dark zone. Although mutant EZH2 GC B cells exhibited normal engagement quality with FDCs, they showed shorter interaction times and reduced surface engagement with Tfh cells. Notably, EZH2 mutant B cells required prior contact with FDC before engaging with Tfh cells, thus impairing DZ recycling. This motility phenotype scaled with local mutant clone abundance, suggesting a behavioral strategy underlying how mutant cells outcompete WT cells. Lastly, we developed scMOTIPh, a computational framework that integrates single-cell behavioral features with transcriptomic profiles. Applying scMOTIPh to mutant GC B cells within the FDC-rich zone revealed enhanced ATP production, metabolic and antigen-presentation programs, and suppression of cell-death pathways, which is consistent with a tendency for malignant transformation and survival fitness. These findings provide an in vivo, single-cell view of how an epigenetic lesion rewires the local microenvironment by modulating single-cell behaviors within native GCs, revealing a dynamic mechanism for early lymphomagenesis.

**Teaser:** EZH2 mutation alters the emergent behaviors and interaction dynamics of germinal center B cells

## INTRODUCTION

The dynamic behaviors of immune cells are central to the proper functioning of germinal centers^1,2^. Within germinal centers (GC), B cells migrate between the dark (DZ) and light (LZ) zones, guided by interactions with follicular dendritic cells (FDCs) and T follicular helper (Tfh) cells that coordinate receptor affinity-based selection^3–8^. The histone methyltransferase EZH2 is required for the formation and maintenance of GCs. Somatic gain-of-function mutations in EZH2 occur in ∼25–30% of follicular lymphoma (FL) and germinal center B-cell like diffuse large B-cell lymphoma (GCB-DLBCL)^9–12^. These mutations enhance the conversion of H3K27me2 to H3K27me3, a histone mark associated with repression of transcription. EZH2-mutant GC B cells fail to properly recycle from the LZ back to the DZ, disrupting the canonical cyclic re-entry program that underlies affinity maturation. EZH2 mutations initiate lymphomagenesis by silencing genes critical for immune synapse signaling, reducing GC B-cell dependency on T-cell help, while also increasing dependence on FDC-derived signals^13,14^. In established lymphomas, EZH2 promotes immune evasion by dampening immunogenicity, thereby hindering tumor-T cell interactions^14^. This impaired recycling and modified interactions imply that EZH2 mutations fundamentally alter their dynamic behaviors (i.e., their motility and interactions) within the GC microenvironment. However, it remains unknown how EZH2 mutations in GC B cells affect the single-cell motility and dynamic interactions with Tfh cells and FDCs in vivo.

The dynamics processes that govern GC reactions in vivo have been illustrated through intravital microscopy studies^8,15,16^. Most GC B cells were shown to exhibit continuous motility with a net directional flow from DZ to LZ^6–8,15^. Beyond locomotion, intravital studies have visualized B–Tfh cell surface engagement that drives positive selection^16–18^. Additional studies suggested that GC B-cell speed, and the frequency and/or quality of contacts with FDCs influence GC output, underscoring the assumption that single-cell motility and interaction dynamics determine B-cell fate^19^. However, these studies profiled only a small set of features (e.g., speed, turning angle, contact duration/frequency), without providing higher-order information that describes the structure of movement trajectories and interaction networks that likely contribute to the encoding of specific cellular functions^20^. Moreover, previous studies have not reported how GC B cell motility and microenvironment interaction dynamics are perturbed by lymphoma-associated mutations at single-cell resolution. Unlike static molecular or transcriptional states, dynamic single-cell behaviors capture how B cells navigate, engage with neighboring cells, and transition between the GC microenvironment, thereby reconstructing the full sequence of immune events that govern B cell selection and fate.

Another open challenge is linking these dynamic behaviors to their molecular mechanisms to identify dysregulated gene programs and establish therapeutically relevant pathways^21^. Live imaging provides spatiotemporal measurements of functional behavior, whereas conventional scRNA-seq provides genome-wide molecular profiles^22^. However, these modalities often remain unpaired, limiting the ability to determine how specific transcriptional programs are manifested as cellular behaviors in vivo^23,24^. Their integration is therefore essential for connecting GC B cell behavior to the molecular programs that govern their migration, selection, and malignant transformation.

To address this gap, we combined intravital two-photon microscopy with single-cell behavioral and transcriptomic analyses to develop scMOTIPh, a computational framework that maps unpaired behavioral and transcriptomic profiles through shared functional programs. This approach allows us to define how EZH2 mutation reshapes GC B cell dynamics in vivo. By resolving the single-cell behavior changes at high temporal resolution within intact GCs and linking these behaviors to transcriptional programs, our study elucidates a dynamic, mechanistic view of early lymphomagenesis. Moreover, these results establish an integrative cellular framework for identifying functional signatures of malignant cells that are linked to disrupted time-dependent behaviors, providing a basis for novel therapeutic strategies.

## RESULTS

A combined intravital microscopy and multidimensional single-cell behavioral profiling framework enables high-resolution mapping of GC B cell dynamics in vivo

We conducted intravital 2-photon (2P) microscopy imaging of popliteal lymph nodes in mixed bone marrow chimeric mice to investigate the intricate dynamics of GC B cells interacting with follicular dendritic cells (FDC) and T follicular helper (Tfh) cells within the GC. This allowed us to determine how GC B cell dynamics were altered by the lymphoma-associated gain-of-function EZH2 mutation. Transplantation experiments used the Cγ1-cre strain^25^ to drive GC-specific expression of *Ezh2*^Y641F^ from the endogenous *Ezh2* locus^26^, mutant EZH2 (MT EZH2). WT, MT EZH2 GC B cells and Tfh cells were fluorescently labeled with CFP, YFP, and tdTomato, respectively. The combined bone marrow cells were transplanted into lethally irradiated C57BL6 recipient mice. Once engrafted, the recipient mice were immunized with NP-OVA, and popliteal lymph nodes were imaged at the peak of GC reactions, 9-11 days post-immunization (**Figure 1a**, panel 1). One day before imaging, transplanted mice were injected with AlexaFluor594-conjugated anti-CD35 antibody to label FDCs. Intravital 2P microscopy was conducted for 90 minutes per GC, capturing 3D volumes (230 x 230 x 90 µm^3^) every 30 seconds (**Figure S1**, **Supplementary Video 1**). We optimized imaging conditions across all four fluorophores (**Figure S2a**) to maximize fluorescence signal (**Figure S2a-c**) while preventing laser-induced damage^27–29^.

**Figure 1.**
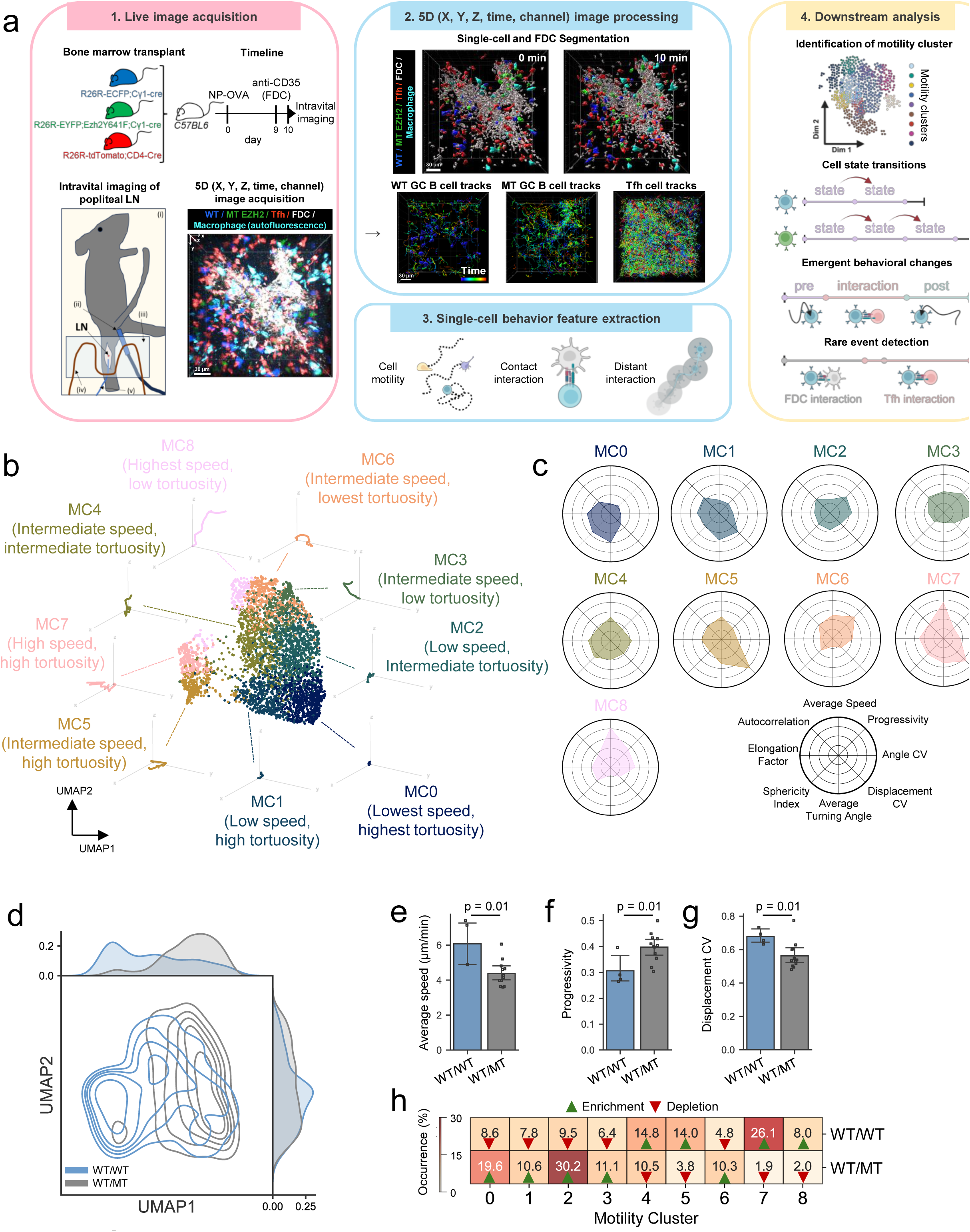
Intravital microscopy and multidimensional single-cell behavior analysis reveal that the presence of mutant GC B cells reshapes the motility patterns of WT GC B cells. **a.** Schematic illustration of the live-image generation, image processing, and single-cell behavior analysis pipeline. 1. Multicolor GC mouse models were generated via bone marrow transplantation using R26R-ECFP;Cγ1-cre, R26R-EYFP;Ezh2^Y641F^;Cγ1-cre, R26R-tdTomato;CD4-cre donor mice. FDCs were labeled in vivo with AlexaFluor-conjugated anti-CD35 antibody. 9-11 days after the injection of NP-OVA, intravital 2P microscopy was conducted to capture 3D dynamic behaviors of GC cells. 2. 3D segmentation was performed for each channel, followed by single-cell tracking of WT, MT GC B cells, and Tfh cells. 3. Motility features were extracted, including speed, turning angle, and displacement and colocalization features were quantified based on the spatiotemporal proximity and contact dynamics with FDCs and Tfh cells. 4. Downstream analysis included the identification of motility clusters using unsupervised clustering, cell-state transition analysis based on the longitudinal record of each single-cell, emergent behavior change of GC B cells pre/post contact with Tfh cells, and spatiotemporal description of tri-interaction. **b.** UMAP representation of unsupervised k-means clustering identifies nine unique motility clusters (MCs, MC0 n = 697, MC1 n = 463, MC2 n = 825, MC3 n = 386, MC4 n = 561, MC5 n = 301, MC6 n = 404, MC7 n = 240, MC8 n = 192 10-minute track of cells). **c.** Average Z score quantified for interpretable motility features per MC. **d.** Two-dimensional UMAP kernel density estimation (KDE) representation of multidimensional single-cell motility computed for each WT from WT-only GCs (WT/WT) and WT coexisting with MT EZH2 cells (WT/MT) (WT/WT n = 973 and WT/MT n = 1,482 10-minute track of cells). Top, 1D KDE across UMAP1. Right, 1D KDE across UMAP2. e-g. Quantification of average speed (e), progressivity (f) and displacement coefficient of variance (CV) (g) within 10 min duration per GC (mean ± 95% C.I.). Each dot represents one GC, with values averaged across all cells within that GC. P values are from Mann-Whitney U test (WT/WT n = 4, WT/MT n = 12 GCs, 112-352 tracks/GC). h. Fractional occurrence of nine MCs across WT/WT and WT/MT. Enrichment and depletion represent statistical over-representation and under-representation compared to random sampling of GC B cells, respectively.

Following image acquisition, we applied a 5-dimensional (5D: X, Y, Z, time, channel) image processing framework, developed within our in-house computational pipeline^20,30^ to analyze the collected behavioral data. We applied single-cell segmentation and tracking using commercial software (IMARIS) to reconstruct cell morphologies and trajectories for WT and MT EZH2 GC B cells, as well as Tfh cells (**Figure 1a**, panel 2; **Supplementary Video 2**). Cell trajectories were normalized by dividing them into 10-minute trajectories, allowing for consistent comparison of cellular behaviors across time and across cell types. To characterize single-cell behaviors, we first registered cell trajectories and then extracted 123 motility features to capture detailed movement patterns. Additionally, we quantified spatial proximities and interactions between GC B cells, Tfh cells, and the FDC network using 135 colocalization features (**Figure 1a**, panel 3; **Supplementary Table 1 and 2**). To gain a detailed understanding of GC B cell behaviors, we performed downstream analyses of motility and colocalization features extracted across all cell trajectories (**Figure 1a**, panel 4). To quantify the heterogeneous motility behaviors, we applied unsupervised k-means clustering, which grouped thousands of individual GC B cells into distinct behaviors using a modified version of the CaMI pipeline^20^ (see Methods). This spatio-temporal analysis allowed us to quantify transitions between different behaviors, evaluate the temporal variability of immune events, and observe changes in B cell behavior within the GC microenvironment. By analyzing at single-cell resolution, we were able to detect rare events within the GC, such as interactions involving three cell types: GC B cells, Tfh cells, and FDCs. Notably, this type of detail was not achievable using analyses of population averages.

Here, we characterized distinct motility patterns of GC B cells, revealing nine distinct motility clusters (MC) across 4,069 GC B cell trajectories. Each motility cluster represented a unique set of movement characteristics within the GC (**Figure 1b, Figure S3a-b**). For example, MC0 was characterized by the lowest speed, indicating minimal movement, a high average turning angle, and a high sphericity index, representing tortuous movement and circular trajectories (**Figure 1c, Figure S3c**). In contrast, MC8 displayed the highest speed, suggesting rapid and extensive movements across the GC. MC2 represented a distinctive motility profile, with low speed and low turning angle, implying a directed migration pattern over short distances. The overall distribution of MCs was consistent across days 9–11 post-immunization and between fluorophore-swapped bone marrow chimeras, showing a cross-configuration correlation of 0.85 (**Figure S3d**).

### The presence of mutant EZH2 GC B cells reshapes the motility patterns of WT GC B cells

To determine whether these motility patterns reflect an intrinsic property of WT GC B cells or are influenced by the presence of mutant GC B cells, we performed an additional experiment in which both fluorophore-labeled populations were WT, approximating a purely physiological GC environment. Surprisingly, WT GC B cells from WT-only GCs (WT/WT) occupied a distinct region of the single-cell motility space compared to WT cells coexisting with MT EZH2 cells (WT/MT) (**Figure 1d**). In the WT-only setting, cells exhibited significantly higher speeds (**Figure 1e**), lower progressivity (**Figure 1f**), and greater displacement coefficient of variance (CV) (**Figure 1g**), reflecting faster yet more tortuous movement with highly fluctuating displacement (high transition between low speed and high speed), respectively. This shift was evident at the level of MC composition. WT-only cells were enriched in MC7, characterized by high speed and high displacement CV, and increased abundance of cells in MC4, MC5, and MC8, all of which correspond to highly motile states (**Figure 1h**). Together, these results indicate that WT GC B cell motility is strongly influenced by the presence of MT EZH2 cells, suppressing physiologically high motility behaviors.

### MT EZH2 GC B cells are characterized by increased motility and spatiotemporal heterogeneity

We investigated the effects of EZH2 mutation on the motility patterns of GC B cells. MT EZH2 GC B cells traveled consistently greater distances across all observed time intervals compared to WT GC B cells (**Figure 2a**). Further analyses were performed using standardized 10-minute trajectories to ensure comparable sampling and statistical power. Representative trajectories normalized to the origin confirmed the enhanced motility of MT EZH2 GC B cells, which exhibited greater displacement across the GC microenvironment (**Figure 2b, c**). Consistently, quantifying the average speed of WT and MT GC B cells per GC showed that MT GC B cells exhibit significantly higher average speeds than WT (**Figure 2d**). We observed no significant differences between WT and MT GC B cells in progressivity that measures directional persistence of B cell movement (**Figure 2e**).

**Figure 2.**
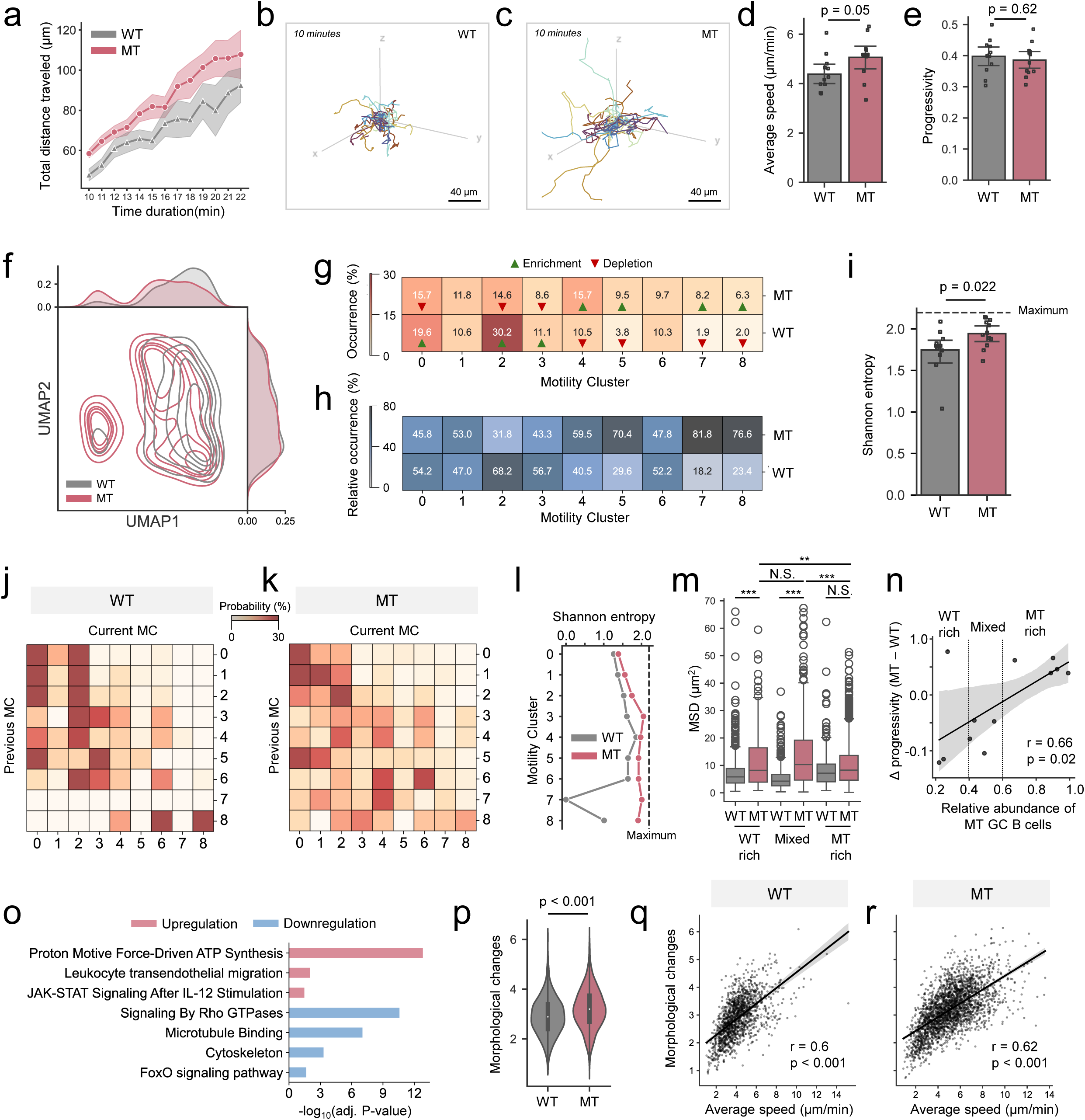
Characterization of WT and MT EZH2 GC B cell motility patterns in single-cell. **a.** Total distance traveled over the time duration with 30s bin (mean ± 95% C.I.). **b-c.** Thirty example movement trajectories from each WT (**b**) and MT (**c**) GC B cells. Each color represents a unique track. **d-e.** Quantification of average speed (**d**) and progressivity (**e**) within 10 min duration per GC (mean ± 95% C.I.). Each dot represents one GC, with values averaged across all cells within that GC. P values are from Mann-Whitney U test (WT n = 12, MT n = 12 GCs, 161-561 tracks/GC). **f.** Two-dimensional UMAP kernel density estimation (KDE) representation of multidimensional single-cell motility computed for each WT and MT GC B cells (including WT n = 1,482 and MT n = 2,587 10-minute track of cells). Top, 1D KDE across UMAP1. Right, 1D KDE across UMAP2. **g.** Fractional occurrence of nine MCs across WT and MT GC B cells. Enrichment and depletion represent statistical over-representation and under-representation compared to random sampling of GC B cells, respectively. **h.** Column-wise normalized relative occurrence of nine MCs across WT and MT GC B cells. **i.** Quantification of Shannon entropy of fraction occurrence of nine MCs per GC (mean ± 95% C.I.). Each dot represents one GC. P value is from Mann-Whitney U test (WT n = 12 and MT n = 12 GCs, including 161-561 tracks/GC). **j-k.** Matrix of transition from previous MC (row) to current MC (column) computed for each WT (**j**) and MT GC B cells (**k**). Each row is normalized to sum 100%. Transition to MC7 does not exist in WT. **l.** Shannon entropy computed for each previous MC (row-wise) in the transition matrix. Transition to MC7 does not exist in WT. **m.** Mean squared displacement (MSD) of GC B cells stratified according to the relative abundance of MT GC B cells as WT rich (<0.4), Mixed (0.4–0.6) or MT rich (>0.6). P value is from one-way ANOVA followed by Tukey’s post-hoc test (WT WT rich n = 592, MT WT rich n = 186, WT Mixed n = 656, MT Mixed n = 525, WT MT rich n = 234, MT MT rich n = 1876 cells; **p<0.01, ***p<0.001, N.S. means non-significant). Each point represents one cell. Line inside box represents median, boundaries of the box represent first and third quartile. Lines outside box represent whiskers. **n.** Difference in median progressivity between MT and WT GC B cells within each GC against the relative abundance of MT GC B cells. r indicates Pearson correlation coefficient with corresponding p value (12 GCs). Each dot represents a GC. The line indicates the linear regression fit, and shading denotes the 95% confidence interval. Vertical dotted lines indicate the thresholds used to define WT-rich, mixed and MT-rich GCs. **o.** Gene Set Enrichment Analysis (GSEA) in centrocytes based on bulk-RNA seq. **p.** Quantification of cellular average morphological changes within a 10-min duration. Each point represents 10-minute track of cell. White dot represents median, boundaries of black box represent first and third quartile. P value is from Mann-Whitney U test (WT n = 1,482, MT n = 2,587 10-minute track of cells). **q-r.** Correlation between average speed and morphological changes per WT (**q**) and MT GC B cells (**r**). r indicates Pearson correlation coefficient with corresponding p value (WT n = 1,482, MT n = 2,587 10-minute track of cells). Each dot represents 10-minute track of cell.

Beyond bulk trend, single-cell motility analysis revealed distinct motility patterns between WT and MT GC B cells, particularly along the UMAP 1 axis (**Figure 2f**). Analyzing the fractional abundance of GC B cells across the MCs showed that WT GC B cells were predominantly localized in MC0 and MC2, which exhibited lower mobility patterns (**Figure 2g**). Conversely, MT GC B cells were significantly enriched in MC4, MC5, MC7, and MC8, clusters characterized by higher motility. Given the similar number of WT and MT cells, 76.6% of GC B cells in MC8 were EZH2 MT GC B cells, whereas 68.2% of GC B cells in MC2 were from WT GC B cells (**Figure 2h**). To quantify cellular heterogeneity among MT and WT GC B cells, we computed the Shannon entropy. Higher Shannon entropy indicates greater heterogeneity, meaning that cells are more evenly distributed across the MCs. In contrast, lower Shannon entropy indicates less heterogeneity, which cells showing more homogeneous behaviors or preferentially occupying only a few MCs. Results revealed a significantly higher cellular heterogeneity for MT GC B cell motility compared to WT, reflecting the ability of MT GC B cells to adopt a broad range of movement strategies (**Figure 2i**).

Furthermore, we performed transition analysis of MC, which quantifies changes in a cell’s MC identity across consecutive time windows within the same trajectory. This captures how cells switch between distinct movement patterns over time. Results revealed relatively consistent transitions among WT GC B cells, primarily towards low-motility clusters MC0 and MC2 (**Figure 2j**). In contrast, MT GC B cells exhibited more variable and heterogeneous transitions across multiple MCs beyond MC0 and MC2 (**Figure 2k**). This was also reflected in the elevated transition entropy of MT GC B cells across all MCs, especially those originating from MC3 to MC8, denoting intermediate-to-high motility clusters (**Figure 2l**). These findings suggest that the EZH2 mutation not only enhances the speed capability of GC B cells but also expands the movement strategies by increasing the MC switching, indicating plastic motility programs that reshape how cells explore GC microenvironment.

Given our earlier finding that the presence of MT GC B cells reshapes WT GC B cell motility (**Figure 1d-h**), we next investigated whether variation in the relative abundance of MT GC B cells across individual GCs contributed to this motility heterogeneity. Although the chimeras were generated using an intended 1:1 WT:MT ratio, MT cell abundance varied across individual GCs, which were classified as WT-rich (MT abundance < 0.4), mixed (0.4 < MT abundance < 0.6) or MT-rich (MT abundance > 0.6). MT cells showed greater mean squared displacement (MSD) than WT cells in WT rich and mixed GCs, whereas WT and MT displacement was comparable in MT rich GCs (**Figure 2m**). Moreover, the difference in progressivity between MT and WT increased as MT cell abundance rose (**Figure 2n**). This pattern suggest a shift from the high-displacement, tortuous movements in WT-rich GCs to the lower-displacement, more directionally persistent movement in MT-rich GCs. Together, these findings indicate that the relative abundance of MT GC B cells contribute, at least in part, to heterogeneity in the motility phenotype of GC B cells.

### High motility of MT EZH2 GC B cells is linked to enhanced morphological changes and motility-associated gene signatures

To investigate the molecular programs associated with the altered motility of MT GC B cells, we analyzed bulk RNA-seq of sorted WT and MT centroblasts and centrocytes^13^. Differential gene expression analysis revealed 449 down- and 148 upregulated genes in MT EZH2 centroblasts, and 671 down- and 189 upregulated genes in MT EZH2 centrocytes (**Figure S4a-b, Supplementary Table 3 and 4**). Specifically, centrocyte gene set enrichment analysis (GSEA) using the Molecular Signatures Database (MSigDB) revealed a significant downregulation of cytoskeleton and microtubule pathways in MT GC B cells (**Figure 2o, Supplementary Table 5**). Downregulation of cytoskeleton related genes in MT GC B cells implies decreased mechanical stiffness, or increased deformability. This is further supported by the strong downregulation of Rho GTPase signaling genes (e.g. TIAM1, DAAM1, ARHGEF25) (**Figure S4c**), a key regulator of cytoskeleton formation^31^. This trend was not unique in centrocytes, as corresponding pathways were also downregulated in MT EZH2 centroblasts (**Figure S4d**). This suggests increased cellular deformability and an upregulation of leukocyte trans-endothelial migration pathways, which would require substantial morphological changes to allow cells to squeeze and traverse endothelial barriers. Further, MT GC B cells also exhibited upregulation of ATP synthesis, indicative of elevated metabolic activity, which is reported to be associated with enhanced lymphocyte or cancer cell migration^32^ (**Figure 2o**).

To link the transcription-associated deformability pathway results with the imaging data, we assessed the morphological changes of individual GC B cells by constructing a multi-dimensional morphology space and measuring the average displacement of each cell’s morpho-trajectory, which captures the magnitude of dynamic shape changes over time. This analysis revealed that MT GC B cells display significantly higher morphological changes relative to WT, representing high cellular deformability (**Figure 2p**). In addition, morphological changes were highly correlated with the speed in both WT and MT GC B cells (**Figure 2q-r**), indicating that enhanced GC B cell motility is closely coupled with cellular morphological changes.

To further illustrate the transcriptional programs underlying enhanced motility in EZH2 MT GC B cells, we hypothesized that transcriptional determinants of enhanced migratory behavior may be shared between centroblasts and centrocytes, given that both MT EZH2 subtypes exhibited increased deformability and motility. Focusing on genes consistently up- or downregulated in both populations (**Figure S4b**), we identified a curated set of 49 motility-associated genes with known functions in chemotaxis, cytoskeletal organization, migration, polarity, and invasion (**Figure S5a, Supplementary Table 6**). To contextualize these genes within known migratory behaviors, we curated gene sets from the literature associated with 3D amoeboidal migration^33^ (**Figure S5b**), chemotaxis^33^ (**Figure S5c**), chemokinesis^33^ (**Figure S5d**), and morphological plasticity^34^ (**Figure S5e**) in the context of EZH2 MT GC B cells. Based on these gene sets, we utilized Louvain community detection to assess gene-level overlap across motility-related pathways, revealing tight modular association between genes in downregulation of Rho GTPase signaling and actin remodeling, and the partial gene overlap with 3D amoeboidal migration, chemotaxis, and chemokinesis programs (**Figure S5f**). Specifically, GSEA conducted on the motility-associated genes displayed downregulation of cytoskeletal and Rho GTPase signaling, implying increased cellular deformability (**Figure S5g**). Together, these results link transcriptional downregulation of cytoskeletal regulators with increased physical deformability and enhanced motility in MT GC B cells.

### Differential motility patterns of MT EZH2 GC B cells are spatially regulated relative to the FDC network

The observed differential motility patterns in MT EZH2 GC B cells suggested that the variations may stem from distinct spatial regulation within the GC. To investigate this, we identified the dark zone (DZ) and light zone (LZ) based on the presence of the FDC. We further subdivided the LZ into a sparse light zone (sLZ), which contains low FDC density, and a dense light zone (dLZ), which corresponds to the top 10% of the LZ volume with the highest FDC density (**Figure 3a**). Regions without FDCs were categorized as DZ. This spatial segmentation introduced nonlinear and biologically relevant boundaries within the GC to more accurately capture the complex architecture of the FDC network^35^.

**Figure 3.**
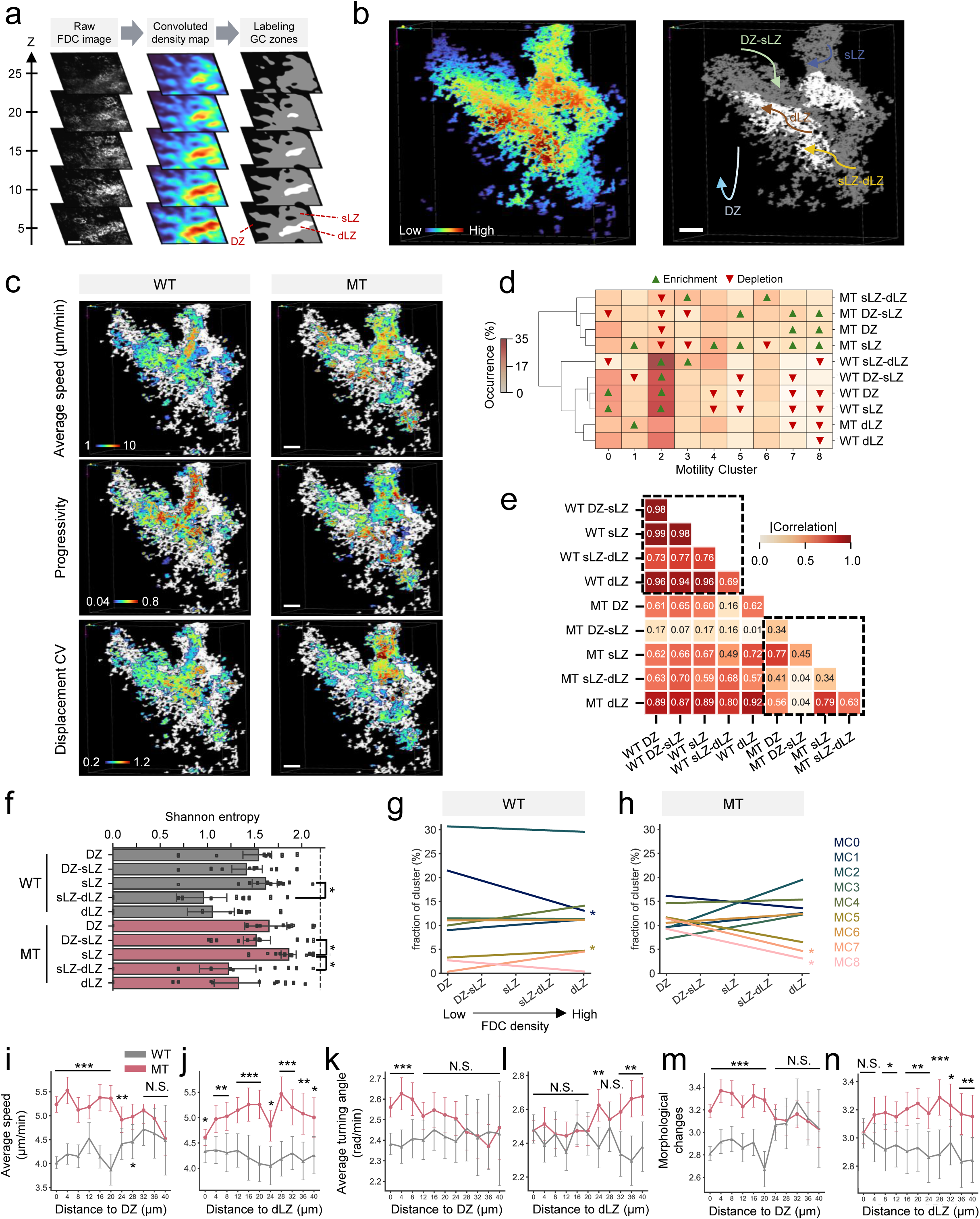
Differential motility patterns of MT EZH2 GC B cells are regulated by spatial location relative to the FDC network. **a.** Defining zones within GC based on FDC density map. 3D convolution is applied to FDC mask, followed by adaptive thresholding to identify DZ, sLZ and dLZ. Scale bar, 40 µm. **b.** (Left) 3D FDC density map. (Right) Labeling cell trajectories based on the duration of a cell’s presence in the zone. Black, DZ; gray, sLZ; white, dLZ. **c.** Projection of magnitude of motility features onto 3D FDC mask for WT (left) and MT (right). Scale bar, 25 µm. **d**. Heatmap of fractional occurrence of 9 MCs across WT and MT GC B cells in distinct zones. Hierarchical clustering is based on the Euclidean distance using Ward linkages. **e.** Cross-correlation between WT and MT GC B cells in zones based on the occurrence of MCs. Top left and bottom right dotted boxes represent correlation within WT and MT, respectively. Enrichment and depletion represent statistical over-representation and under-representation compared to random sampling of GC B cells, respectively. **f.** Shannon entropy computed for each WT and MT GC B cells in zones (mean ± 95% C.I.). Each dot represents one GC. P values are from two-sided unpaired t-test followed by Benjamini-Hochberg correction for multiple tests. p<0.05 (*). (WT DZ n = 10, WT DZ-sLZ n = 9, WT sLZ n = 12, WT sLZ-dLZ n = 12, WT dLZ n = 12, MT DZ n = 12, MT DZ-sLZ n = 12, MT sLZ n = 12, MT sLZ-dLZ n = 11, MT dLZ n = 12 GCs). **g-h.** Pearson’s correlation between MC cluster occurrence and FDC density computed for each WT (**g**) and MT GC B cells (**h**). p<0.05 (*). **i-j.** Change of average speed computed at each distance to DZ (**i**) and dLZ (**j**). **k-l**. Average turning angle computed at each distance to DZ (**k**) and dLZ (**l**). **m-n**. Morphological changes computed at each distance to DZ (**m**) and dLZ (**n**). **i-n**. Mean ± 95% C.I. Each point represents average value across cells within that distance bin. P values are from Mann-Whitney U test. p<0.05 (*); p<0.01 (**); p<0.001 (***). (Distance to DZ: WT n = 24 - 507 tracks, MT n = 74 - 720 tracks, Distance to dLZ: WT n = 70 - 192 tracks, MT n = 116 - 389 tracks)

To incorporate spatial regulation in our motility analysis, we labeled the movement trajectories of GC B cells according to the time they spent within each of the defined zones (**Figure 3b**). Specifically, cells spending more than 60% of the time in one zone were labeled according to that zone, otherwise trajectories were categorized as transition zones, such as DZ-sLZ or sLZ-dLZ. Using these definitions, we analyzed pairwise cell distances across zones and identified characteristic spatial length scales that defined the in vivo GC landscape. For example, GC B cells residing in the DZ–sLZ transition zone and dLZ were most frequently found at 2.97 µm and 41.66 µm from the DZ boundary, respectively (**Figure S6a**). Conversely, GC B cells located in the sLZ–dLZ transition zone and DZ exhibited peak distance probabilities at 3.34 µm and 45.73 µm from the dLZ boundary, respectively (**Figure S6b**). Tfh cells were heterogeneously distributed across the zones (dLZ 4.4%, sLZ-dLZ 6.8%, sLZ 53.6%, DZ-sLZ 12.4%, DZ 22.8%), mostly populating within sLZ at more than 50% (**Figure S6c**).

In these newly defined zones, we observed that the overall DZ and LZ boundaries did not significantly alter the motility patterns of either WT or MT GC B cells (**Figure S6d**). However, their motility patterns were strongly influenced by the local spatial arrangement within the FDC network, as evidenced by heterogeneous motility features across GC B cells (**Figure 3c**). Furthermore, the UMAP representation of the motility space revealed a distinct low UMAP1 “island” of motility clusters associated with MC5 and MC7, which was uniquely enriched in the sLZ of MT GC B cells, a pattern that was less prominent in WT (**Figure S6e**).

We quantified spatially regulated motility by examining MC distributions across GC zones. WT B cells showed uniform MC patterns, but MT B cells exhibited distinct, dynamic behaviors in the DZ, sLZ, and transitional zones that converged to WT-like motility in the dLZ. MT cells in the dLZ were depleted in MC7 and MC8, matching the WT pattern, but enriched for these clusters elsewhere (**Figure 3d**). Quantifying the similarity of MC distributions across zones, we calculated pairwise cross-correlations of MC frequencies within and between WT and MT (**Figure 3e**). WT GC B cells exhibited consistently high correlations across all zones (0.69 - 0.99), whereas MT GC B cells showed much lower inter-zonal correlation values (0.04 - 0.79), reflecting motility patterns with increased spatial heterogeneity. Notably, MT GC B cells in DZ-sLZ had very low correlations (0.01 – 0.17) with WT GC B cells in all zones, and also with MT GC B cells in sLZ-dLZ and dLZ, highlighting the distinct MC distribution of MT GC B cells in the DZ-sLZ transition state. MC distributions for MT GC B cells in the dLZ exhibited high similarity to WT cells in all zones (correlation range: 0.80–0.92), further supporting the assumption that MT cells adopt WT-like motility behavior upon entering the spatially constrained dLZ (**Figure 3e**).

Further, the heterogeneity of cells among distinct MCs changes according to the density of the FDC network. For example, we observed a significant decrease in motility state entropy during the transition from sLZ to sLZ-dLZ in both WT and MT GC B cells, indicating a decrease of behavioral diversity as cells approach the dense FDC network (**Figure 3f**). Conversely, transitions from DZ-sLZ to sLZ showed a marked increase in MT GC B cell heterogeneity, suggesting a recovery of diverse motility upon entering sLZ from the DZ-sLZ transition state. (**Figure 3f**). Notably, MT GC B cells in DZ-sLZ had very low correlations (0.01 – 0.17) with WT GC B cells in all zones, and also with MT GC B cells in sLZ-dLZ and dLZ, highlighting the distinct MC distribution of MT GC B cells in the DZ-sLZ transition state (**Figure 3e**). These findings indicate that EZH2 mutation enhances regional difference in GC B cell motility and induces distinct motility patterns compared to WT, except in the dLZ.

To further examine how local FDC density influences motility state composition, we quantified fractional changes in each MC across zones of increasing FDC density. WT GC B cells exhibited relatively stable MC distributions, with only modest, density-dependent trends, such as a slight increase in MC5 (linked to large displacement variability) and a significant decrease in MC0 (associated with minimal movement) (**Figure 3g**). Conversely, MT GC B cells displayed dynamic and pronounced shifts in MC composition, characterized by steeper slopes. Notably, MT GC B cells showed a significant depletion of MC7 and MC8 (clusters associated with high motility), as FDC density increased (**Figure 3h**). The steeper slopes in MT compared to WT implied that MT cells exhibited more frequent behavioral shifts in response to the dense FDC network, while WT GC B cell motility adapted in a regulated and constrained manner.

To understand GC B cell motility at fine spatial resolution, we quantified motility features at increasing physical distances from the DZ and dLZ. Although WT and MT GC B cells exhibited similar speeds at distances far from the DZ, in approaching the DZ, MT GC B cells increased their speed, while WT gradually slowed (**Figure 3i**). Conversely, MT GC B cells farthest from the dLZ exhibited significantly higher speeds than WT GC B cells and, as they approached the dLZ, their speeds slowed to match WT speeds, suggesting that the physical constraints of the dLZ limit MT cells’ ability to maintain higher speeds. WT GC B cells exhibited steady motility regardless of their distance from the dLZ (**Figure 3j**). In terms of directional persistence, MT GC B cells displayed increased tortuosity in the regions nearest to DZ (**Figure 3k**) and farthest from dLZ (**Figure 3l**), resulting in the largest difference compared to WT GC B cells near DZ. Morphological changes also showed divergent profiles between WT and MT GC B cells according to distance from DZ and dLZ (**Figure 3m-n**). MT GC B cells showed the greatest shape change near the DZ (**Figure 3m**), which decreased toward the dLZ, whereas WT GC B cells exhibited the opposite trend, showing greater shape change near the dLZ (**Figure 3n**), ultimately converging to similar morphodynamic profiles. These results further support that FDC-high density regions enable MT GC B cells to adopt WT-like behavior. Altogether, EZH2^Y641F^ mutation profoundly alters the FDC-dependent spatial regulation of GC B cell motility, contributing to the emergence of distinct motility behaviors within the GC.

### MT EZH2 GC B cells alter interzonal transitions that reduce recycling to DZ and increase dLZ localization

Given the spatially regulated motility patterns observed in the MT EZH2 GC B cells, we hypothesized that these patterns reflect altered mechanisms governing interzonal traversal across the GC microenvironment. Based on 5 zones (DZ, DZ-sLZ, sLZ, sLZ-dLZ, dLZ), we constructed a Markov chain diagram to visualize the likelihood of GC B cells transitioning from one zone to another, providing a comprehensive view of interzonal dynamics (**Figure 4a-b**). Notable findings from this analysis included decreases in the probability of MT GC B cells transitioning from DZ-sLZ to DZ (WT 37.9%, MT 25.5%) and from sLZ to DZ-sLZ (WT 9.6%, MT 5.1%) (**Figure 4a**), and an increased probability of transitioning from sLZ-dLZ to dLZ (WT 32.5%, MT 43.9%) (**Figure 4b**). To evaluate whether the altered interzonal transition dynamics observed in MT GC B cells translate into distinct localization patterns, we conducted a Monte Carlo simulation based on the WT and MT-specific Markov chain to generate a long-term equilibrium distribution of GC B cells across all zones (**Figure 4c**). Notable differences in zonal localization patterns were observed including: WT GC B cells were more localized in the imaged DZ (25.3%) than MT (18.8%), while MT GC B cells accumulated more strongly in the dLZ (15.1%) than WT (11.2%). These results demonstrate that the rewired transition landscape in MT GC B cells ultimately reshapes their spatial occupancy within the GC microenvironment.

**Figure 4.**
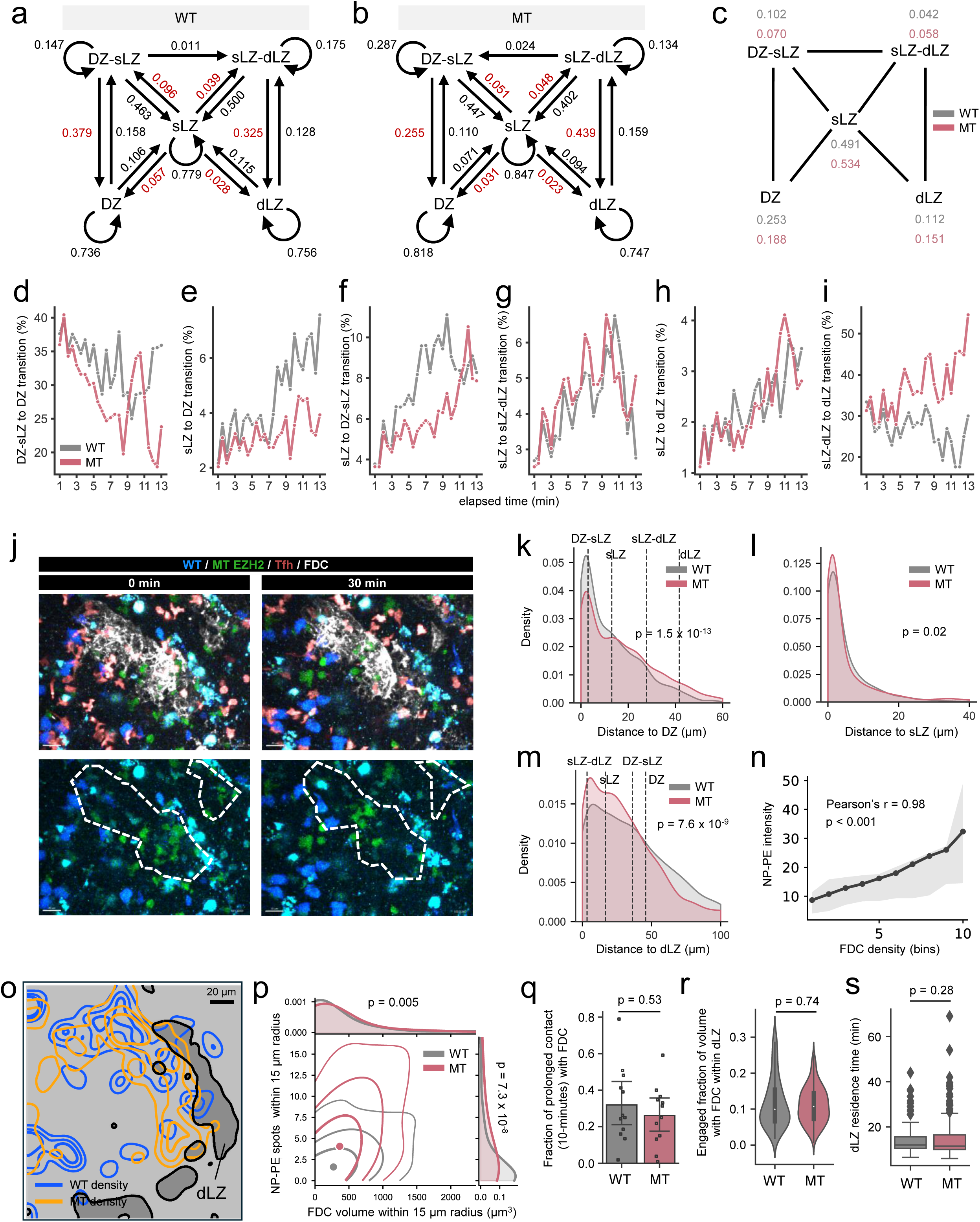
EZH2 mutation on GC B cells alters interzonal transition dynamics that lead to localization to the dLZ while separated from the DZ. **a-b.** Markov chain diagram depicting interzonal transitions occurring between two consecutive 8-minute tracks computed for each WT (**a**) and MT GC B cells (**b**) (WT n=483, MT n=793 cells). Red color highlights the transition of interest. **c.** Steady-state solution of Markov process based on the Markov chain diagram via Monte Carlo simulation with 10^6^ iterations. **d-i.** Zonal transitions computed at various elapsed times of track across all GCs: DZ-sLZ to DZ (**d**), sLZ to DZ (**e**), sLZ to DZ-sLZ (**f**), sLZ to sLZ-dLZ (**g**), sLZ to dLZ (**h**), and sLZ-dLZ to dLZ (**i**) transitions. **j.** Two-photon microscopy images showing localization of MT GC B cells within dense FDC networks (dotted white line). Top images show all four channels, and bottom images show only GC B cell channels (blue and green). Scale bars, 20 µm. **k-m.** GC B cell KDE distribution of average distance to DZ (**k**), sLZ (**l**), dLZ (**m**). Vertical dotted line depicts highest probability distance from the distance distribution for every other zone. Distance is calculated based on the average of cell trajectories to the boundaries of each zone. P values are from Mann-Whitney U test. **n.** Mean NP–PE intensity across ten FDC density quantile bins. For each movie and frame, all voxels were assigned to one of ten quantile bins according to continuous FDC density, and mean NP–PE intensity was calculated within each bin. The line indicates the median of the movie-level values, and shading indicates the 95% hierarchical bootstrap confidence interval obtained by resampling movies and then frames within movies over 5,000 iterations. Data comprises n = 3 movies and 295 observations per density bin. Pearson correlation calculated across the ten plotted bin medians. **o.** Smoothed spatial density contours of segmented WT (blue) and MT (orange) GC B cell positions overlaid on the dLZ mask (gray region). Density contours were generated by projecting the 3D centroids of B cells in XY and applying a Gaussian kernel with σ= 18. Contours represent the 75th, 90th and 97th percentiles of positive density-map values. Scale bars, 20 µm. **p.** Joint distribution of local FDC volume and NP–PE spot abundance surrounding 15 µm radius of individual WT and MT GC B cells at one midpoint frame. Filled circles indicate group means. P values on the marginal distributions are from Mann–Whitney U test (WT n = 122, MT n = 236 cells). **q.** Fraction of GC B cells persistently contacting FDCs for 10 min per GC (mean ± 95% C.I.). Each dot represents one GC. P value is from Mann-Whitney U test (including WT n = 12 and MT n = 12 GCs, and 161-561 tracks/GC). **r.** Average overlapping volume fraction of GC B cell with FDC in dLZ. Each point represents a cell. White dot represents median, boundaries of black box represent first and third quartile. P value is from Mann-Whitney U test (WT n = 120, MT n = 196 cells). **s.** Average dLZ residence time for GC B cells labeled as dLZ. P value is from Mann-Whitney U test (WT n = 120, MT n = 196 cells). Each point represents one cell. Line inside box represents median, boundaries of the box represent first and third quartile. Lines outside box represent whiskers.

To further assess this observation, we calculated the interzonal transition probability over all time intervals (**Figure 4d-i**). This analysis revealed that MT GC B cells exhibit reduced transitioning DZ-sLZ to DZ (**Figure 4d, Figure S7a**), sLZ to DZ (**Figure 4e, Figure S7b**) and sLZ to DZ-sLZ (**Figure 4f, Figure S7c**) compared to their WT counterparts. This suggests reduced DZ transition recycling of MT GC B cells, confirming previous observations^29^. We also found that MT GC B cells showed comparable sLZ to sLZ-dLZ (**Figure 4g, Figure S7d**) and sLZ to dLZ transitioning (**Figure 4h, Figure S7e**), but significantly higher sLZ-dLZ to dLZ transition probabilities compared to WT GC B cells across most time intervals (**Figure 4i, Figure S7f**).

The imaging data provided direct visual evidence that, as a result of altered interzonal transitions, MT GC B cells were more frequently situated within the dense FDC network compared to WT GC B cells throughout our observations (**Figure 4j, Supplementary Video 3**). To substantiate this, we quantified the spatial localization of GC B cells within each of the defined GC zones by calculating the distance from cells to the boundary of the zone. The results showed that WT GC B cells were more frequently localized near the DZ (**Figure 4k**), which aligns with their higher probability of recycling to this zone. In contrast, MT GC B cells showed increased sLZ localization (**Figure 4l**), with a significant difference also observed in the dLZ where MT GC B cells were more likely to be found compared to WT GC B cells (**Figure 4m**). Clearly, EZH2^Y641F^ mutation alters the behavioral programs promoting the accumulation of GC B cells within the dLZ, with fewer in the DZ, further supporting the premise that MT EZH2 GC B cells have increased dependency on the FDC network^29^.

We next asked whether this spatial bias toward the dLZ was associated with increased antigen access and was independent of initial B cell receptor (BCR) affinity. We adoptively transferred fluorescently labeled WT and MT B cells carrying the B1-8^hi^ background, which confers a shared high-affinity NP-reactive BCR. Following NP-OVA immunization, fluorescent NP–PE was administered and conducted intravital imaging 24 h later (**Figure S7g**). NP-PE spots were concentrated within the dLZ and enriched relative to the volume occupied by this region (**Figure S7h-i**). Consistently, NP-PE intensity increased with FDC density, identifying the dLZ as an antigen-enriched FDC niche (**Figure 4n**). Spatial density mapping further showed greater localization of MT than WT GC B cells within the dLZ despite their shared B1-8^hi^ background (**Figure 4o**). At the single-cell level, MT GC B cells were surrounded by greater local FDC volume and NP-PE spot abundance within a 15 µm radius than the WT (**Figure 4p**). Thus, the preferential dLZ localization of MT GC B cells persisted under a standardized initial BCR affinity and increased their exposure to FDC-associated antigen.

### FDC dependence of MT EZH2 GC B cells is not due to the intrinsic enhancement of FDC interactions

Observed increases in transitioning of MT EZH2 GC B cells toward dLZ may reflect altered quality and nature of interactions with FDCs. To investigate this, we quantified the fraction of GC B cells persistently contacting FDCs over a 10-minute duration. Surprisingly, our analysis revealed no significant difference in the overall persistence of FDC contacts between WT and MT GC B cells (**Figure 4q**). Furthermore, to probe the nature of FDC contacts, we focused specifically on the dLZ, where both GC B cell types make persistent contact within the confined region throughout the observations (**Figure S7j**). Within the dLZ, we quantified the degree to which GC B cells physically engaged with FDCs based on the overlapping volume fraction of GC B cells with FDCs, serving as an indicator of physical engagement during interactions. Our analysis revealed no statistical difference between WT and MT cells in the volume fraction engaged with FDC (**Figure 4r**) suggesting that MT GC B cells do not exhibit enhanced physical engagement with FDCs. This result was consistent with GC B cells that contact over 10-minutes within sLZ, showing comparable interaction surface size with FDC (**Figure S7k**). We also evaluated the residence time of GC B cells within dLZ to identify how long individual cells remain within dLZ or the DZ and found no statistical difference (**Figure 4s**, **Figure S7l**). We next asked whether the relative abundance of MT GC B cells within individual GCs influenced their engagement with FDCs. MT cell abundance was not significantly correlated with either closer positioning to FDCs (**Figure S7m**) or longer FDC contacts by MT cells relative to WT cells (**Figure S7n**). Together, these results highlight the important distinction that, while the EZH2^Y641F^ mutation alters the migration patterns and spatial distribution of GC B cells favorable to dLZ, it does not significantly enhance their physical interactions with FDCs within the dLZ.

### MT EZH2 GC B cells exhibit decreased quality of interaction with Tfh cells

Following the quantification of FDC interactions, we investigated whether the EZH2^Y641F^ mutation altered the interaction dynamics between GC B cells and Tfh cells, which could affect their selection and survival. We have previously shown that MT EZH2 GC B cells exhibit reduced dependence on Tfh cell-derived signals, impaired affinity maturation, and restricted clonal diversity during the GC reaction^13^. However, it remains unknown whether these deficits in Tfh dependence and affinity maturation inferred from molecular readouts are reflected in the cell-level behavioral traits of GC B– Tfh cell interactions, or whether MT GC B cells form qualitatively distinct or biophysically weak immune interactions with Tfh cells. We found that WT and MT EZH2 GC B cells have similar distance patterns from Tfh cells (**Figure 5a**), indicating non-significant differences in the overall normalized number of Tfh contacts for both WT and MT GC B cells (**Figure 5b**). However, our imaging data showed that WT GC B cells engage in longer or more persistent interactions with Tfh cells compared to MT GC B cells (**Figure 5c, Supplementary Videos 4-5**). To explore the quality of these interactions, we quantified the fraction of GC B cells undergoing prolonged contact with Tfh cells for continuous 10-minute observations. We found a significantly higher prolonged-contact fraction in WT than MT GC B cells (**Figure 5d**). Additionally, we analyzed the fraction of volume overlap between GC B cells and Tfh cells to quantify engagement size during these persistent interactions, finding that WT GC B cells exhibited significantly larger engagement volume with Tfh cells compared to MT GC B cells (**Figure 5e, Figure S8a**).

**Figure 5.**
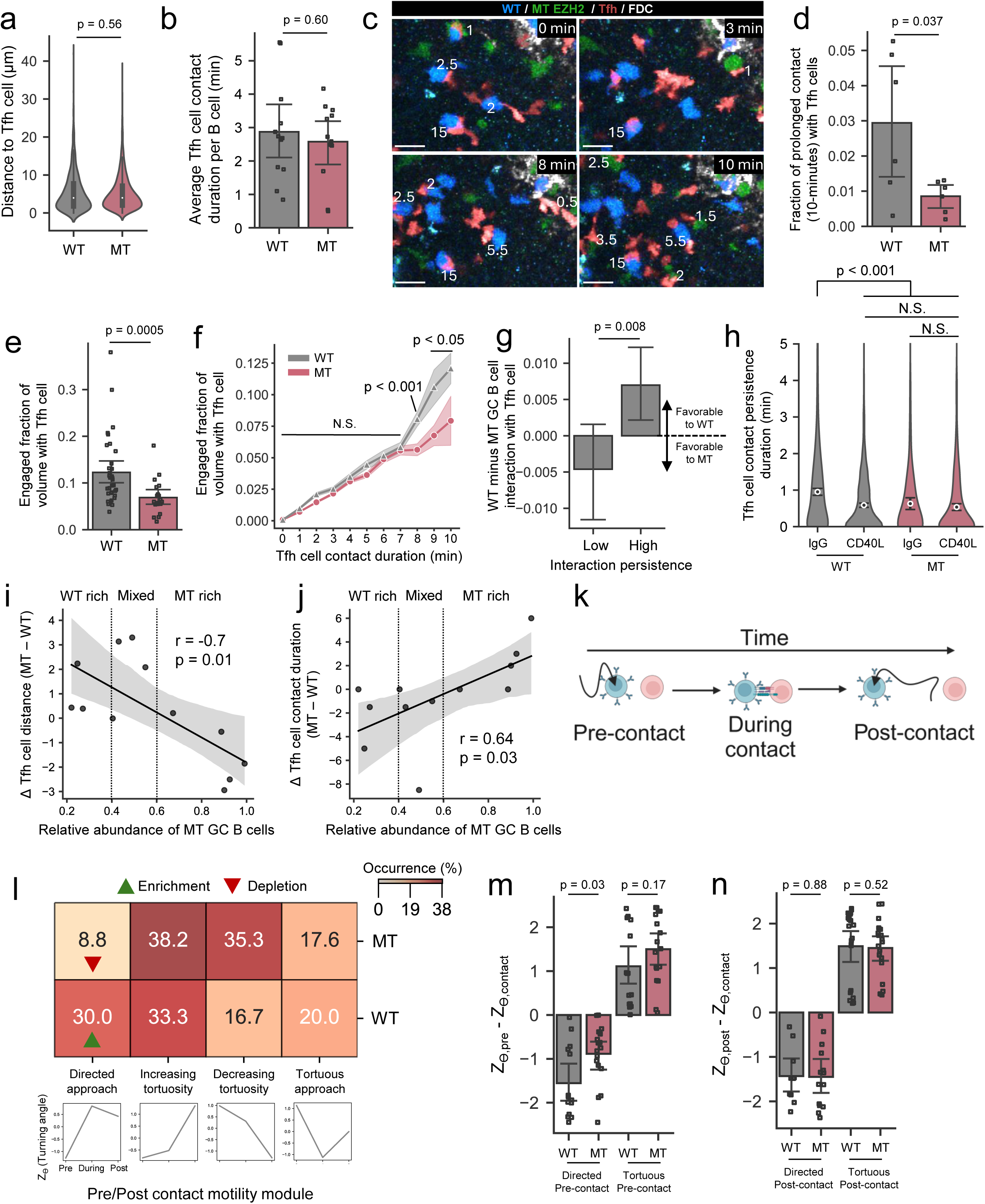
MT EZH2 GC B cells display impaired high-quality contact with Tfh and disrupted contact-coupled motility responses. **a.** Average distance to closest Tfh cell. Each point represents 10-minute track of cell. White dot represents median, boundaries of black box represent first and third quartile. P value is from Mann-Whitney U test (including WT n = 1,482 and MT n = 2,587 10-minute track of cells). **b.** Total number of Tfh cell contact divided by the total number of GC B cells per GC (mean ± 95% C.I.). Each dot represents one GC. P value is from two-sided unpaired t-test (WT n = 12, MT n = 12 GCs). **c.** Two-photon microscopy images showing longer contact of WT with Tfh cell compared to that of MT. The numbers indicate contact durations (unit: minutes). Scale bars, 20 µm. **d.** Fraction of GC B cells persistently contacting with any Tfh cell sequentially for 10 min per GC (mean ± 95% C.I.). Each dot represents one GC. P value is from two-sided unpaired t-test (WT n = 6, MT n = 6 GCs). **e.** Average overlapping volume fraction of GC B cell with Tfh cells that maintain high persistent interactions for 10 minutes (mean ± 95% C.I.). Each dot represents a cell. P value is from Mann-Whitney U test (WT n = 34, MT n = 20 cells). **f.** Average overlapping volume fraction of GC B cells with Tfh cells according to contact duration (mean ± 95% C.I.). P values are from Mann-Whitney U test (mean ± sem). (WT n = 35 - 349 tracks, MT n = 31 - 616 tracks). **g.** Difference of interaction fraction between WT and MT GC B cells with Tfh cells for every GC across each interaction duration (mean ± 95% C.I.). Interaction events were stratified into low interaction duration (0–7 minutes) and high interaction duration (7.5–10 minutes). P value is from Mann-Whitney U test (mean ± 95% C.I., Low n = 132, High n = 72 GCs per Tfh cell interaction duration. **h.** Duration of persistent Tfh cell contact for IgG control and CD40L inhibition (mean ± 95% C.I.). Each point represents a cell. P values are from Kruskal-Wallis test followed by Dunn’s post-hoc test (WT IgG n = 619, WT CD40L n = 950, MT IgG n = 201, MT CD40L n = 341 cells). **i-j.** Difference in median distance from each GC B cell to its nearest Tfh cell (**i**) and Tfh cell contact duration (**j**) between MT and WT GC B cells within each GC against the relative abundance of MT GC B cells. r indicates Pearson correlation coefficient with corresponding p value (12 GCs). Each dot represents a GC. The line indicates the linear regression fit, and shading denotes the 95% confidence interval. Vertical dotted lines indicate the thresholds used to define WT rich, mixed and MT rich GCs. **k.** Schematic illustration depicting the measurement of emergent motility changes of GC B cell pre-contact, during contact, and post-contact with Tfh cell. **l.** Fractional occurrence of four pre/post contact motility modules identified by time-series K-means clustering of average turning angle based on GC B cells that interact with Tfh cell more than 7 minutes. (WT n = 30, MT n = 34 cells). Bottom, time-series cluster barycenter shows the representative fluctuation of turning angle for each module. Z_Ө_ represents the average Z score of turning angle. Enrichment and depletion represent statistical over-representation and under-representation compared to random sampling of GC B cells, respectively. **m.** Difference in Z score of turning angle between pre-contact and during contact with Tfh cell (mean ± 95% C.I.). Each dot represents a cell. P value is from Mann-Whitney U test (WT Directed Precontact n = 14, MT Directed Precontact n = 17, WT Tortuous Precontact n = 16, MT Tortuous Precontact n = 17 cells). **n.** Difference in Z score of turning angle between post-contact and during contact (mean ± 95% C.I.). P value is from Mann-Whitney U test (WT Directed Postcontact n = 10, MT Directed Postcontact n = 14, WT Tortuous Postcontact n = 20, MT Tortuous Postcontact n = 20 cells).

We next examined how contact duration influences engagement size between GC B and Tfh cells. For short-lived interactions (less than 7 minutes), engagement sizes were comparable between WT and MT GC B cells. However, for prolonged interactions (more than 7 minutes), WT GC B cells exhibited significantly larger engagement areas than MT cells, suggesting that the 7-minute mark represents a threshold beyond which qualitative differences in interaction become evident (**Figure 5f**). Stratifying contacts by this 7-minute threshold revealed that MT GC B cells were more inclined toward low-persistence interactions and WT toward high-persistence interactions (**Figure 5g, Figure S8b, Materials and Methods**). This implies that the dominance of low affinity MT clones previously described is mirrored at the cell behavioral level by their inability to form stable, high-quality Tfh cell interactions^13^. We also found that inhibition of CD40L significantly reduced the Tfh cell contact persistence duration of WT GC B cells to levels comparable to MT, whereas MT GC B cells were unaffected by CD40L blockade, which was consistent with their already reduced interaction quality and confirmed our previous observations^13^ (**Figure 5h**). We next asked whether Tfh cell interactions varied with the relative abundance of MT GC B cells within individual GCs. At low MT cell abundance, MT cells were positioned farther from Tfh cells (**Figure 5i**) and maintained shorter contacts than WT cells (**Figure 5j**), however, these relationships reversed as MT cell abundance increased. In summary, the shift toward lower-quality interactions in MT GC B cells suggests that while MT GC B cells encounter Tfh cells more frequently due to their enhanced motility, these encounters do not always result in the formation of robust or high-quality contacts due to reduced affinity maturation, although Tfh cell engagement was increased in MT rich GCs.

### WT GC B cells and Tfh cells dynamically adjust their motility to facilitate subsequent interactions

Given the compromised ability of MT EZH2 GC B cells to establish high-quality contact with Tfh cells, we sought to determine whether the difference between WT and MT arose only at the moment of Tfh cell contact, reflecting the previously described lower affinity of MT cells, or whether MT cells already exhibit distinct behavioral program before encountering Tfh cells. Specifically, we asked whether GC B cells exhibit motility signatures that are modulated around the time of Tfh cell interaction. To examine these dynamics, we analyzed the emergent motility patterns of WT and MT GC B cells immediately preceding, during, and following Tfh interactions (**Figure 5k**). Time-series k-means clustering of turning angles revealed four pre/post contact motility modules, namely: direct approach, increasing tortuosity, decreasing tortuosity, and tortuous approach (**Figure S8c-d**). We found that the “direct approach” module, which depicted high progressive motion of GC B cells before interacting with Tfh cells (compared to during and post interactions), was highly enriched in WT and depleted in MT (**Figure 5l**). Quantification of turning angle Z score differences between pre-contact and during contact showed more directed movement before the contact with Tfh and WT, compared to MT (**Figure 5m**). The difference in the motility post-Tfh interaction was comparable between WT and MT (**Figure 5n**). This suggests that WT GC B cells are capable of purposeful movement toward Tfh cells prior to interaction, allowing them to efficiently close the distance and initiate interactions, whereas MT GC B cells do not appear to adapt their responses prior to the contact with Tfh cells. In addition to the turning angle, we examined speed and volume by time-series clustering. While there was no significant difference in speed (**Figure S8e-g**), MT GC B cells were enriched for an ‘Expansion during’ module (increase of cell volume during interaction), suggesting interaction-dependent size adjustment (**Figure S8h**).

We next analyzed Tfh cell motility before, during and after GC B cell interactions. Time-series clustering of instantaneous Tfh speed revealed a ‘fast approach–fast escape’ pattern (high speeds both before and after contact) enriched with WT but not MT GC B cells, suggesting that Tfh cells engagement dynamics depend on the B cell partner (**Figure S8i**). In contrast, turning angle analysis showed no significant differences between WT and MT (**Figure S8j**), suggesting that speed modulation, rather than directional control, serves as a dominant strategy for Tfh cells in efficiently locating and transiently engaging with WT GC B cells. Collectively, these results highlight that pre-contact motility adjustments are coordinated for interaction between WT GC B cells and Tfh cells, but MT EZH2 GC B cells exhibit impaired pre-contact directional motility as well as volume expansion during Tfh interactions. This implies that WT GC B cells and Tfh cells interactively provide cues that elicit efficient recognition and scanning, whereas MT cells fail to trigger such adaptive Tfh responses, a previously unrecognized mechanism by which oncogenic EZH2 mutation undermines T cell-dependent selection.

### Tri-interaction analysis reveals FDC-dependent priming of Tfh engagement and altered post Tfh contact zonal trafficking in MT EZH2 GC B cells

After quantifying the differential interaction dynamics of GC B cells with FDCs and Tfh cells, we sought to determine whether the relationship between GC B cell–FDC and GC B cell–Tfh interactions was altered by the EZH2^Y641F^ mutation. To investigate this, we analyzed only WT and MT GC B cells that engaged in persistent contact with both FDC and Tfh and computed the time difference between the initiation of GC B cell–FDC and GC B cell–Tfh persistent contacts. WT GC B cells exhibited no significant bias in order of interacting with FDC or Tfh, with some cells persistently contacting FDCs before interacting with Tfh cells, while others initiating interactions in the reverse order (**Figure 6a**). A significant portion of WT cells displayed marginal delay, suggesting simultaneous interactions with FDCs and Tfh cells. In contrast, MT GC B cells demonstrated a distinct bias toward persistent FDC contact occurring before Tfh interactions (**Figure 6a**).

**Figure 6.**
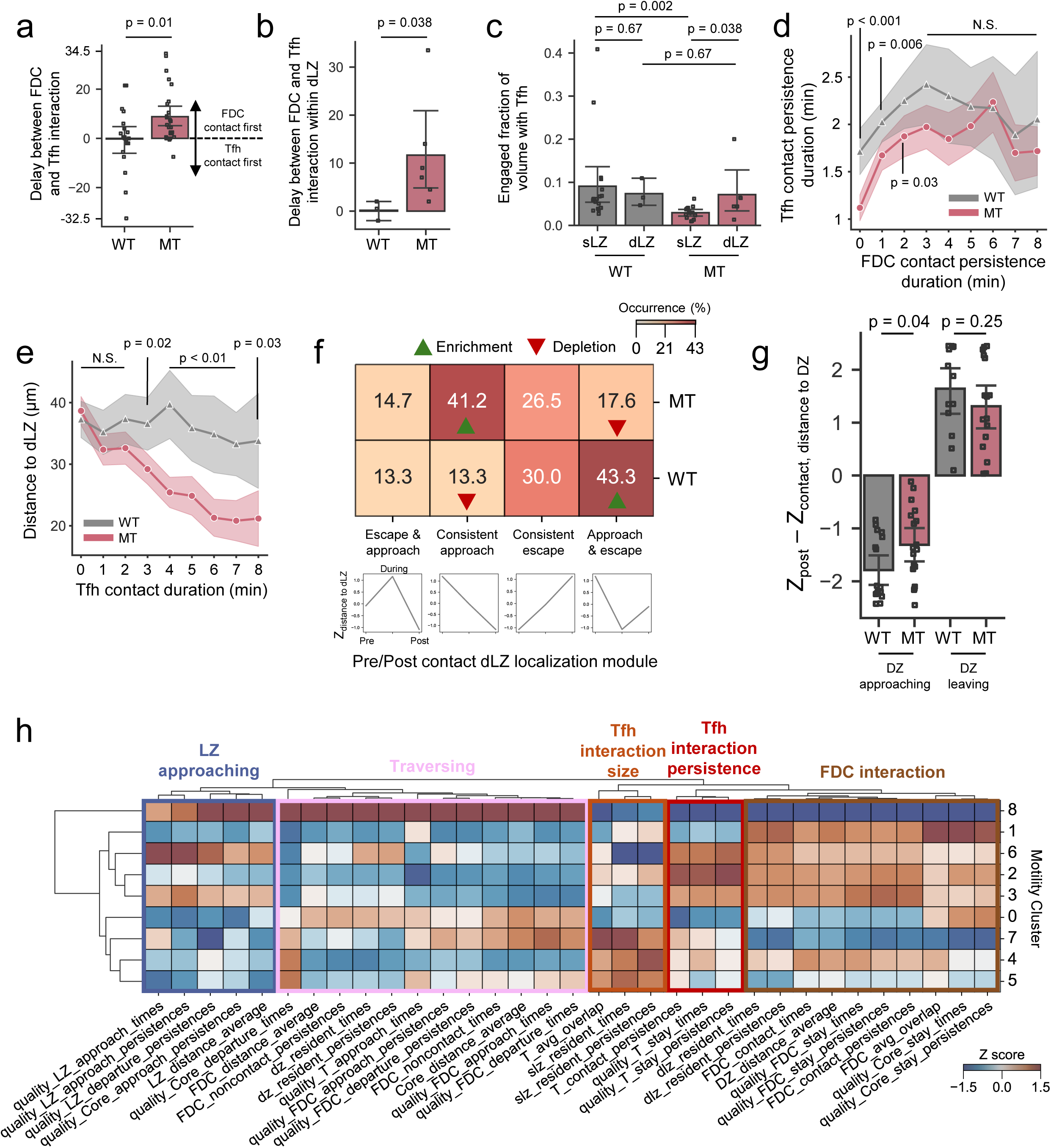
Tri-interaction analysis between GC B cells, Tfh, and FDCs reveals FDC-dependent priming of Tfh engagement and altered post Tfh contact zonal trafficking in MT EZH2 GC B cells. **a.** Time difference between initial persistent contact (more than 7 minutes with both interactions) time points with FDC and Tfh (mean ± 95% C.I.). Each dot represents a cell. P value is from Mann-Whitney U test (WT n = 20, MT n = 26 cells). **b.** Time difference between initial persistent contact (more than 7 minutes with both interactions) time points with FDC and Tfh restricted to dLZ GC B cells (mean ± 95% C.I.). Each dot represents a cell. P value is from Mann-Whitney U test (WT n = 3, MT n = 6 cells). **c.** Average overlapped volume between zone-specific GC B cells that maintain high persistent interactions (more than 7 minutes) with Tfh cells (mean ± 95% C.I.). Each dot represents a cell. P value is from Kruskal-Wallis test followed by Dunn’s post-hoc test (WT sLZ n = 18, WT dLZ n = 3, MT sLZ n = 14, MT dLZ n = 6 cells). **d.** Quantification of Tfh contact persistence duration at each FDC contact persistence duration (mean ± 95% C.I.). (WT n = 35-258, MT n = 88 - 349 cells). **e.** Quantification of distance to dLZ at each Tfh contact duration (mean ± 95% C.I.). (WT n = 57-349, MT n = 69-616 cells). **f.** Fraction occurrence of four pre/post contact dLZ localization module identified by time-series K-means clustering of dLZ distance. Bottom, time-series cluster barycenter shows the representative fluctuation of dLZ distance for each module. Z_distance to dLZ_ represents the average Z score of dLZ distance. Enrichment and depletion represent statistical over-representation and under-representation compared to random sampling of GC B cells, respectively. **g.** Difference in Z score of distance to DZ between post-contact and during contact with Tfh cell (mean ± 95% C.I.). Each dot represents a cell. P value is from Mann-Whitney U test (WT DZ approaching n = 17, MT DZ approaching n = 18, WT DZ escaping n = 13, MT DZ escaping n = 16 cells). **h.** Average Z score of selected colocalization features based on the statistical significance between WT and MT by Mann-Whitney U test. Hierarchical clustering is based on the Euclidean distance using the Ward method.

To determine whether this observed bias of MT GC B cells was linked to their localization within the dLZ, we tracked persistent B–Tfh cell interactions specifically in this region. WT GC B cells maintained simultaneous interactions with both FDCs and Tfh cells within the dLZ, consistent with their broader interaction patterns (**Figure 6b**). However, MT GC B cells consistently showed interactions with FDC prior to Tfh engagement, reinforcing the premise that MT GC B cells require FDC contact as a precursor to Tfh interactions. To understand this, we quantified the GC B cell’s engaged volume fractions with Tfh cells that engage in long persistent contact with Tfh cell that occurred within the sLZ and dLZ. In contrast to WT GC B cells, which showed no significant difference in engaged volume with Tfh cells between sLZ and dLZ, MT GC B cells exhibited lower engaged volume with Tfh cells in the sLZ, but increased engaged volume in the dLZ, reaching WT level (**Figure 6c**). In MT cells, both engagement size and interaction persistence with Tfh cells increased with greater FDC contact persistence, reaching levels comparable to WT (**Figure 6d**). We also confirmed that MT GC B cells tend to form longer Tfh interactions in close proximity to dLZ, whereas the duration of Tfh interaction was independent of proximity to dLZ for WT GC B cells (**Figure 6e**). This was characterized by lower fractions of MT sLZ B cells (**Figure S9a**), and a comparable fraction of MT dLZ B cells that make more than a 7-minute Tfh cell contact, compared to WT (**Figure S9b**). This provides further evidence that MT GC B cells rely on their interaction within the FDC-rich dLZ to facilitate Tfh engagement, while WT cells display no dependence on dLZ for Tfh interaction.

To examine how Tfh contact influences the subsequent behavior or fate of GC B cells, we applied the same pre/post-contact module identification strategy, identifying four distinct modules based on cell distance to the dLZ before, during, and after Tfh interaction lasting more than 7-minutes. Among these, WT GC B cells were significantly enriched in the “approach-and-escape” module, where cells approached the dLZ prior to Tfh interaction but subsequently migrated away from the dLZ post-contact (**Figure 6f**). This directional movement was further supported by an increased frequency of WT GC B cells approaching the DZ following Tfh interaction (**Figure 6g**), a pattern consistent with canonical DZ recycling during selection. In contrast, MT GC B cells were depleted in this module and instead enriched in the “consistent approach” module, showing sustained localization within the dense FDC niche even after Tfh engagement (**Figure 6f**). This difference diminished in comparison of the pre/post B cell behavior based on short duration Tfh interaction (2 to 7 minutes) characterized by a comparable approach-and-escape module (**Figure S9c**). The contact surface size did not significantly affect the fraction of WT and MT B cells following this module (**Figure S9d**), suggesting that contact duration is a greater determinant of B cell fate than contact size. These findings suggest that GC B cells undergo DZ recycling after prolonged interactions with Tfh cells in the LZ, and that EZH2 mutation impairs this process.

To establish whether motility itself encoded such functional interactions, we examined the spatial and interaction properties (colocalization features) of distinct MCs (**Figure 6h**). Notably, MC8 was enriched for cells traversing GC zones, suggesting a role in inter-zonal trafficking. MC1–3, which exhibit slower movement, were associated with higher levels of FDC interaction. Conspicuously, MC2 showed the strongest enrichment for Tfh interaction persistence, which is associated with interaction time, while MC7 was distinguished by the largest Tfh overlapping volume, associated with interaction size. The distribution of MC was altered following inhibition of CD40L, which targets Tfh cell interaction, although to a lesser extent in MT relative to WT, decreasing the MC2 fraction and enriching towards MC5-8 (**Figure S9e**). These results support the view that motility patterns are functionally informative and tightly linked to interaction strategies within the GC microenvironment.

### scMOTIPh maps GC B cell behaviors to underlying transcriptomic programs

To elucidate the molecular mechanisms underlying distinct GC B cell behaviors, we expanded our behavioral assessments to incorporate both 123 motility dynamics (**Supplementary Table 1**) and 135 GC microenvironmental colocalization features (**Supplementary Table 2**), specifically analyzing interactions with Tfh cells and FDCs. Unsupervised clustering within this integrated behavior space identified eight behavior clusters (BCs) (**Figure 7a**, **Materials and Methods**), which captured distinct motility patterns and reflected specific GC microenvironmental associations (**Figure 7b, Figure S10a**). For example, BC0 represented the slowest cells enriched in DZ, BC1 comprised relatively slow sLZ cells, BC2 corresponded to dLZ cells engaging with FDCs, BC3 was enriched for cells with strong Tfh interactions, and BC7 captured the fastest migrating cells. BCs were unevenly distributed between WT and MT. For example, BC3, which is associated with robust Tfh engagement, was enriched in WT, consistent with the observed Tfh interaction deficits in MT (**Figure 7c**). Furthermore, BCs mapped explicitly to GC zones, for example, BC0 and BC6 consisted of cells in DZ with low and high motility, respectively, while BC2 and BC4 were associated with dLZ and LZ–dLZ, respectively (**Figure S10b**). In parallel, we profiled GC B cells using single-cell RNA-seq^13^, identifying 10 transcriptional clusters via Leiden clustering (**Figure S11a**). Leveraging curated gene signatures for centroblasts, centrocytes, memory precursors, and recycling B cells (**Figure S11b**), cluster-specific differentially expressed genes (**Figure S11c**), and community detection with literature-derived subtype gene sets^36^ (**Figure S11d**), we annotated clusters as DZ, DZ-LZ, LZ, and pre-memory GC B cells (**Figure S11e**). Because our imaging data focused on intra-GC behaviors, pre-memory cells were excluded from downstream mapping.

**Figure 7.**
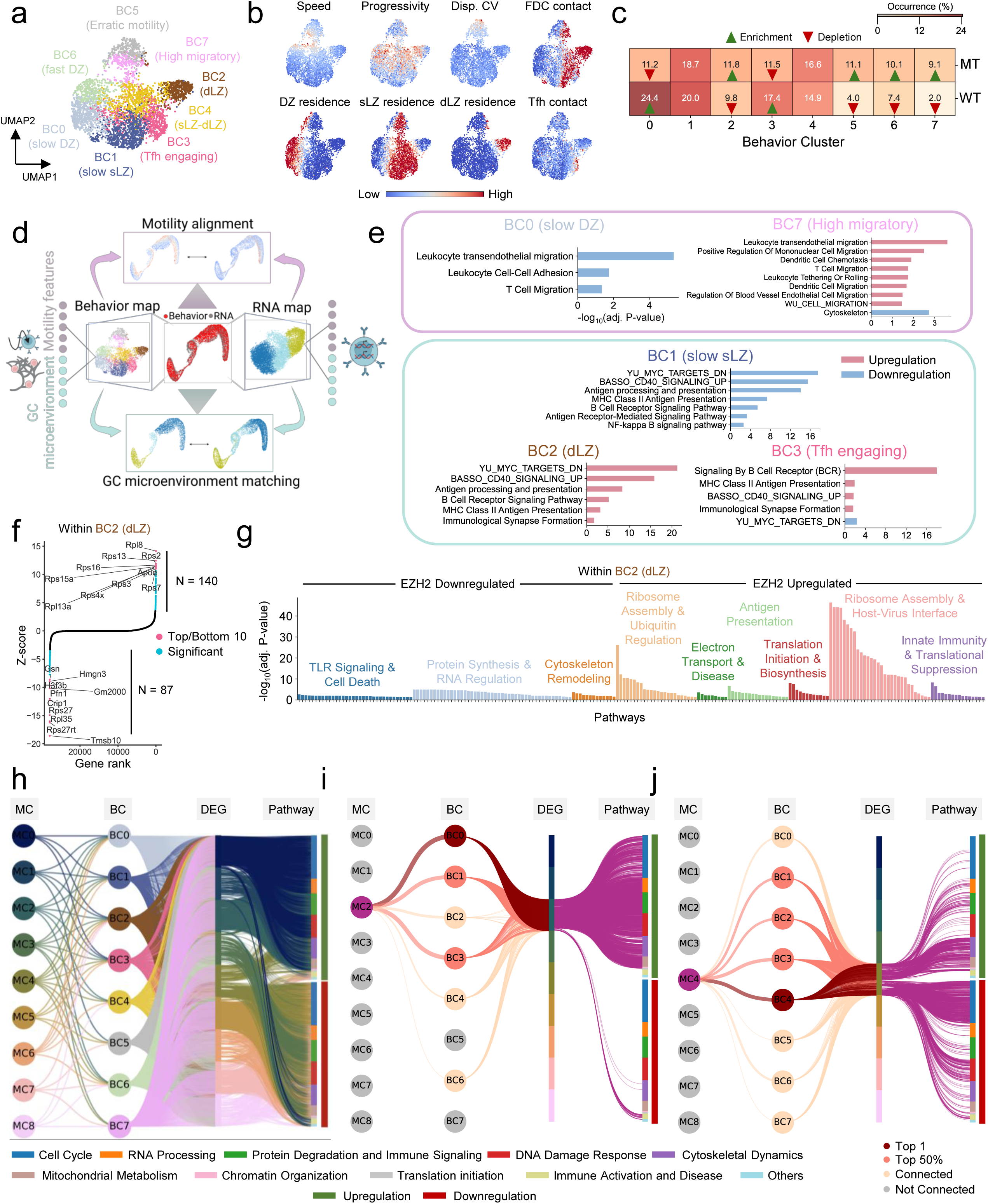
Integrated GC microenvironmental behavior-transcriptome analysis reveals dLZ signatures and MC-specific pathways. **a.** UMAP representation of unsupervised k-means clustering identifies eight unique BCs (n = 4,069 10-minute track of cells). **b.** Interpretable motility features and colocalization features projected onto the UMAP space. **c.** Fractional occurrence of eight BCs across WT and MT GC B cells (WT n = 1,482 and MT n = 2,587 10-minute track of cells). Enrichment and depletion represent statistical over-representation and under-representation compared to random sampling of GC B cells, respectively. **d.** Schematic illustration of scMOTIPh using Biorender. **e.** Validation of behavior-transcriptome integration using BC-specific GSEA. Top and bottom panels represent validation of motility and GC microenvironment association, respectively. **f.** Z score gene ranking based on MT EZH2 vs WT within BC2 (dLZ). **g.** Pathway analysis in BC2 (dLZ). Colors represent group of pathways based on Louvain community detection. **h.** Multipartite network visualization linking MCs, BCs, MC-specific differentially expressed genes (DEGs), and pathway modules. Pathway modules are defined based on Louvain community detection. Edges from MCs to BCs represent a fraction of MCs contributing to each BC, BC to DEGs represent shared gene signatures between BC and MC, DEGs to pathways are connected if the gene is a member of the corresponding pathway gene set. **i-j.** MC-specific multipartite network highlighting behavior clusters, DEGs, and enriched pathways selectively associated with MC2 (**i**) and MC4 (**j**).

We designed and applied a custom workflow named scMOTIPh (**s**ingle-**c**ell **M**ultiscale **O**ptimal **T**ransport for **I**mputing **Ph**enotypes), in which both behavior and transcriptomic spaces were defined at single-cell resolution. scMOTIPh is a custom computational framework for aligning cell-level behavior features with molecular-level profiles of single cells. Optimal transport finds the minimum-cost coupling between two distributions that map each point in one space (behavior) to points in the other space (RNA), thereby aligning cells across two different modalities^37–39^. As a result, locally similar cells are matched between two spaces, while incorporating two biologically meaningful axes: (1) a motility axis, matching continuous motility behaviors quantified in parallel with expression scores of curated motility-related genes (**Supplementary Table 6**), and (2) a GC zonal axis, aligning cells across DZ, DZ-LZ, and LZ compartments in both modalities (**Figure 7d**). The resulting alignment (**Figure S12a**) allowed us to directly assign molecular meaning to BCs using downstream gene expression analysis based on a nearest neighbor imputation scheme.

We validated the semantic integrity of the mapping by performing GSEA on differentially expressed genes for each BC. In the motility dimension, BC7, characterized by high migratory behaviors, showed strong enrichment for pathways including transendothelial migration, mononuclear cell migration, T cell and dendritic cell chemotaxis, and leukocyte rolling (**Figure 7e**, top panel; **Supplementary Table 7**), providing confidence in the single-cell mapping based on gene signatures (**Figure S12b**). In contrast, BC0 showed a marked downregulation of these migration-associated programs. Interestingly, cytoskeleton-related pathways were downregulated in BC7, possibly reflecting a more deformable migration behavior. In the microenvironmental context, BC2 and BC3, which are associated with FDC and Tfh engagement, respectively, were enriched for antigen processing and presentation, MHC class II pathways, BCR signaling, and immunological synapse formation (**Figure 7e**, bottom panel; **Supplementary Table 7**), all representing tightly interconnected pathways (**Figure S12b**). In contrast, BC1, although LZ-associated, was characterized by downregulation of these interaction-related signatures, as expected. Notably, BC3 showed canonical downregulation of MYC downregulated genes, consistent with recycling into the DZ, whereas BC2 showed upregulation of these genes, suggesting that dLZ-associated FDC-interacting cells may fail to recycle, which aligns with prior observations that MT GC B cells favor dLZ localization and display defective recycling dynamics^13^.

### Transcriptomic profiling of GC B cell behavioral states reveals dLZ and MC-specific gene programs

Building on our scMOTIPh mapping, we investigated how transcriptomic programs diverged within dLZ-associated BC2. We identified dLZ populations in the transcriptomic space and conducted differential expression analyses comparing MT EZH2 and WT GC B cells within dLZ, which revealed 140 upregulated and 87 downregulated genes (**Figure 7f, Supplementary Table 8**). This result contrasts with the canonical role of EZH2 as a gain-of-function repressor, for which we observed more downregulated than upregulated genes in bulk-RNA seq (**Figure S4a**). Conversely, MT GC B cells in dLZ showed substantial transcriptional upregulation. Ribosomal proteins such as Rpl8, Rpl13a, and Rps2 were prominently upregulated, suggesting enhanced translational output. Apoe was also upregulated, implicating altered lipid processing. For example, B cells are known to present lipid antigens captured by Apoe to Natural Killer T cells leading to restrained B cell differentiation and limited affinity maturation^40,41^. Conversely, downregulated genes included Gelsolin (Gsn), Profilin 1 (Pfn1), and Thymosin Beta 10 (Tmsb10), suggesting altered cytoskeletal remodeling.

To clarify the functional role of dLZ in the context of EZH2 mutation, we applied GSEA between MT EZH2 and WT based on dLZ-associated differentially expressed genes, revealing 9 pathway modules (**Figure 7g, Supplementary Table 9**). Among these, the *Toll-like receptor (TLR) signaling & cell death* module was significantly downregulated together with pathways related to apoptosis and programmed cell death, suggesting that the EZH2 mutation epigenetically silences death signaling in dLZ to enhance survival. Furthermore, the downregulation of TLR signaling suggests impairment of normal B cell functions, including B cell differentiation, survival, and antibody responses^42–44^. In addition, we observed downregulation of ‘cytoskeleton remodeling,’ with potentially significant implications on the biomechanics of immune synapses and abilities for discriminating antigen affinities^45^. The ‘antigen presentation’ module was upregulated in MT dLZ GC B cells, comprising pathways such as antigen presentation and MHC class II binding. These findings suggest that although Tfh engagement is generally reduced in MT GC B cells, key antigen presentation mechanisms remain active in dLZ. Upregulation of ‘ribosome assembly,’ ‘electron transport,’ and ‘translation initiation’ modules further suggest a hypermetabolic, biosynthetically active state, pointing to increased growth signals that support MT GC B cells in dLZ.

To understand how such transcriptional states relate to migratory behavior, we investigated the transcription-based functional states of MCs. Differentially expressed MC-specific genes were defined based on distinctiveness score (**Materials and Methods**), resulting in 300 gene signatures for each MC (**Figure S13a**). Using transcriptomic-based functional analysis of MCs, we mapped motility clusters (MCs) to behavior clusters (BCs), together with their differentially expressed genes and associated functional modules derived from related pathway groups (**Figure 7h**). Our analysis revealed that each MC was associated with distinct functional modules and showed characteristic patterns of up- or downregulation. MC2, representing a relatively low-motility, WT-enriched state (**Figure 1c**, **Figure 2g**), was most prevalent in BC0 (DZ GC B cells) and was strongly influenced by BC3, which represents Tfh-engaging cells (**Figure 7i**). This aligns with its strong association with upregulation of the cell cycle module and related pathways, including ‘cell cycle G1/S phase transition’, ‘mitotic cytokinesis’, and ‘Cyclin D associated events in G1’, consistent with a DZ-associated proliferative state (**Supplementary Table 10**). We also observed upregulation of pathways related to ‘protein degradation and immune signaling’ such as ‘NF-kappaB activation in B cells,’ ‘antigen processing,’ and ‘downstream B cell receptor signaling,’ likely reflecting contributions from BC3 cells with high Tfh interaction (**Supplementary Table 10**).

Other functional modules like RNA processing, DNA damage response, cytoskeleton dynamics, and mitochondrial metabolism were upregulated, whereas the immune activation and diseases module was downregulated, emphasizing distinct transcription-based functional roles in MC2. In contrast, MC4, a medium-motility state enriched in MT dLZ-associated cells (**Figure 1c**, **Figure 2g**), was heterogeneously influenced by all BCs but most abundant in BC4, which corresponds to the sLZ-dLZ region (**Figure 7j**). MC4 showed downregulation of cell cycle-related pathways, together with partial upregulation of protein degradation and immune signaling modules. The latter included B cell receptor signaling and beta-catenin-independent WNT signaling, which is known to promote GC B cell survival through Wnt5a secreted by FDCs^46^ (**Supplementary Table 10**). Conversely, partial downregulation within this module involved programmed cell death pathways, suggesting activation of survival mechanisms. MC4 also exhibited upregulation of cytoskeletal dynamics and mitochondrial metabolism and downregulation of DNA damage response, showing both overlap with and distinctions from MC2.

Across other functional modules – including metabolism, immune activation, and DNA damage response, MCs diverged, underscoring the specialization of transcriptional programs embedded within GC motility patterns (**Figure S13b–h**). Collectively, these results demonstrate that GC B cell motility behaviors encode distinct transcriptional states reflecting proliferation, selection, survival functions within the GC architecture.

## DISCUSSION

In this study we combine intravital two-photon imaging with single-cell analysis to track GC B cell behavior during maturation and investigated how it is affected by a lymphoma-associated point mutation in an epigenetic modifier. Although many studies have examined GC-derived lymphomas, the mechanisms by which B cells alter motility and cell-cell interactions to enable lymphomagenesis remain unclear^47,48^. Prior efforts to profile B-cell dynamics largely summarized B cell populations with a focus on few features (e.g., average speed or average turning angle)^49^. While informative, these approaches do not capture the higher-order trajectory structure, contact geometry, and cell-to-cell variability, all of which contribute to driving cellular heterogeneity^20^.

To address the need for a more highly-resolved understanding, we establish a robust cellular framework for measuring how EZH2 mutation alters the coordination between movement and microenvironmental interactions within intact GCs. Our approach involved defining multivariate behavioral states, quantifying their interactions at single-cell resolution, and linking behaviors to specific transcriptomic programs. This approach enabled the identification and classification of well-defined behavioral subtypes that are linked to specific aspects of GC function, such as immune synapse formation and intrazonal trafficking within the LZ.

A central outcome of this work is a mechanistic interpretation of the reduced Tfh cell reliance and increased FDC dependence of MT EZH2 GC B cells. We previously demonstrated that MT EZH2 GC B cells exhibit impaired interactions with Tfh cells and increased dependency on FDC signaling, supported by their upregulation of genes involved in FDC interaction, such as *Tnfrsf13c* (BAFF receptor), *Tnf* (Tumor Necrosis Factor), *Ltb* (Lymphotoxin Beta), and integrins including *Itgb2* (Integrin Subunit Beta 2) and *Itgb4* (Integrin Subunit Beta 4)^13^. Consistently, we visually observed that MT EZH2 GC B cells form less stable and lower-quality contacts with Tfh cells.

These weak interactions likely stem in part from the lower affinity maturation and reduced sensitivity to CD40L-dependent Tfh cell signals previously described for MT cells, but our results reveal an additional mechanism: MT cells fail to execute the pre-contact motility that WT GC B cells use to initiate productive Tfh synapses, both through the WT B cell’s increase in directional motion and corresponding increase in Tfh cell speed before engagement. This observation is consistent with recent findings showing that high-affinity B cells upregulate CD40-induced CCL17/CCL22 to attract CCR4-expressing Tfh cells from a distance to increase the likelihood of productive help, which is reflected by the elevated Tfh cell speed preceding interactions with WT GC B cells^18^. Our results extend this chemokine-based model by demonstrating that WT GC B cells also perform intrinsic pre-contact motility adjustments that actively position them for Tfh engagement. In contrast, MT EZH2 GC B cells fail to generate this coordinated search behavior, indicating that their interaction deficit begins before physical contact and is encoded at the level of cell motility rather than the affinity alone. We also demonstrated enhanced reliance on FDCs. Importantly, these single-cell imaging findings show that this increased FDC dependency is not just observed at the molecular scale but is behaviorally encoded through biased migration and subsequent localization toward the dLZ, which contains a dense FDC network.

This finding is important because a recent study showed that only FDCs located in the dLZ serve as long-lived antigen reservoirs, due to elevated CR2 expression that prolongs retention of the immune complex^35^. Consistent with this, we showed that NP-PE was enriched in FDC-dense dLZ region, where MT GC B cells remained preferentially localized and maximized antigen encounter. This spatial importance of FDC function implies that the biased migration of MT GC B cells toward the dLZ may serve not only to enhance survival and differentiation cues but also to ensure exposure to the most antigen-rich FDCs.

The FDC dependence of MT EZH2 GC B cells was further extended by our clarification of the temporal ordering in cell behavioral interactions, showing that FDC engagement precedes and appears to prime subsequent Tfh cell interactions. This FDC-first requirement, which was not observed in WT GC B cells, suggests a compensatory mechanism by which MT GC B cells may partially restore Tfh cell engagement once localized to the dLZ. Although the surface engagement of Tfh interactions with MT is impaired, our findings show that physical Tfh contacts with MT GC B cells increase specifically within the dLZ. This supports the premise that FDC-mediated positioning can functionally modulate Tfh signaling. Further, the requirement of FDC engagement prior to Tfh interaction may reflect a selection strategy that maximizes responsiveness to limiting antigen, thereby facilitating subsequent Tfh-mediated selection. Additionally, MT GC B cells displayed WT-like motility behaviors in the dLZ in contrast to unique and dynamic motility behaviors in the DZ, sLZ, and transitional zones (**Figure 3d**). Our behavior-transcriptomic integration analysis using scMOTIPh revealed enriched molecular programs linked to survival, antigen presentation, and metabolism of MT in the dLZ. This finding further confirms the heightened growth signals that support MT GC B cells in the dLZ (**Figure 7g**). These results suggest that MT EZH2-driven alterations in GC B cell motility and spatial preference represent an adaptive response to preserve key selection processes within a disrupted signaling landscape, leveraging the spatial organization of the antigen-retaining dLZ FDC network, as a scaffold for functional compensation.

Variation in GC B cell behavior by MT cell abundance may reflect a shift in competitive strategy during local clonal expansion. MT EZH2 GC B cells acquire a GC fitness advantage, outnumbering WT cells even more when MT cells were initially underrepresented^13^. Our results suggest a behavioral basis for this early competitive advantage. In WT rich GCs, reflective of early lymphomagenesis, enhanced migration with limited Tfh cell engagement may represent an exploratory search state that increases spatial sampling of the GC microenvironment that leads to preferential occupancy of the antigen rich dLZ. As MT cells become more prevalent, their shift toward reduced motility and greater Tfh cell engagement may reflect a transition toward niche stabilization, reinforcing clonal expansion and niche dominance. Thus, the behavioral changes we observe reveal a continuum of adaptive strategies from early phases of clonal seeding and expansion to later stages of dominance that offers a conceptual link between GC dynamics and lymphoma progression.

Our observation that MT EZH2 GC B cells exhibit increased spatiotemporal and behavioral heterogeneity across microenvironmental zones is also consistent with the recent findings that MT EZH2 drives widespread epigenetic reprogramming and increases epigenetic heterogeneity in B cell lymphomas, extending beyond H3K27 methylation changes^50^. Collectively, these observations of mutant GC B cells suggest that increased single-cell motility and positional plasticity may stem from increased epigenetic diversity, enabling individual cells to adopt a broader range of transcriptional and behavioral states in response to microenvironmental cues. Such changes in motility could disrupt normal GC zoning and lead to abnormal interactions with stromal and immune cells. Such disruption is directly linked to B cell fate and can contribute to malignant progression by allowing cells to escape the spatial constraints and checkpoint controls that typically restrict B cell expansion and selection within the GC. These findings highlight a functional consequence of EZH2-driven epigenetic remodeling that may contribute to early steps in lymphomagenesis through both cell-intrinsic and cell-extrinsic (i.e., microenvironmentally-driven) mechanisms.

Our scMOTIPh framework reframes the concept of a “cell” in GC biology by mapping transcriptomic programs onto dynamic behavioral phenotypes such as motility, interzonal trafficking, and cell–cell interactions. This multi-scale mapping bridges static molecular readouts with live imaging data, enabling bidirectional information flow between mechanistically rich transcriptomic patterns and cell behaviors that directly capture adaptation and decision-making at high spatiotemporal resolution. In effect, scMOTIPh implements a spatiotemporal-omics perspective that enables the study of previously unexplored dynamics of lymphomagenesis across space and time within live tissues, an essential step towards deciphering the heterogeneous mechanisms driving malignant transformation.

Beyond the GC context, this integrated framework supports the development of a cell state embeddings that are aligned with emerging “virtual cell” or “cellular digital twin” paradigms ^51^. One aspect of these frameworks is the ability to conduct in silico perturbations to predict gene expression outcomes from genetic or pharmacological perturbations by leveraging gene-gene relationships (e.g., gene regulatory network, gene ontology) or chemical structures of drugs^52–55^. scMOTIPh naturally integrates with these in-silico perturbation models by propagating predicted expression changes through its learned molecular-behavior mapping, thereby potentially enabling the prediction of behavioral outcomes based on specific perturbations or interventions. For example, EZH2 inhibitors have shown clinical activity and favorable therapeutic performance in GC-derived lymphomas^56^, in part by restoring lymphoma B cell immunogenicity and B–T immune synapse function^14^. By combining perturbation models with scMOTIPh, we can putatively predict how EZH2 inhibitor alters gene expression programs, and, in turn, how behavior dynamics are reconfigured in both WT and MT B cells, while also defining the precise timeline of immune events throughout the GC response. More broadly, this approach can expand towards building a ‘virtual GC’ model that could serve as a next-generation screening platform to identify molecular drivers of early lymphomagenesis, and design therapeutics capable of reprogramming pathogenic behaviors into non-pathogenic trajectories to shape GC outcomes.

## Acknowledgements

We thank Dr. Hao Shen for managing and maintaining the mouse colonies. W.B. was supported by NIH NCI R01 CA270245, LLS TRP 6641-22, LLS TRP 6679-24, FLF CURE FL 224362, LRF FL PRG 226504, ASH Junior Faculty Scholar Award 202414 and 223390, and Gilead Sciences Research Scholars Program GSI 231647-01. J.M.P. was supported by NIH NIGMS R35 GM157099, and the 2024 Salisbury Family and Center for Innovative Medicine Human Aging Project (HAP) Scholar. C.X. was supported by NIH R21 CA277513.

## Author contributions

Conceptualization, C.M., K.C., J.M.P., and W.B.; Methodology, C.M., K.C., X.C., I.K., J.M.P., and W.B.; Software and Formal Analysis, C.M., K.C., X.C., N.S., and J.M.P.; Investigation, C.M., K.C., X.C., J.M.P., and W.B.; Resources, C.X., A.M.M., J.M.P., and W.B.; Writing – Original Draft, C.M., K.C., J.M.P, W.B.; Writing – Review & Editing, C.M., K.C., X.C., C.X., A.M.M., J.M.P., and W.B.; Visualization, C.M., K.C., and X.C.; Supervision, J.M.P., and W.B.; Funding Acquisition, C.X., A.M.M., J.M.P., and W.B.

## MATERIALS AND METHODS

### Mouse models

We conducted bone marrow transplantation to generate a mouse model with multicolor fluorescently labeled GC cells. Bone marrow cells were harvested from 8–12-week-old female donor mice. The donors for MT and WT EZH2 GC B cells were generated by crossing Cγ1-cre mice (#010611, The Jackson Laboratory) with EZH2^Y641F^ mice^26^. For fluorescent labeling with CFP or YFP, these mice were additionally crossed with R26-CFP^57^ or R26R-EYFP mice (#006148, The Jackson Laboratory), respectively. The donors for tdTomato^+^ Tfh cells were generated by mating CD4-Cre mice (#022071, The Jackson Laboratory) with R26R-tdTomato mice (#007909, The Jackson Laboratory). Fluorescently labeled MT and WT EZH2 bone marrow cells, together with R26R-tdTomato;CD4-Cre bone marrow cells, were mixed at ratios of 12.5%, 12.5%, and 12.5 – 50%, respectively, and injected intravenously into lethally irradiated C57BL/6J host female mice (#000664, The Jackson Laboratory). Irradiation was conducted with 450 rads the day before and an additional 450rads two hours before transplantation. Each injection contained a total of one million bone marrow cells, including unlabeled (“colorless”) MT and WT EZH2 cells at 31.25 – 12.5% each. To generate control mice containing only WT EZH2 GC B cells, bone marrow cells from CFP^+^ WT EZH2 and YFP^+^ WT EZH2 donor mice were transplanted. For immunization, 50 μl of an NP-OVA solution (2 mg/ml in PBS; N-5051, Biosearch Technologies) mixed 1:1 (v/v) with alum was injected into the mouse footpad 9-11 days before imaging. To label FDCs in the popliteal lymph node, 2 μg (30 μl) of an anti-CD35 antibody (558768, BD Bioscience) conjugated to Alexa-Fluor-594 or Alexa-Fluor-647 (A20185 or A20186, Thermo Fisher) was injected into the mouse footpad approximately 12 hours before the imaging. For CD40L inhibition experiments, 20 μg of anti-CD40L antibody (BE0017-1, BioXCell) or isotype control IgG (BE0091, BioXCell) was added into the anti-CD35 antibody solution for co-injection (total 30 μl). Mice in which GC B cells were too densely packed to allow individual cell tracking were excluded from imaging analyses. All imaging and analyses presented in this study were conducted using pooled data from both bone marrow transplantation configurations.

For B1-8^hi^ adoptive transfer experiments, splenic B cells were isolated from B1-8^hi^ EZH2-mutant donor mice (*B1-8^hi^; Cγ1-Cre; Ezh2^Y^*^641^*^F^; Rosa26^YFP^*) and B1-8^hi^ EZH2 wild-type donor mice (*B1-8^hi^; Cγ1-Cre; Rosa26^CFP^*). Spleens were harvested and processed into single-cell suspensions, and total B cells were enriched by negative selection using the EasySep^TM^ mouse B cell isolation kit. Purified WT and EZH2-mutant B cells were counted and mixed at a 1:1 ratio. A total of 5 × 10^6^ B cells, consisting of 2.5 × 10^6^ WT CFP^+^ B cells and 2.5 × 10^6^ EZH2-mutant YFP^+^ B cells, were transferred intravenously via the tail vein into C57BL6 recipient mice. One day after adoptive transfer, recipient mice were immunized by subcutaneous footpad injection with 50 µL of a 1:1 mixture of NP-OVA (2 mg/mL in PBS) and alum to induce a GC response. 8 days after immunization, mice received anti-CD35-AF647 antibody by footpad injection. For antigen-tracking experiments, mice received 5 µg NP-PE per mouse prior to imaging. Imaging was performed at 24 h after NP-PE administration, with each imaging session lasting 30-60 min.

All animal experimentation and housing procedures complied with institutional guidelines established by Cornell University Institutional Animal Care and Use Committee, the Weill Cornell Medical College, the Guide for the Care and Use of Laboratory Animals (National Academy of Sciences 1996), and the Association for Assessment and Accreditation of Laboratory Animal Care International. Mice were maintained in the Cornell University animal facility under conventional housing conditions (22.3–22.7°C; 38–40% humidity; 12-h light/dark cycle) with ad libitum access to food and water.

### Intravital two-photon microscopy

Four to six months after bone marrow transplantation, intravital two-photon (2P) microscopy was conducted. Popliteal lymph node (LN) preparation for intravital imaging followed a. previously described protocol^29^.

Mice were anesthetized with 1–1.5% isoflurane in oxygen during surgery and imaging. The leg was shaved using an electric shaver and hair removal cream (Nair). AN incision was made in the skin behind the knee, and the fatty tissue covering the popliteal LN was removed with microdissection forceps. The exposed area was filled with saline and covered with a glass coverslip. The space between the coverslip and leg was sealed with grease to prevent evaporation of saline.

An implantable temperature probe (IT-23, Braintree Scientific) was positioned near the LN (beneath the coverslip) to monitor temperature, which was maintained at 36.5 ± 0.5°C using a nichrome heating wire placed above the coverslip. Core body temperature was maintained at 36–37°C with a temperature controller (FHC) consisting of a rectal probe and a heating pad.

Images were acquired with a two-photon microscope (Bergamo II, Thorlabs) equipped with ThorImage 4.1 software and a 1.05 NA objective (XLPLN25XWMP2, Olympus). The excitation source was a mode-locked Ti:sapphire laser (Chameleon Vision-S, Coherent) operating at 80 MHz pulse repetition rate. The pulse duration under the objective (measured by second-order interferometric autocorrelation) was approximately 90 fs. The diameter of the excitation beam at the back aperture of the objective (∼15 mm diameter clearance aperture) was determined by knife-edge scanning and measured at approximately 16 mm (1 / e^2^).

Simultaneous detection of CFP, YFP, tdTomato and AlexaFluor594 used 475 ± 21 nm, 525 ± 25 nm, 585 ± 15 nm, and 623 ± 16 nm band-pass-filters (FF01-475/42, FF03-525/50, FF01-585/29, FF01-623/32, Semrock). Fluorescence separation was achieved with 488 nm, 562 nm, and 594 nm dichroic-beam-splitters (Di02-R488, FF562-Di03, Di02-R594, Semrock).

For experiments using AlexaFluor647 (instead of AlexaFluor594), tdTomato and AlexaFluor647 were detected with a 607 ± 35 nm band-pass-filter and 633 nm long-pass-filter (FF01-607/70, BLP01-633R, Semrock) separated by a 649 nm dichroic-beam-splitter (FF649-Di01, Semrock).

Images were acquired at 256 × 256 pixels (230 μm x 230 μm field of view) at 59 Hz and averaged over 30 frames. A z-stack of 30 planes with 3-μm step size was collected every 30 seconds.

### Image processing

For the processing of 3D live imaging data, we used Imaris software v.10.0 (Oxford Instruments) for visualization, instance segmentation, single-cell tracking, and extraction of basic statistics. To segment GC B cells, Tfh cells, and auto-fluorescent macrophages, we utilized the Surface module, applying Gaussian smoothing and background subtraction thresholding with a sphere diameter of 5 µm to detect moving cell objects while reducing image noise. Single-cell instances were captured by conducting morphological splitting based on the distance transform and watershed algorithm with a 5 µm diameter to ensure accurate cell separation. Volumetric and edge filters were applied to retain objects larger than 10 voxels and to exclude objects touching the image boundaries.

To ensure the accuracy of single-cell segmentation for each cell type, we manually inspected and corrected mislabeled objects by iteratively training the built-in machine learning classifier until the segmentation passed visual inspection. FDCs were segmented using the same Surface module, but Gaussian smoothing and background subtraction thresholding were disabled, with a sphere diameter of 2 µm to preserve the detailed branching structure of FDC networks. No morphological splitting was performed for FDCs, treating them as a continuous meshwork. For FDC segmentation, volumetric filters were applied to retain objects larger than 15 voxels. For tracking the movement of single cells, we employed the Tracks module using the Brownian Motion algorithm, which is based on the Crocker-Grier algorithm for particle tracking^58^.

To optimize tracking performance, we set the maximum linking distance between frames to 15 µm and the maximum gap size to 0, preventing linking of cells that moved more than 15 µm in a single time step and eliminating trajectories with high ambiguity or noise. Data was excluded in cases where tracks were frequently misconnected, making the tracks appear unsmooth (mostly due to excessively fluorescence-labelled GC cells). The extraction of single-cell features over time was achieved using the Statistics module. This allowed us to obtain key morphological features, 3D coordinates, and colocalization data for each cell type. These features were subsequently used to calculate morphodynamic properties, such as cell volume and shape changes, and motility features, including speed, and directionality. Colocalization features were computed to capture the spatial proximity and interaction dynamics between GC B cells and FDCs and Tfh cells.

For B1-8^hi^ adoptive transfer model image processing, tissue drift was corrected by estimating translations from the FDC channel by Gaussian-smoothing each FDC volume at physical standard deviations of 3.0, 0.9 and 0.9 µm in z, y and x, respectively, background centered by subtracting its 20th percentile, clipped at zero and downsampled twofold in x and y. Consecutive axial displacements were estimated by 3D phase cross-correlation with tenfold subpixel upsampling. Lateral displacement was estimated from maximum-intensity projections by pyramidal Lucas–Kanade tracking of up to 750 Shi– Tomasi FDC features using a 31 × 31 pixel window and three pyramid levels. B cell channels were subjected to smooth-background subtraction using Gaussian standard deviations of 9, 15 and 15 µm in z, y and x, followed by Gaussian denoising at 1.5, 0.7 and 0.7 µm. NP–PE underwent background subtraction at 6, 8 and 8 µm and Gaussian denoising at 1.5, 0.6 and 0.6 µm. The FDC channel underwent background subtraction at 12, 25 and 25 µm and denoising at 1.5, 0.6 and 0.6 µm. B cells were segmented independently using the Cellpose-SAM model^59^ with native three-dimensional inference with the measured z-to-xy anisotropy, automatic intensity normalization, a cell-probability threshold of 0, flow threshold of 0.4 and model-estimated diameter. Objects smaller than 15 Cellpose pixels were rejected during inference, and three-dimensional instances smaller than 50 µm³ were subsequently removed using the physical voxel volume. FDC channel was smoothed with an isotropic physical Gaussian standard deviation of 0.8 µm. A single threshold for the entire movie was obtained by Otsu thresholding positive values sampled from up to nine evenly spaced frames. Binary masks were cleared of components smaller than 30 µm³ and closed with a 1 µm radius ellipsoidal structuring element defined in physical space. The same fixed threshold was applied to every frame. The NP-PE antigen channel was Gaussian-smoothed with a 3 µm physical standard deviation, and a movie-wide Otsu threshold was estimated from up to nine sampled frames. Components smaller than 30 µm³ were removed, and the mask was closed with a 1 µm physical ellipsoid. NP-PE spots were detected in 3D by a scale-normalized blob response based on the negative Laplacian of Gaussian (LoG). The target spot radius was 1 µm, converted independently along z, y and x, with Gaussian sigma equal to radius divided by √3. Candidate local maxima were required to exceed the median LoG response by six robust standard deviations, where robust standard deviation was 1.4826 x MAD. Non-maximum suppression used a 1.5 µm ellipsoidal footprint. Candidates were additionally required to exceed the 99.8th percentile of positive NP-PE intensity in the same frame.

### Trajectory processing

To eliminate biased influence of trajectory length or direction, we standardized all trajectories to a duration of 10 minutes for quantification of motility and colocalization features. Trajectories shorter than 10 minutes were excluded to ensure high-quality tracking, while those longer than 20 minutes were divided into multiple segments to maximize sample size, and only the first 10-minute segment was used. To account for directional independence, which can vary based on the region of interest, we defined a primary migration axis based on the maximum displacement over time, a secondary migration and a tertiary migration axis that are orthogonal^60^. We derived these axes by applying singular value decomposition (SVD) to the displacement matrix *A*, expressed as:

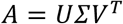

Here, *U* is a matrix of eigenvectors of *AA^T^*, *V^T^* is a matrix of eigenvectors of *A^T^A*, and Z is a diagonal matrix of singular values. The matrix of coordinates *R* was then multiplied by *V* to obtain *R_reg_* = *RV*, where the first, second and last columns of *R_reg_* represent the movement paths along the primary, secondary and tertiary axes, respectively. Additionally, all trajectories were translated to set the initial point as the origin.

### Quantification of single-cell motility features

To describe the single-cell motility of GC lymphocytes, we evaluated a total of 123 motility features. The motility features included displacement-based metrics and turning angle-based parameters designed to capture anisotropic persistent random walk (APRW)^61^, displacement magnitude, tortuosity, displacement distribution descriptors, angle distribution descriptors, signal descriptors, temporal correlations, elongation, sphericity, entropy, and decomposed motility metrics. These features provided a comprehensive characterization of cell motility patterns and their statistical distributions over time. A complete list of motility features is included in **Supplementary Table 1**.

### Quantification of cellular colocalization features

To assess the dynamic context of interactions within the GC microenvironment, we extracted a total of 138 colocalization features. These features comprehensively captured diverse aspects of cellular interactions, including distant interaction, spatial positioning, rate of change in position, contact interaction, and residence time (time spent in a specific vicinity/subregion). These colocalization metrics allowed us to quantify the spatial and temporal relationships between GC B cells, Tfh cells, and FDCs, providing insights into their interaction dynamics within the GC architecture. A complete list of colocalization features is shown in **Supplementary Table 2.**

### Quantification of morphological changes/plasticity

We constructed a three-dimensional morphology PCA space using 5 morphology features that were derived from the Imaris v.10.0 output, including surface area, volume, sphericity, oblate ellipticity (i.e., shaped more like a spheroid, soccer ball), and prolate ellipticity (i.e., shaped more like an ellipsoid/football). For each time duration, morpho-trajectories traversing the three-dimensional morphology space were recorded and average displacements within the space were calculated to capture morphological changes from multidimensional morphology descriptors. For example, cell shapes that colocalized with a similar region of the three-dimensional morpho-PC space defined cells with minimal morphological changes, and cells that spanned large distances within the morpho-PC space defined cells with large morphological changes.

### Identification and classification of motility clusters (MCs)

To identify and classify single-cell motility behaviors we implemented a modified version of our CaMI pipeline^20^. Here, we completed a principal component analysis (PCA) on the multidimensional feature dataset to eliminate covariances between parameters and utilized principal components (PCs) accounting for 95% of the cumulative variance to create two-dimensional UMAP projections^62^. For identifying MCs, we applied the K-means++ unsupervised algorithm to identify nine distinct motility clusters from the PCs. To validate these clusters, we visualized the trajectories corresponding to each motility cluster, allowing for a qualitative assessment of trajectory similarities. Additionally, we quantified the heterogeneity of motility states by calculating the Shannon entropy *S*.

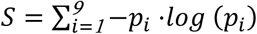

Here *p_i_* is the fraction of cells within each motility cluster *i*.

### Enrichment and depletion of MCs and pre/post contact modules

To assess whether specific motility clusters (MCs) or behavior modules were significantly enriched or depleted in a given condition (e.g., cell type, or zone), we performed a randomized permutation testing. For each condition, we computed the observed frequency of a specific MC or module. To generate the null distribution. We randomly shuffled group labels while preserving the overall number of cells and repeated this procedure for N iterations, computing the simulated frequency of the same MC or module in each iteration. Formally, for observed statistic *x_obs_* and simulated statistics *x_1_*, *x_2_*, ⋯, *x_N_*:

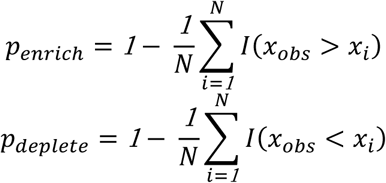

Here *I*(·) is the indicator function. *p_enric_*_ℎ_ and *p_deplete_* denote the statistical p values for the enrichment and depletion, respectively.

### Motility cluster (MC) transition analysis

Each cell was longitudinally tracked until it disappeared from the field of view. To enable detailed analysis of motility state transitions, each cell trajectory was subdivided into 10-minute temporal compartments, during which the corresponding MC was assigned based on the extracted motility features. Trajectories that did not contain at least two distinct MCs were excluded from further analysis. Transitions between MCs were recorded for all remaining trajectories, capturing the prior and post-transition MC states for each cell. Transition probabilities were then calculated by normalizing the number of observed transitions by the total number of transitions for each MC, enabling us to quantify the likelihood of cells moving between different motility clusters over time.

### Bulk RNA-seq and analysis

Methods for the preparation of bulk RNA-seq for centroblasts and centrocytes were described previously^13^. Briefly, Naive B cells, centroblasts, and centrocytes were sorted from spleens of Cγ1-cre and Ezh2(Y641F)^fl/WT^;Cγ1-cre mice 8 days post-SRBC immunization. For each sample, 50,000–100,000 cells were collected in TRIzol (Invitrogen), and total RNA was extracted per manufacturer’s protocol. RNA quality was assessed using the Agilent 2100 Bioanalyzer, and only samples with RNA Integrity Number (RIN) ≥ 8 proceeded to library preparation. mRNA libraries were prepared with the Illumina TruSeq RNA kit using poly(A) selection, fragmentation, and cDNA synthesis, followed by adapter ligation and 15 cycles of PCR amplification. Libraries were quality-checked and pooled with unique indices, then sequenced (50 bp single-end reads) on an Illumina HiSeq 2500 platform to a depth of ∼80 million reads per sample. Sequencing data were processed using Illumina RTA and demultiplexed with CASAVA 1.8.2. Adapter trimming was performed with Trim Galore! (v0.4.1), and reads were aligned to the mm10 genome using STAR (v2.5.1b) with Gencode M12 annotation. Gene-level quantification was performed with featureCounts (subread package). Differentially expressed genes (DEGs) were obtained by performing DESeq2 on gene counts comparing MT to WT based on log2 fold-change > 1 and p value < 0.05 after adjusting p value with Benjamini-Hochberg correction (**Supplementary Tables 3 and 4**). Gene Set Enrichment Analysis (GSEA) was performed using gseapy enrichr method based on the gene ranking of log2 fold-change statistics from DESeq2. Pathways were from Molecular Signatures Database (MSigDB) and community detection of those were performed using Louvain method with resolution = 1.

### Motility associated gene signature analysis

From DEGs, we determined that 83 genes were commonly upregulated and 314 genes were downregulated in MT in both centroblasts and centrocytes. After extensive literature curation, we identified 49 genes showing well-established association with cell motility such as Rho GTPase signaling, chemotaxis and cytoskeleton remodeling as well as literatures^33,34^ (**Figure S5a**, **Supplementary Table 6**).

### Defining FDC density-dependent zones

To define FDC density-dependent zones within the GC, we generated a density map from the segmented FDC masks. The FDC masks were derived from the 3D segmentation of FDC networks, capturing the spatial organization of cells within the GC. A convolution operation was applied to the segmented masks using a 3D kernel, which allowed us to compute the spatial correlation of FDC density and create a smoothed representation of the FDC distribution. Once the convoluted density map was generated, we employed adaptive thresholding to define specific density-dependent zones. This thresholding was reached in a manner that preserved the initial volume of the LZ, ensuring that the LZ occupied approximately 30–50% of the total region of interest (ROI), depending on the specific image being analyzed. Within the LZ, the dLZ was defined as the top 1-10% dense region of the LZ volume, which constitutes approximately 10% of the total LZ volume. DZ was defined as the rest of the region that has minimal FDC density. To label cell trajectories based on their location within these zones, we defined the zone of each trajectory according to the duration that the cell spent in a particular zone. Specifically, if a cell resided within a particular zone for 60% or more of its trajectory duration, it was labeled as belonging to that zone. This labeling allowed us to categorize the behavior of cells relative to the FDC density in different parts of the GC, generating the non-linear, time-varying boundaries.

### Quantification of interzonal transitions

For each longitudinal zonal recording of the cells, we generated lists of batches with size *T_step_*, where *T_step_* represents the time step for the next transition in the zonal state. For each batch, the zone state was defined based on the same criteria (60% or more residence time) as the representative state to summarize the zonal behavior of the cell during that time interval. This method ensured that transient or less significant fluctuations in the zonal state were minimized, providing a robust indicator of the cell’s primary location within the germinal center. Transitions between these representative zone states were recorded for each cell, capturing both the prior and post-transition states. This allowed us to track how cells moved between different zones over time and to analyze patterns of spatial organization within the GC.

### Monte Carlo simulation of interzonal transition using a Markov chain

To model the steady-state distribution of GC B cells across defined GC zones, we performed a Monte Carlo simulation based on a Markov chain model describing transitions between five zones: DZ, DZ-sLZ, sLZ, sLZ-dLZ, and dLZ. Cell trajectories were divided into 8-minute intervals and assigned to zones according to where the cells spent the majority of their time. From longitudinal history of zone transitions per cell, we counted every transition at next time step and obtained the transition probability matrix by transforming counts to probability. We initialized the simulation from sLZ and performed 10^6^ Monte Carlo iterations, where at each step the cell state was updated based on the probabilistic transition rules. This process was repeated until the system converged to a steady-state distribution, which represents the long-term occupancy probabilities of GC B cells in each zone under the given transition dynamics.

### Quantification of EZH2 MT interaction preference score

To evaluate the relative preference of GC B cells for Tfh interactions, we compared the frequency of GC B cell–Tfh cell contacts between MT and WT GC B cells across all interaction-positive instances. Only GC regions and time frames in which MT–Tfh interaction frequency exceeded that of WT were included in the analysis to specifically capture MT-favored contexts. Tfh interaction events were categorized based on their duration into low-persistence interactions (0–6.5 minutes) and high-persistence interactions (7.5–10 minutes). For each persistence category, we computed the difference in Tfh interaction frequency between MT and WT GC B cells. This metric was used to assess whether MT GC B cells were disproportionately associated with shorter or longer Tfh engagements compared to their WT counterparts.

### Tracking pre and post contact cell behaviors and module identification

To investigate the motility behavior of GC B cells pre-, during, and post-long-lasting interactions with Tfh cells, we initially identified and extracted cell trajectories associated with sustained interactions, which were defined as continuous contacts maintained between GC B cells and Tfh cells for 7 minutes. To analyze emergent motility pre/post contact, we computed two instantaneous motility metrics at every time point– speed and turning angle – providing insights into the magnitude of the movement or directional shifts, respectively. Each trajectory was then divided into three distinct interaction phases: pre-contact, representing the period before the cell initiated contact with the interaction partner; during contact, covering the duration of sustained contact; and post-contact, representing the period after the interaction ceased. The average motility for each phase was calculated by averaging the instantaneous speed and turning angle values across all time points within that phase. To analyze dLZ localization pre/post contact, we used distance to dLZ at each time point and quantification was done likewise.

To identify distinct patterns of behavioral dynamics before and after the Tfh interaction, we conducted time-series K-means clustering using Euclidean distance on the time-series data of standard-scaled instantaneous speed, turning angle, and distance to dLZ across the three interaction phases (pre, during, post). The clustering allowed grouping of trajectories into four modules that captured common temporal patterns of behavioral change. Each module was defined by the barycenter (mean trajectory) of the cluster, and descriptive names were assigned based on the characteristic shape of these barycenters across the pre-contact, contact, and post-contact phases. These modules allowed us to categorize GC B cell responses to Tfh engagement into discrete, interpretable behavioral classes.

### Single-cell RNA-sequencing and preprocessing

Methods for the preparation of single-cell RNA-seq for centroblasts and centrocytes were described previously^13^. Briefly, YFP⁺IgD⁻ splenocytes were sorted from R26-lox-stop-lox-YFP;Cγ1-cre and R26-lox-stop-lox-YFP;Ezh2(Y641F)^fl/WT^;Cγ1-cre mice 8 days after SRBC immunization (n = 3 mice per group). For each sample, ∼10,000 sorted cells were processed using the 10X Genomics Chromium Single Cell 3′ v2 platform, and libraries were prepared following manufacturer protocols. Sequencing was performed on an Illumina HiSeq 4000 (R1: 26 bp, I7: 8 bp, R2: 98 bp) with an average of 250 million reads per sample. Raw reads were processed with Cell Ranger v3.0.2 (10X Genomics) using default parameters for demultiplexing, alignment to the mm10 genome, UMI counting, and gene quantification. On average, we obtained ∼2,853 cells per sample, with 100,000 reads/cell and 77% sequencing saturation. Across libraries, we detected ∼14,625 mean genes per library, 1,448 median genes and 3,654 median UMIs per cell.

We used variational autoencoder-based scVI Solo model^63,64^ to conduct doublet detection and removed ∼15% of cells predicted to be doublets based on the top 2000 genes with presence of at least 10 cells. We used the scanpy package^65^ to perform quality control of the genes, and cells to determine the inclusion for downstream analysis. Cells with bottom 2% gene counts and mitochondrial gene expression fraction more than 20% were removed from the analysis. To identify GC B cell population and remove plasmablasts, we performed Leiden clustering with resolution of 0.06 based on the 50 PCs.

### Identification of GC B cell transcriptomic subtypes

After removing the plasmablasts, we recomputed the top 3000 variable genes and performed Leiden clustering of resolution 1.4 and UMAP based on the principal components up to 40% variance. We identified differentially expressed genes (DEGs) of each Leiden cluster by rank_genes_groups method in scanpy based on Wilcoxon-ranksum test with Benjamini-Hochberg correction. Subtype annotations were based on the gene signatures collected from the literature^36^ and our manually collected gene sets. Scores on the expression of manually collected gene sets are calculated by score_genes method in scanpy that subtracts average expression of randomly sampled reference genes from the average expression of the target gene set. Gene sets from the literature^36^, top 100 genes based on the log-fold change were extracted for each subtype. Community detection is based on the IoU score, and the community is generated by Louvain algorithm with a resolution of 1.6.

### Identification of Behavior clusters (BCs) and analysis

GC microenvironmental BCs were based on integrated behavior space that comprises 123 motility (**Supplementary Table 1**) and 96 curated colocalization features (**Supplementary Table 2**) that encode zone, Tfh cell and FDC interaction. We applied the K-means++ unsupervised algorithm to identify eight distinct BCs from the PCs with variance up to 75%.

### Inferring single-cell transcriptomic profiles on behavioral space

To link behavioral states with scRNA-seq profiles, we developed scMOTIPh (single-cell Multiscale Optimal Transport for Imputing Phenotypes), a computational framework that solves alignment and mapping between molecular and cellular scale. scMOTIPh is formulated as an optimal transport problem^37,38^ designed to satisfy three key objectives: (1) maintaining global structural integrity—preserving the variance and overall geometry of the original transcriptomic and behavioral manifolds; (2) preserving local structural correspondence—ensuring that neighborhood relationships within each modality remain coherent after transformation; and (3) incorporating prior biological knowledge, such as known cell labels and motility magnitude, to guide the mapping to be biologically meaningful.

If 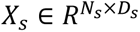 and 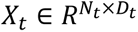 represents a data matrix for *N_s_* and *N_t_* cells from source and target space, respectively. Here, *X_s_* and *X_t_* represent single-cell molecular (scRNA-seq) and behavior profiles across *D_s_* and *D_t_* features, respectively. We sought to solve Fused Gromov-Wassterstein optimal transport problem^39,66^, which is defined as:

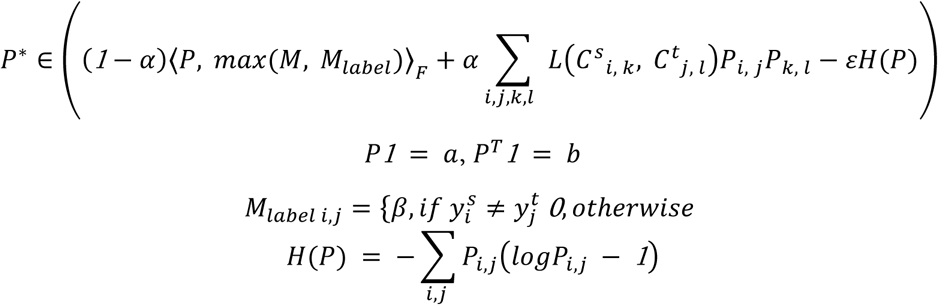

We solve for *P*^∗^, which is optimal coupling matrix from possible set of coupling matrices 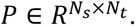 Here, ⟨·⟩*_F_* is Frobenius inner product, *a* and *b* are marginal distributions over source and target space, respectively, and are assumed uniform for simplicity. 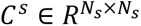 and 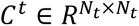 refer to pairwise Euclidean distance matrix in the source and target space defined by principal components that capture 40% and 75% variance, respectively. The distance matrices are used to compute squared Euclidean loss function *L*, which preserves local structural correspondence. Shared feature cost matrix 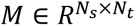 and label cost matrix 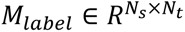 are critical components to guide motility and label-informed mapping, respectively. *M* is the squared Euclidean distance matrix between motility scores based on the curated set of motility associated genes computed by score_genes function in scanpy and average speed values in behavior space. In this semi-supervised learning approach, *M_label_* penalizes *M* if cell label vector (DZ, DZ-LZ, and LZ) *y^s^* and *y^t^* in the source and target space are mismatched, ensuring both alignment of GC microenvironment and motility. *α* is a trade-off parameter between structural correspondence and biological knowledge, *β* is a penalty coefficient for label mismatch, and *ε* is a regularization strength that controls entropic regularization term *H*(*P*). After solving for *P*^∗^ with *α* = *0*.*1*, *β* = *100*, and *ε* = *1*.*3*, we can quantify source space profiles in the transported target space 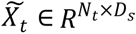 by:

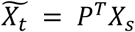

Consequently, 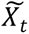 contains inferred transcriptomic profiles for every behavior state. Further downstream analyses consist of KNN-based imputation of BCs and MCs with *number of neig*ℎ*bors* = *15*, allowing inference of transcriptomic profiles of each BCs and MCs as well as validation of scMOTIPh by GSEA performed on each BCs.

### Transcription-based functional analysis of MCs

To connect motility-defined dynamics with transcriptomic signatures and pathway-level interpretations, we constructed a multiscale mapping between MCs, BCs, DEGs, and enriched pathway modules. To quantify the association between MCs and BCs, we computed the fractional contribution of each MC to each BC by tabulating the proportion of cells from a given MC that were assigned to each BC. Following the integration of behavioral and transcriptomic data via scMOTIPh, we performed DEG analysis across MCs. For each gene, a distinctiveness score was computed to prioritize MC-specific DEGs, defined as:

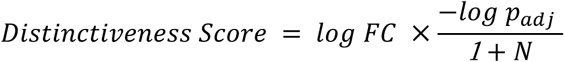

where N is the number of MCs in which the gene was significantly expressed (adjusted p < 0.05). This formulation favors genes that are selectively significant in a single or few MCs. For each MC, the top 300 genes with the highest scores were selected as MC-specific DEGs. To interpret these gene signatures at the pathway level, we performed GSEA using curated gene sets from MSigDB. Enriched pathways were identified for each MC-specific DEG list (adjusted p<0.05), and pathway similarity was assessed using the IoU score of gene content between significant pathways. Based on the resulting IoU similarity matrix, we applied Louvain community detection (resolution = 1.06) to group pathways into 10 functional modules. Each module was then manually annotated based on frequency of pathway terms and relevance to biological themes represented within the community.

### Statistics and reproducibility

To establish statistical comparisons between two groups we used the nonparametric Mann-Whitney U test unless stated otherwise. Any statistical comparisons between multiple groups, we performed Kruskal-Wallis test followed by Dunn’s post-hoc test unless stated otherwise. Correlations of linear regression fits were evaluated using r and p-values, respectively. Data was scaled, where applicable, to normalize distributions for gaussian-based models. The experiments were not randomized.

### Software

All 3D single cell segmentations and tracking were 7 using optimized workflows in Imaris. Behavior feature extraction was performed using a custom algorithm combining Imaris outputs with in-house algorithm. All analyses were performed in Python with the following software specifications: python 3.9.13, scipy 1.13.1, scikit-learn 1.1.13, tslearn 0.6.3, scikit-image 0.19.3, scikit-posthocs 0.9.0, statsmodels 0.13.5, pandas 1.5.2, matplotlib 3.6.2, seaborn 0.11.2, umap-learn 0.5.3, and numpy 1.23.5, cmcrameri 1.9, EntropyHub 0.2, pyarrow 12.0.1, scanpy 1.9.3, scvelo 0.3.2, scvi-tools 1.1.5, POT 0.9.5, napari 0.4.17, network 2.8.8.

## Code availability

The source code used to generate the data and findings presented in this work is deposited on the https://github.com/Phillip-Lab-JHU/scDynamics. The code for CaMI is available at https://github.com/PhillipLab-JHU/CaMI.

## Data availability

The GEO accession numbers are GSE138032 for bulk RNA-seq and GSE138033 for single-cell RNA-seq.

## FIGURE CAPTIONS

**Figure S1. Supplementary information for** Figure 1**. The imaging area in GC. a.** XY, ZY, and XZ section views (15μm-thick) at the solid lines in the large imaging volume (480 x 480 x 180 μm^3^). Light zone (LZ, yellow dotted line) and dark zone (DZ, blue dotted line) are identified by CD35+ FDC (AlexaFluor594) and the other regions in the GC, respectively. The GC is where GC B cells (CFP and GFP) exist. Macrophages (arrow) are identified in multi detection channels (shown as white color combined with blue, green and red) and are obviously larger than GC B cells and T cells. **b.** To acquire high speed timelapse volumetric images (Supplementary Video 1), the smaller volume (230 x 230 x 90 μm^3^) was selected (yellow box) from the shallowest part of the GC to 90 µm deeper, containing the entire LZ in depth direction. **c.** XY section views at the indicated depths. Yellow boxes indicate the area of timelapse volumetric imaging in the GC.

**Figure S2. Supplementary information for** Figure 1**. Optimization of 2P imaging. a.** Determine an optimal laser wavelength for simultaneous 4-color imaging. Fluorescent intensity changes with laser wavelength at the same laser power. Each data point in graphs shows the mean value of fluorescent intensity of 1-10 cells for CFP, YFP, and tdTomato. Each data point for AlexaFluor594 or AlexaFluor647 is the mean pixel value of the GC area in each image. We determined 880 nm as the optimal laser wavelength for simultaneous 4-color imaging. Scale bars, 50 μm. **b.** An example of high pulse energy-induced damage. Tracks of tdTomato+ T cells (red) in 0-10 minutes and 74-84 minutes at pulse energy of 0.34 nJ at focal plane. **c.** Determine a safe range of laser pulse energy at focus and laser average power at LN surface. Damage and safe cases are indicated by red and white color, respectively. Each data point represents each GC imaged over 90 or 40 minutes. The laser-induced damage is determined by the arrest of cell migration. Below the average power of 80 mW, damage seems to be more affected by pulse energy. We determined 0.23 nJ as the threshold of pulse energy. For calculating the pulse energy (*E*) at the focus, we used the following equation: *E* = (*P*/*R*)e^-^*^Z^*^/^*^L^*; *P*, average power under the objective lens; *R*, pulse repetition rate; Z, depth at the focus; *L*, effective attenuation length. For measuring the effective attenuation length (*L*), we measured the fluorescent intensity (*F*) at various depths (Z) and obtained the slope (-2/*L*) of the following equation through linear fitting: ln(*F* / P^2^) = - (2/*L*) *Z* + constant.

**Figure S3. Supplementary information for** Figure 1**. a.** Average Z score of motility features for each MC. Hierarchical clustering is based on the Euclidean distance using the Ward linkage. **b.** Thirty example 10-minute trajectories for each MC. N represents the number of tracks within each MC. **c.** Interpretable motility features projected onto the UMAP space. **d.** Cross-correlation between post-immunization days (D9, D10, D11) combined with bone marrow transplant configurations based on the occurrence of MCs. MT and WT EZH2 GC B cells were labeled with CFP and YFP, respectively, and labeled with YFP and CFP, respectively.

**Figure S4. Supplementary information for** Figure 2**. Bulk RNA-seq of centrocytes and centroblasts. a.** Volcano plot of DEGs between MT EZH2 and WT. Genes with *log*_2_*FC* > 3 or *log*_2_*p_adj_* > 40 are annotated. **b.** Number and list of overlapping genes from centroblasts and centrocytes. **c.** Pathway association based on gene IoU score. Colors represent group of pathways based on Louvain community detection. Red arrow indicates downregulation in MT GC B cells. **d.** GSEA on centroblasts between MT EZH2 and WT.

**Figure S5. Supplementary information for** Figure 2**. Identification of motility-associated genes and pathways for GC B cell. a.** Motility associated gene sets that significantly co-expressed in centroblast (CB) and centrocytes (CC). Log-fold change is between MT EZH2 and WT. **b-e.** Other gene sets that significantly co-expressed in centroblast and centrocytes associated with 3D amoeboidal migration (**b**), chemotaxis (**c**), chemokinesis (**d**), and morphological plasticity (**e**). Log-fold change is between MT EZH2, and WT. **f.** Pathway association based on gene IoU score. Colors represent group of pathways based on Louvain community detection. Motility associated signatures are highlighted in red. Pathways are associated with downregulation in MT GC B cells except for motility associated signatures, chemokinesis, chemotaxis, and 3D amoeboidal. **g.** GSEA based on motility associated genes between MT EZH2 and WT.

**Figure S6. Supplementary information for** Figure 3**. a-b.** KDE distribution of distance from GC B cells in each zone to the boundary of DZ (**a**), and dLZ (**b**). The highest probability distance is denoted for each zone. **c.** Distribution of Tfh cells for each zone (mean ± 95% C.I.). **d-e.** Two-dimensional UMAP KDE representation of multidimensional single-cell motility computed for each DZ and LZ (**d**), sLZ and dLZ (**e**). For each panel, top represents 1D KDE across UMAP1, right represents 1D KDE across UMAP2 (WT DZ n = 405, WT LZ n = 899, MT DZ n = 568, MT LZ n = 1,767, WT sLZ n = 660, WT dLZ n = 145, MT sLZ n = 1,301, MT dLZ n = 305, 10-minute track of cells).

**Figure S7. Supplementary information for** Figure 4**. a-f.** Zonal transitions across all elapsed times: DZ-sLZ to DZ (**a**), sLZ to DZ (**b**), sLZ to DZ-sLZ (**c**), sLZ to sLZ-dLZ (**d**), sLZ to dLZ (**e**), and sLZ-dLZ to dLZ (**f**) transitions (mean ± 95% C.I.). Data points represent the transition at each elapsed time (1-13 minutes). **g.** Maximum-intensity projection of an intravital imaging of GC 24 h after NP injection in the B1-8hi model. Left image shows WT (blue) and MT GC B cell (orange) channels, center image shows NP-PE (red) and CD35^+^ FDC (white) channels, and right image shows all four channels. Scale bars, 20 µm. **h.** Smoothed XY normalized density map of detected NP–PE spots. A total of 380 detected spots were projected and smoothed with a Gaussian kernel with σ = 12. The cyan outline denotes the projected dLZ boundary. Scale bars, 20 µm. **i.** NP–PE spots enrichment was calculated as the fraction of all detected spots located in a zone divided by the fraction of the volume occupied by that zone. Dashed line represents spot localization proportional to zone volume. Each point represents the median of a GC. Horizontal lines indicate group medians. P value is from Kruskal-Wallis test (n = 3 GCs per zone). **j.** Frequency of FDC contact of GC B cells within dLZ for each time duration. **k.** Average volume fraction of GC B cell overlapping with FDC in sLZ when direct contact occurred for 10-minutes. Each point represents a cell. White dot represents median, boundaries of black box represent first and third quartile. P value is from Mann-Whitney U test (WT n = 147, MT n = 358 cells). **l.** DZ residence duration. Each point represents a cell. Line inside box represents median, boundaries of the box represent first and third quartile. Lines outside box represent whiskers. P value is from Mann-Whitney U test (WT n = 405, MT DZ n = 568 cells). **m-n.** Difference in median FDC distance (**m**) and FDC contact duration (**n**) between MT and WT GC B cells within each GC against the relative abundance of MT GC B cells. r indicates Pearson correlation coefficient with corresponding p value (12 GCs). Each dot represents a GC. The line indicates the linear regression fit, and shading denotes the 95% confidence interval. Vertical dotted lines indicate the thresholds used to define WT-rich, mixed and MT-rich GCs.

**Figure S8. Supplementary information for** Figure 5**. a.** KDE distribution of overlapping volume fraction of GC B cell with Tfh cells. Dotted box highlights the overlapping volume fraction range from 0.05 to 0.2. **b.** Relative preference of GC B cells for Tfh engagement by calculating differential interaction frequency between MT and WT GC B cells and measuring the number of Tfh interactions where MT frequency exceeded that of WT (mean ± 95% C.I.). Interaction events were stratified by contact duration into low-persistence interactions (0–6.5 minutes) and high-persistence interactions (7–10 minutes). P value is from Mann-Whitney U test (Low n = 69, High n = 19 instances). **c-d.** Membership of pre/post contact motility modules (**c**) and four example members (**d**). **e.** Fraction occurrence of four pre/post contact motility module identified by time-series K-means clustering of GC B cell’s average speed (WT n = 30, MT n = 34 cells). Bottom, time-series cluster barycenter shows the representative fluctuation of speed for each module. Z_speed_ represents the average Z score of speed. **f-g.** Difference in Z score of speed between pre-contact and during contact with a Tfh cell (**f**), and between post-contact and during contact with a Tfh cell (**g**) (mean ± 95% C.I.). Each dot represents a cell. P values are from Mann-Whitney U test (WT Slow Precontact n = 13, MT Slow Precontact n = 17, WT Fast Precontact n = 17, MT Fast Precontact n = 17, WT Slow Postcontact n = 16, MT Slow Postcontact n = 17, WT Fast Postcontact n = 14, MT Fast Postcontact n = 17 cells). **h.** Fraction occurrence of four pre/post contact motility module identified by time-series K-means clustering of GC B cell’s volume (WT n = 30, MT n = 34 cells). **i-j.** Fraction occurrence of four pre/post contact motility modules identified by time-series K-means clustering of Tfh cells’ average speed (**i**), and average turning angle (**j**) (WT n = 19, MT n = 20 cells). Bottom, time-series cluster barycenter shows the representative fluctuation of feature for each module. Z_volume_, Z_speed_, and Z_Ө_ represents the average Z score of volume, speed, and turning angle.

**Figure S9. Supplementary information for** Figure 6**. a-b.** Fraction of GC B cells within sLZ (**a**) and dLZ (**b**) that directly contact a Tfh cell for more than 7 minutes per GC (mean ± 95% C.I.). Each dot represents one GC. P value is from two-sided unpaired t-test (WT n = 6, MT n = 6 GCs). **c.** Fraction occurrence of cells including low contact duration (less than 7 minutes but more than 2 minutes) based on four pre/post contact dLZ localization module identified by time-series K-means clustering of dLZ distance. Enrichment and depletion represent statistical over-representation and under-representation compared to random sampling of GC B cells, respectively (WT n = 197, MT n = 399 cells). **d.** Fractional occurrence of four pre/post contact dLZ localization module identified by time-series K-means clustering of dLZ distance. Low and high represents lower and higher than 7% of engaged fraction of surface with Tfh cell, respectively. Enrichment and depletion represent statistical over-representation and under-representation compared to random sampling of GC B cells, respectively (WT Low n = 11, WT High n = 19, MT Low n = 21, MT High n = 13 cells). **e.** Fraction occurrence of MCs across conditions. (WT Control n = 1,482, MT Control n = 2,587, WT CD40L n = 1,245, MT CD40L n = 365 cells).

**Figure S10. Supplementary information for** Figure 7**. Analysis on GC microenvironmental BCs. a.** Average Z score quantified for interpretable motility and colocalization features per BC. **b.** Fractional occurrence of eight BCs across GC zones.

**Figure S11. Supplementary information for** Figure 7**. Single-cell RNA-seq analysis. a.** UMAP representation of unsupervised Leiden clustering identifies ten unique clusters (n = 10,534 cells). **b.** Customized modules scores projected onto the UMAP space. **c.** Top 10 differentially expressed genes for each Leiden cluster. **d.** Gene set association within and between literature-derived subtypes and Leiden clusters based on IoU score. Colors represent group of subtypes based on Louvain community detection. **e.** Cell subtype labels projected onto the UMAP space.

**Figure S12. Supplementary information for** Figure 7**. RNA-behavior integration analysis. a.** UMAP representation of integrated RNA-behavior space with each BC projection. **b.** BC-specific pathway association based on gene IoU score. Colors represent group of pathways based on Louvain community detection. Pathways that are not shown is not connected to any pathways.

**Figure S13. Supplementary information for** Figure 7**. Transcriptomic analysis on MC. a.** Upset plot depicting overlapping differentially expressed genes between MCs. Each column label is indicated by black dot. **b-h.** MC-specific multipartite network highlighting behavior clusters, DEGs, and enriched pathways selectively associated with MC0 (**b**), MC1 (**c**), MC3 (**d**), MC5 (**e**), MC6 (**f**), MC7 (**g**), and MC8 (**h**).

## SUPPLEMENTARY TABLES

**Table S1. List of motility features, related to Figure 1**.

**Table S2. List of colocalization features, related to Figure 1**.

**Table S3. Differentially expressed genes of MT vs WT centroblasts bulk RNA-seq, related to Figure 2**.

**Table S4. Differentially expressed genes of MT vs WT centrocytes bulk RNA-seq, related to Figure 2**.

**Table S5. GSEA pathway analysis from bulk RNA-seq of centrocytes, related to Figure 2**.

**Table S6. Motility associated gene sets, related to Figure 2**.

**Table S7. GSEA of BC as scMOTIPh validation, related to Figure 7**.

**Table S8. Differentially expressed genes of MT vs WT in BC2 after scMOTIPh mapping, related to Figure 7**.

**Table S9. Pathway community detection in BC2, related to Figure 7**.

**Table S10. Motility cluster gene signature lists after scMOTIPh mapping, related to Figure 7**.

## VIDEO CAPTIONS

**Supplementary Video 1. Intravital 2P microscopy of GC.** Imaging volumes (230 x 230 x 90 µm^3^) were acquired every 30 seconds for 90 minutes. CFP^+^ WT GC B cells (blue), YFP^+^ MT GC B cells (green), tdTomato^+^ Tfh cells (red), and AlexaFluor594-labeled CD35^+^ FDCs (gray). Section view shows 15μm-thickness maximum projections in the XY, ZY, and XZ planes. Time (hh:mm:ss).

**Supplementary Video 2. Single-cell and FDC segmentation.** Raw (left) and segmented images (right). CFP^+^ WT GC B cells (blue), YFP^+^ MT GC B cells (green), tdTomato^+^ Tfh cells (red), and AlexaFluor594-labeled CD35^+^ FDCs (gray). Scale bars, 40 μm. Time (hh:mm:ss).

**Supplementary Video 3. More localization of MT GC B cells than WT in the dense FDC networks.** All four color-detection channels (left) and only two GC B cell channels (right). CFP^+^ WT GC B cells (blue), YFP^+^ MT GC B cells (green), tdTomato^+^ Tfh cells (red), and AlexaFluor594-labeled CD35^+^ FDCs (white). Dotted white lines on the right represent the dense FDC networks region. 15 µm-thickness maximum intensity projection view. Scale bars, 20 μm.

**Supplementary Video 4. Four examples of prolonged contact of WT GC B cells with Tfh cells.** 15μm-thickness maximum projection view in the XY, ZY, and XZ planes. White lines indicate the locations of the cross sections and intersect at the example cells. CFP^+^ WT GC B cells (blue), YFP^+^ MT GC B cells (green), tdTomato^+^ Tfh cells (red), and AlexaFluor594-labeled CD35^+^ FDCs (white). Imaging volumes (230 x 230 x 90 µm^3^). In each example, time (hh:mm:ss) is set to 0 at the start of contact. All examples come from the same 90-minute imaging.

**Supplementary Video 5. Two examples of prolonged contact of MT GC B cells with Tfh cells.** 15μm-thickness maximum projection view in the XY, ZY, and XZ planes. White lines indicate the locations of the cross sections and intersect at the example cells. CFP^+^ WT GC B cells (blue), YFP^+^ MT GC B cells (green), tdTomato^+^ Tfh cells (red), and AlexaFluor594-labeled CD35^+^ FDCs (white). Imaging volumes (230 x 230 x 90 µm^3^). In each example, time (hh:mm:ss) is set to 0 at the start of contact. All examples come from the same 90-minute imaging as Supplementary Video 4.

## REFERENCES

1. Pae, J., Jacobsen, J. T. & Victora, G. D. Imaging the different timescales of germinal center selection*. Immunological Reviews vol. 306 Preprint at 10.1111/imr.13039 (2022).

2. Bannard, O. et al. Germinal center centroblasts transition to a centrocyte phenotype according to a timed program and depend on the dark zone for effective selection. Immunity 39, (2013).

3. Victora, G. D. & Nussenzweig, M. C. Germinal Centers. Annu. Rev. Immunol. 40, 413–442 (2022).

4. Young, C. & Brink, R. The unique biology of germinal center B cells. Immunity vol. 54 Preprint at 10.1016/j.immuni.2021.07.015 (2021).

5. Muppidi, J. R. & Klein, U. Directing traffic in the germinal center roundabout. Nat. Immunol. 21, (2020).

6. Allen, C. D. C., Okada, T., Tang, H. L. & Cyster, J. G. Imaging of germinal center selection events during affinity maturation. Science (1979). 315, (2007).

7. Beltman, J. B., Allen, C. D. C., Cyster, J. G. & De Boer, R. J. B cells within germinal centers migrate preferentially from dark to light zone. Proc. Natl. Acad. Sci. U. S. A. 108, (2011).

8. Schwickert, T. A. et al. In vivo imaging of germinal centres reveals a dynamic open structure. Nature 446, (2007).

9. Bödör, C. et al. EZH2 mutations are frequent and represent an early event in follicular lymphoma. Blood 122, (2013).

10. Morin, R. D. et al. Somatic mutations altering EZH2 (Tyr641) in follicular and diffuse large B-cell lymphomas of germinal-center origin. Nat. Genet. 42, (2010).

11. Okosun, J. et al. Integrated genomic analysis identifies recurrent mutations and evolution patterns driving the initiation and progression of follicular lymphoma. Nat. Genet. 46, (2014).

12. Chapuy, B. et al. Molecular subtypes of diffuse large B cell lymphoma are associated with distinct pathogenic mechanisms and outcomes. Nat. Med. 24, (2018).

13. Béguelin, W. et al. Mutant EZH2 Induces a Pre-malignant Lymphoma Niche by Reprogramming the Immune Response. Cancer Cell 37, (2020).

14. Isshiki, Y. et al. EZH2 inhibition enhances T cell immunotherapies by inducing lymphoma immunogenicity and improving T cell function. Cancer Cell https://doi.org/10.1016/j.ccell.2024.11.006 (2024) doi:10.1016/j.ccell.2024.11.006.

15. Victora, G. D. et al. Germinal center dynamics revealed by multiphoton microscopy with a photoactivatable fluorescent reporter. Cell 143, (2010).

16. Liu, D. et al. T-B-cell entanglement and ICOSL-driven feed-forward regulation of germinal centre reaction. Nature 517, (2015).

17. Shulman, Z. et al. Germinal centers: Dynamic signaling by T follicular helper cells during germinal center B cell selection. Science (1979). 345, (2014).

18. Liu, B. et al. Affinity-coupled CCL22 promotes positive selection in germinal centres. Nature 592, (2021).

19. Reimer, D. et al. B Cell Speed and B-FDC Contacts in Germinal Centers Determine Plasma Cell Output via Swiprosin-1/EFhd2. Cell Rep. 32, (2020).

20. Maity, D. et al. Profiling Dynamic Patterns of Single-Cell Motility. Advanced Science https://doi.org/10.1002/advs.202400918 (2024) doi:10.1002/advs.202400918.

21. Alieva, M., Wezenaar, A. K. L., Wehrens, E. J. & Rios, A. C. Bridging live-cell imaging and next-generation cancer treatment. Nature Reviews Cancer vol. 23 Preprint at 10.1038/s41568-023-00610-5 (2023).

22. Pittet, M. J., Garris, C. S., Arlauckas, S. P. & Weissleder, R. Recording the wild lives of immune cells. Science Immunology vol. 3 Preprint at 10.1126/sciimmunol.aaq0491 (2018).

23. Argelaguet, R., Cuomo, A. S. E., Stegle, O. & Marioni, J. C. Computational principles and challenges in single-cell data integration. Nature Biotechnology vol. 39 Preprint at 10.1038/s41587-021-00895-7 (2021).

24. Camunas-Soler, J. Integrating single-cell transcriptomics with cellular phenotypes: cell morphology, Ca2+ imaging and electrophysiology. Biophysical Reviews vol. 16 Preprint at 10.1007/s12551-023-01174-2 (2024).

25. Casola, S. et al. Tracking germinal center B cells expressing germ-line immunoglobulin γ1 transcripts by conditional gene targeting. Proc. Natl. Acad. Sci. U. S. A. 103, (2006).

26. Béguelin, W., et al. EZH2 and BCL6 Cooperate to Assemble CBX8-BCOR Complex to Repress Bivalent Promoters, Mediate Germinal Center Formation and Lymphomagenesis. Cancer Cell 30, (2016).

27. Hopt, A. & Neher, E. Highly nonlinear photodamage in two-photon fluorescence microscopy. Biophys. J. 80, (2001).

28. Zhao, Z. et al. Two-photon synthetic aperture microscopy for minimally invasive fast 3D imaging of native subcellular behaviors in deep tissue. Cell 186, (2023).

29. Choe, K. et al. Intravital three-photon microscopy allows visualization over the entire depth of mouse lymph nodes. Nat. Immunol. 23, (2022).

30. Phillip, J. M. et al. Fractional re-distribution among cell motility states during ageing. *Commun*. Biol. 4, (2021).

31. Haga, R. B. & Ridley, A. J. Rho GTPases: Regulation and roles in cancer cell biology. Small GTPases vol. 7 Preprint at 10.1080/21541248.2016.1232583 (2016).

32. Marelli-Berg, F. M. & Jangani, M. Metabolic regulation of leukocyte motility and migration. Journal of Leukocyte Biology vol. 104 Preprint at 10.1002/JLB.1MR1117-472R (2018).

33. Belliveau, N. M. et al. Whole-genome screens reveal regulators of differentiation state and context-dependent migration in human neutrophils. Nat. Commun. 14, (2023).

34. Dekkers, J. F. et al. Uncovering the mode of action of engineered T cells in patient cancer organoids. Nat. Biotechnol. 41, (2023).

35. Martínez-Riaño, A. et al. Long-term retention of antigens in germinal centers is controlled by the spatial organization of the follicular dendritic cell network. Nat. Immunol. 24, (2023).

36. Holmes, A. B. et al. Single-cell analysis of germinal-center B cells informs on lymphoma cell of origin and outcome. Journal of Experimental Medicine 217, (2020).

37. Peyré, G. & Cuturi, M. Computational Optimal Transport: With Applications to Data Science. Foundations and Trends® in Machine Learning 11, (2019).

38. Schiebinger, G. et al. Optimal-Transport Analysis of Single-Cell Gene Expression Identifies Developmental Trajectories in Reprogramming. Cell 176, (2019).

39. Klein, D. et al. Mapping cells through time and space with moscot. Nature 638, 1065–1075 (2025).

40. Allan, L. L. et al. Apolipoprotein-mediated lipid antigen presentation in B cells provides a pathway for innate help by NKT cells. Blood 114, (2009).

41. Ramiscal, R. R. & Vinuesa, C. G. T-cell subsets in the germinal center. Immunol. Rev. 252, (2013).

42. Lam, J. H. & Baumgarth, N. Toll-like receptor mediated inflammation directs B cells towards protective antiviral extrafollicular responses. Nat. Commun. 14, (2023).

43. Rookhuizen, D. C. & DeFranco, A. L. Toll-like receptor 9 signaling acts on multiple elements of the germinal center to enhance antibody responses. Proc. Natl. Acad. Sci. U. S. A. 111, (2014).

44. Defranco, A. L., Rookhuizen, D. C. & Hou, B. Contribution of Toll-like receptor signaling to germinal center antibody responses. Immunol. Rev. 247, (2012).

45. Tolar, P. Cytoskeletal control of B cell responses to antigens. Nature Reviews Immunology vol. 17 Preprint at 10.1038/nri.2017.67 (2017).

46. Kim, J. et al. Wnt5a Is Secreted by Follicular Dendritic Cells To Protect Germinal Center B Cells via Wnt/Ca2+/NFAT/NF-κB–B Cell Lymphoma 6 Signaling. The Journal of Immunology 188, (2012).

47. Bakhshi, T. J. & Georgel, P. T. Genetic and epigenetic determinants of diffuse large B-cell lymphoma. Blood Cancer Journal vol. 10 Preprint at 10.1038/s41408-020-00389-w (2020).

48. Basso, K. & Dalla-Favera, R. Germinal centres and B cell lymphomagenesis. Nat. Rev. Immunol. 15, (2015).

49. Schienstock, D. & Mueller, S. N. Moving beyond velocity: Opportunities and challenges to quantify immune cell behavior*. Immunological Reviews vol. 306 Preprint at 10.1111/imr.13038 (2022).

50. Griess, O. et al. Mutant EZH2 alters the epigenetic network and increases epigenetic heterogeneity in B cell lymphoma. PLoS Biol. 23, (2025).

51. Bunne, C. et al. How to build the virtual cell with artificial intelligence: Priorities and opportunities. Cell vol. 187 7045–7063 Preprint at 10.1016/j.cell.2024.11.015 (2024).

52. Roohani, Y., Huang, K. & Leskovec, J. Predicting transcriptional outcomes of novel multigene perturbations with GEARS. Nat. Biotechnol. 42, (2024).

53. Liu, Q., Hu, Z., Jiang, R. & Zhou, M. DeepCDR: A hybrid graph convolutional network for predicting cancer drug response. Bioinformatics 36, (2020).

54. Cui, H. et al. scGPT: toward building a foundation model for single-cell multi-omics using generative AI. Nat. Methods 21, (2024).

55. Hao, M. et al. Large-scale foundation model on single-cell transcriptomics. Nat. Methods 21, 1481–1491 (2024).

56. Morschhauser, F. et al. Taking the EZ way: Targeting enhancer of zeste homolog 2 in B-cell lymphomas. Blood Reviews vol. 56 Preprint at 10.1016/j.blre.2022.100988 (2022).

57. Srinivas, S. et al. Cre reporter strains produced by targeted insertion of EYFP and ECFP into the ROSA26 locus. BMC Dev. Biol. 1, (2001).

58. Crocker, J. C. & Grier, D. G. Methods of digital video microscopy for colloidal studies. J. Colloid Interface Sci. 179, (1996).

59. Pachitariu, M., Rariden, M. & Stringer, C. Cellpose-SAM: superhuman generalization for cellular segmentation. bioRxiv (2025).

60. Wu, P. H., Giri, A. & Wirtz, D. Statistical analysis of cell migration in 3D using the anisotropic persistent random walk model. Nat. Protoc. 10, (2015).

61. Wu, P. H., Giri, A., Sun, S. X. & Wirtz, D. Three-dimensional cell migration does not follow a random walk. Proc. Natl. Acad. Sci. U. S. A. 111, (2014).

62. Becht, E. et al. Dimensionality reduction for visualizing single-cell data using UMAP. Nat. Biotechnol. 37, (2019).

63. Lopez, R., Regier, J., Cole, M. B., Jordan, M. I. & Yosef, N. Deep generative modeling for single-cell transcriptomics. Nat. Methods 15, (2018).

64. Bernstein, N. J. et al. Solo: Doublet Identification in Single-Cell RNA-Seq via Semi-Supervised Deep Learning. Cell Syst. 11, (2020).

65. Wolf, F. A., Angerer, P. & Theis, F. J. SCANPY: Large-scale single-cell gene expression data analysis. Genome Biol. 19, (2018).

66. Vayer, T., Chapel, L., Flamary, R., Tavenard, R. & Courty, N. Fused gromov-wasserstein distance for structured objects. Algorithms 13, (2020).

