## Supplementary Figures for "Intravital single-cell behavior profiling reveals disrupted germinal center B cell motility and interactions by EZH2 gain-of-function mutation"

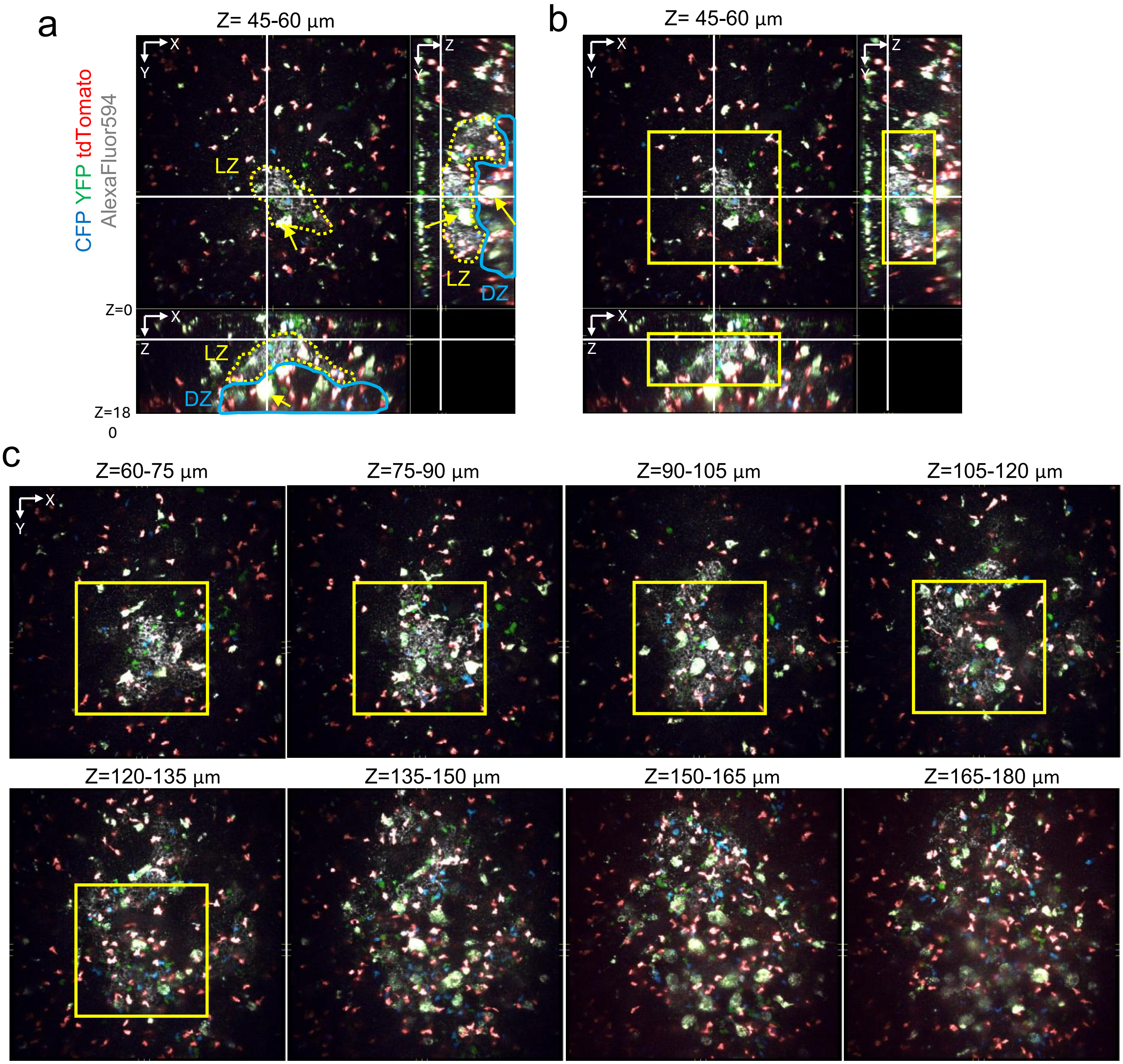
a
Z= 45-60 µm
CFP YFP tdTomato
AlexaFluor594
X
Y
Z
LZ
DZ
Z=0
Z=18
0
b
Z= 45-60 µm
c
Z=60-75 µm
Z=75-90 µm
Z=90-105 µm
Z=105-120 µm
Z=120-135 µm
Z=135-150 µm
Z=150-165 µm
Z=165-180 µm

**Figure S1**

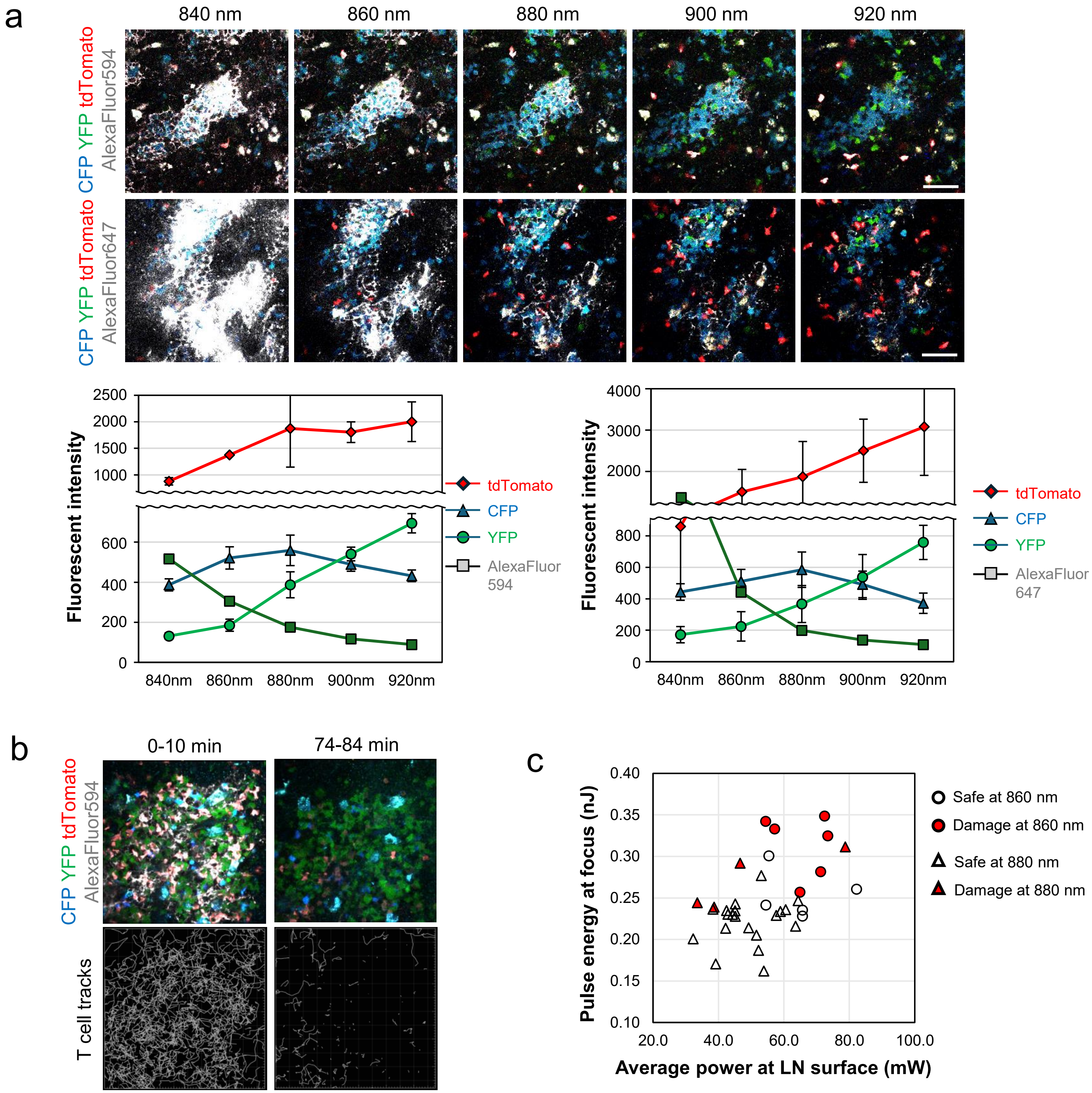
a
840 nm
860 nm
880 nm
900 nm
920 nm
CFP YFP tdTomato AlexaFluor594
CFP YFP tdTomato AlexaFluor647
Fluorescent intensity
tdTomato
CFP
YFP
AlexaFluor 594
AlexaFluor 647
840nm
860nm
880nm
900nm
920nm
b
0-10 min
74-84 min
CFP YFP tdTomato AlexaFluor594
T cell tracks
c
Safe at 860 nm
Damage at 860 nm
Safe at 880 nm
Damage at 880 nm
Pulse energy at focus (nJ)
Average power at LN surface (mW)

### Figure S2

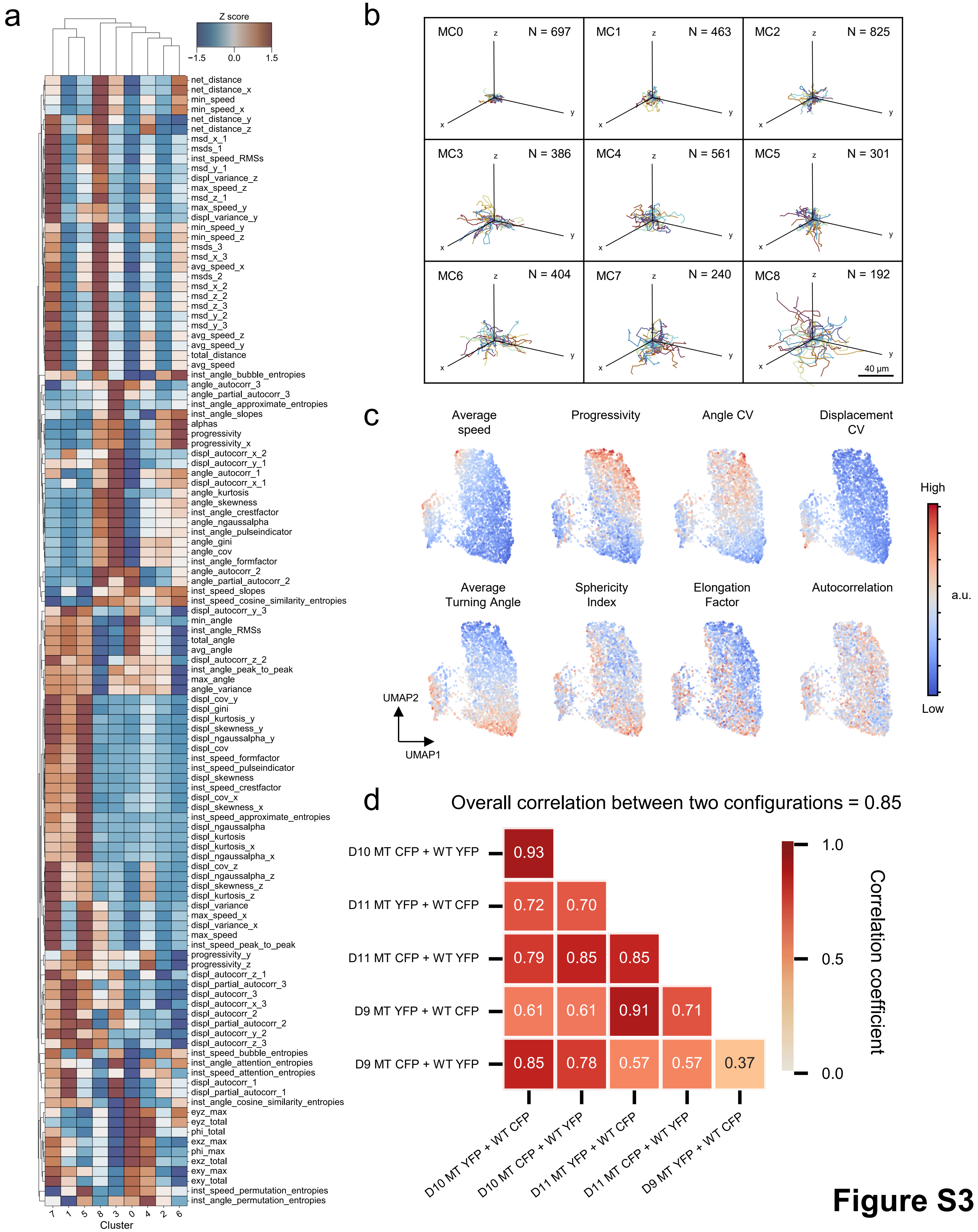
a
Z score
−1.5
0.0
1.5
net_distance
net_distance_x
min_speed
min_speed_x
net_distance_y
net_distance_z
msd_x_1
msds_1
inst_speed_RMSs
msd_y_1
displ_variance_z
max_speed_z
msd_z_1
max_speed_y
displ_variance_y
min_speed_y
min_speed_z
msds_3
msd_x_3
avg_speed_x
msds_2
msd_x_2
msd_z_2
msd_z_3
msd_y_2
msd_y_3
avg_speed_z
avg_speed_y
total_distance
avg_speed
inst_angle_bubble_entropies
angle_autocorr_3
angle_partial_autocorr_3
inst_angle_approximate_entropies
inst_angle_slopes
alphas
progressivity
progressivity_x
displ_autocorr_x_2
displ_autocorr_y_1
angle_autocorr_1
displ_autocorr_x_1
angle_kurtosis
angle_skewness
inst_angle_crestfactor
angle_ngaussalpha
inst_angle_pulseindicator
angle_gini
angle_cov
inst_angle_formfactor
angle_autocorr_2
angle_partial_autocorr_2
inst_speed_slopes
inst_speed_cosine_similarity_entropies
displ_autocorr_y_3
min_angle
inst_angle_RMSs
total_angle
avg_angle
displ_autocorr_z_2
inst_angle_peak_to_peak
max_angle
angle_variance
displ_cov_y
displ_gini
displ_kurtosis_y
displ_skewness_y
displ_ngaussalpha_y
displ_cov
inst_speed_formfactor
inst_speed_pulseindicator
displ_skewness
inst_speed_crestfactor
displ_cov_x
displ_skewness_x
inst_speed_approximate_entropies
displ_ngaussalpha
displ_kurtosis
displ_kurtosis_x
displ_ngaussalpha_x
displ_cov_z
displ_ngaussalpha_z
displ_skewness_z
displ_kurtosis_z
displ_variance
max_speed_x
displ_variance_x
max_speed
inst_speed_peak_to_peak
progressivity_y
progressivity_z
displ_autocorr_z_1
displ_partial_autocorr_3
displ_autocorr_3
displ_autocorr_x_3
displ_autocorr_2
displ_partial_autocorr_2
displ_autocorr_y_2
displ_autocorr_z_3
inst_speed_bubble_entropies
inst_angle_attention_entropies
inst_speed_attention_entropies
displ_autocorr_1
displ_partial_autocorr_1
inst_angle_cosine_similarity_entropies
eyz_max
eyz_total
phi_total
exz_max
phi_max
exz_total
exy_max
exy_total
inst_speed_permutation_entropies
inst_angle_permutation_entropies
7 1 5 8 3 0 4 2 6
Cluster
b
MC0 N = 697
MC1 N = 463
MC2 N = 825
MC3 N = 386
MC4 N = 561
MC5 N = 301
MC6 N = 404
MC7 N = 240
MC8 N = 192
x y z
40 μm
c
Average speed
Progressivity
Angle CV
Displacement CV
Average Turning Angle
Sphericity Index
Elongation Factor
Autocorrelation
High
a.u.
Low
UMAP2
UMAP1
d
Overall correlation between two configurations = 0.85
D10 MT CFP + WT YFP
D11 MT YFP + WT CFP
D11 MT CFP + WT YFP
D9 MT YFP + WT CFP
D9 MT CFP + WT YFP
0.93
0.72 0.70
0.79 0.85 0.85
0.61 0.61 0.91 0.71
0.85 0.78 0.57 0.57 0.37
D10 MT YFP + WT CFP
D10 MT CFP + WT YFP
D11 MT YFP + WT CFP
D11 MT CFP + WT YFP
D9 MT YFP + WT CFP
1.0
0.5
0.0
Correlation coefficient

**Figure S3**

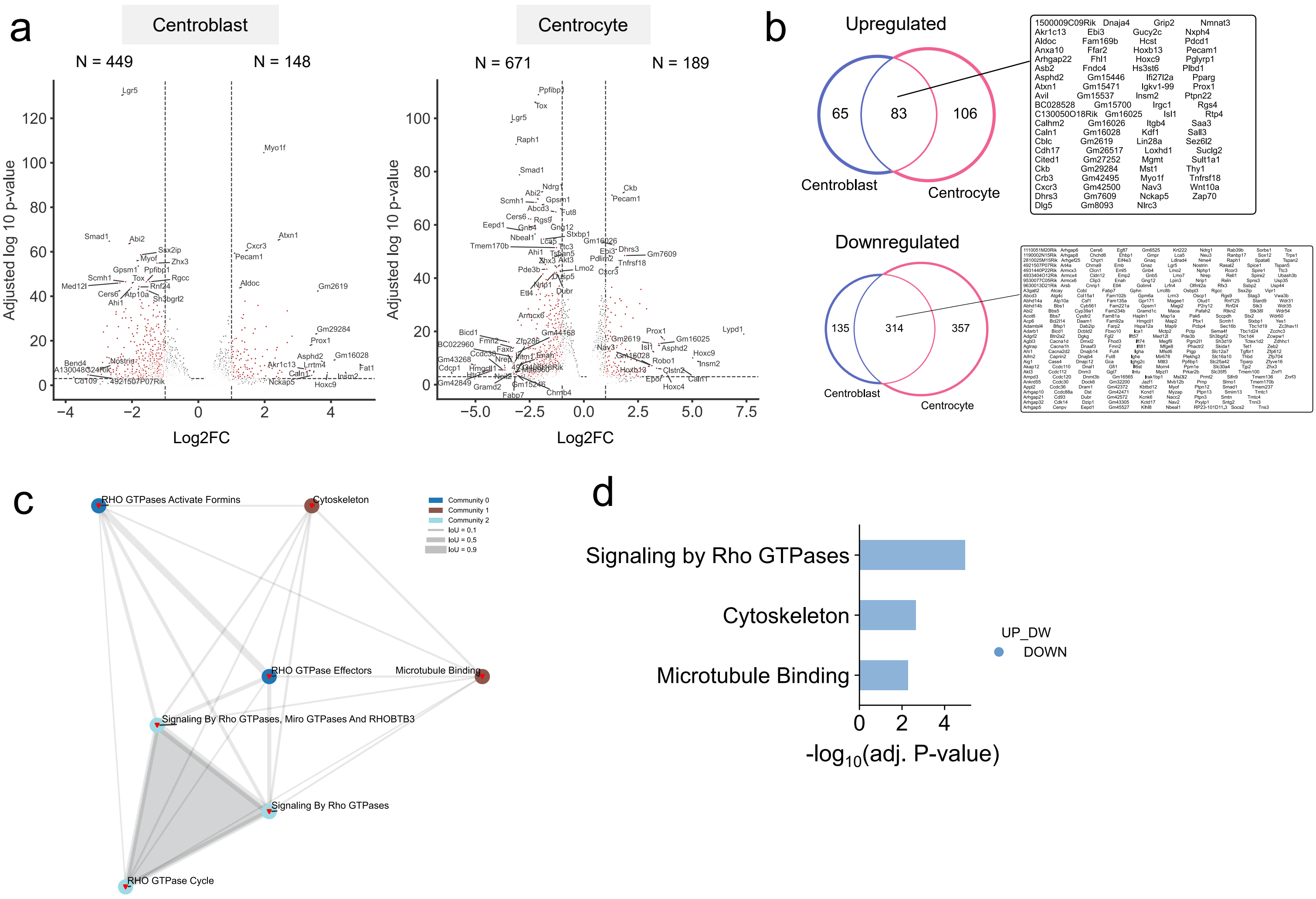

a
Centroblast
Centrocyte
N = 449
N = 148
N = 671
N = 189
Adjusted log 10 p-value
Log2FC
b
Upregulated
65
83
106
Centroblast
Centrocyte
1500009C09Rik Dnaja4 Grip2 Nmnat3
Akr1c13 Ebi3 Gucy2c Nxph4
Aldoc Fam169b Hcst Pdcd1
Anxa10 Ffar2 Hoxb13 Pecam1
Arhgap22 Fhl1 Hoxc9 Pglyrp1
Asb2 Fndc4 Hs3st6 Plbd1
Asphd2 Gm15446 Ifi27l2a Pparg
Atxn1 Gm15471 Igkv1-99 Prox1
Avil Gm15537 Insm2 Ptpn22
BC028528 Gm15700 Irgc1 Rgs4
C130050O18Rik Gm16025 Isl1 Rtp4
Calhm2 Gm16026 Itgb4 Saa3
Caln1 Gm16028 Kdf1 Sall3
Cblc Gm2619 Lin28a Sez6l2
Cdh17 Gm26517 Loxhd1 Suclg2
Cited1 Gm27252 Mgmt Sult1a1
Ckb Gm29284 Mst1 Thy1
Crb3 Gm42495 Myo1f Tnfrsf18
Cxcr3 Gm42500 Nav3 Wnt10a
Dhrs3 Gm7609 Nckap5 Zap70
Dlg5 Gm8093 Nlrc3
Downregulated
135
314
357
Centroblast
Centrocyte
c
RHO GTPases Activate Formins
Cytoskeleton
RHO GTPase Effectors
Microtubule Binding
Signaling By Rho GTPases, Miro GTPases And RHOBTB3
Signaling By Rho GTPases
RHO GTPase Cycle
Community 0
Community 1
Community 2
IoU = 0.1
IoU = 0.5
IoU = 0.9
d
Signaling by Rho GTPases
Cytoskeleton
Microtubule Binding
UP_DW
DOWN
-log10(adj. P-value)


**Figure S4**

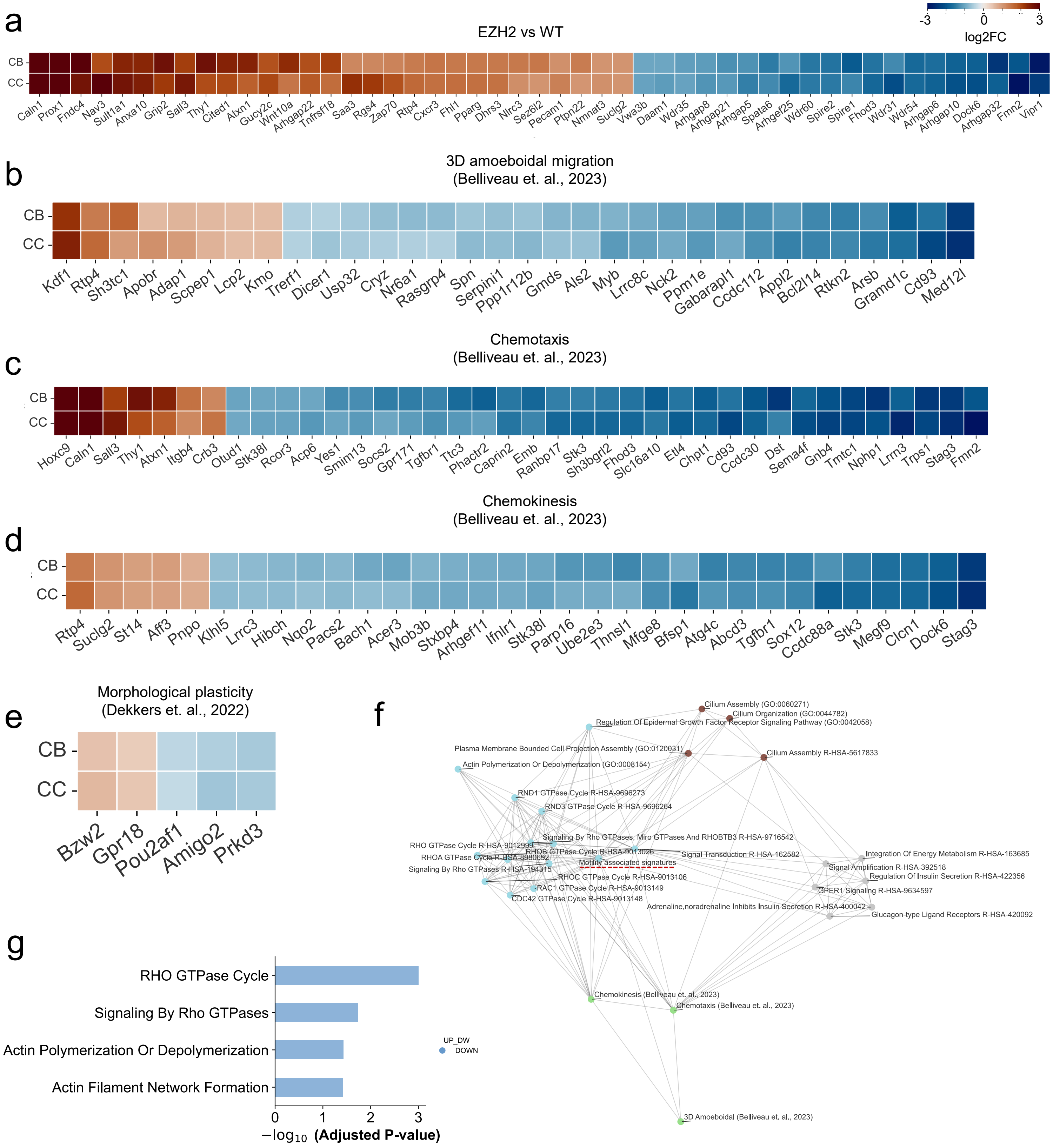

a
EZH2 vs WT
-3
0
3
log2FC
CB
CC
Caln1 Prox1 Fndc4 Nav3 Sult1a1 Anxa10 Grip2 Sall3 Thy1 Cited1 Atxn1 Gucy2c Wnt10a Arhgap22 Tnfrsf18 Saa3 Rgs4 Zap70 Rtp4 Cxcr3 Fhl1 Pparg Dhrs3 Nlrc3 Sez6l2 Pecam1 Ptpn22 Nmnat3 Suclg2 Vwa3b Daam1 Wdr35 Arhgap8 Arhgap21 Arhgap5 Spata6 Arhgef25 Wdr60 Spire2 Spire1 Fhod3 Wdr31 Wdr54 Arhgap6 Arhgap10 Dock6 Arhgap32 Fmn2 Vipr1
b
3D amoeboidal migration
(Belliveau et. al., 2023)
CB
CC
Kdf1 Rtp4 Sh3tc1 Apobr Adap1 Scpep1 Lcp2 Kmo Trerf1 Dicer1 Usp32 Cryz Nr6a1 Rasgrp4 Spn Serpini1 Ppp1r12b Gmds Als2 Myb Lrrc8c Nck2 Ppm1e Gabarapl1 Ccdc112 Appl2 Bcl2l14 Rtkn2 Arsb Gramd1c Cd93 Med12l
c
Chemotaxis
(Belliveau et. al., 2023)
CB
CC
Hoxc9 Caln1 Sall3 Thy1 Atxn1 Itgb4 Crb3 Otud1 Stk38l Rcor3 Acp6 Yes1 Smim13 Socs2 Gpr171 Tgfbr1 Ttc3 Phactr2 Caprin2 Emb Ranbp17 Stk3 Sh3bgrl2 Fhod3 Slc16a10 Etl4 Chpt1 Cd93 Ccdc30 Dst Sema4f Gnb4 Tmtc1 Nphp1 Lrrn3 Trps1 Stag3 Fmn2
d
Chemokinesis
(Belliveau et. al., 2023)
CB
CC
Rtp4 Suclg2 St14 Aff3 Pnpo Klhl5 Lrrc3 Hibch Nqo2 Pacs2 Bach1 Acer3 Mob3b Stxbp4 Arhgef11 Ifnlr1 Stk38l Parp16 Ube2e3 Thnsl1 Mfge8 Bfsp1 Atg4c Abcd3 Tgfbr1 Sox12 Ccdc88a Stk3 Megf9 Clcn1 Dock6 Stag3
e
Morphological plasticity
(Dekkers et. al., 2022)
CB
CC
Bzw2 Gpr18 Pou2af1 Amigo2 Prkd3
f
Cilium Assembly (GO:0060271)
Cilium Organization (GO:0044782)
Regulation Of Epidermal Growth Factor Receptor Signaling Pathway (GO:0042058)
Plasma Membrane Bounded Cell Projection Assembly (GO:0120031)
Cilium Assembly R-HSA-5617833
Actin Polymerization Or Depolymerization (GO:0008154)
RND1 GTPase Cycle R-HSA-9696273
RND3 GTPase Cycle R-HSA-9696264
Signaling By Rho GTPases, Miro GTPases And RHOBTB3 R-HSA-9716542
RHO GTPase Cycle R-HSA-9012999
RHOB GTPase Cycle R-HSA-9013026
Signal Transduction R-HSA-162582
Integration Of Energy Metabolism R-HSA-163685
RHOA GTPase Cycle R-HSA-8980692
Motility associated signatures
Signal Amplification R-HSA-392518
Signaling By Rho GTPases R-HSA-194315
RHOC GTPase Cycle R-HSA-9013106
Regulation Of Insulin Secretion R-HSA-422356
RAC1 GTPase Cycle R-HSA-9013149
GPER1 Signaling R-HSA-9634597
CDC42 GTPase Cycle R-HSA-9013148
Adrenaline,noradrenaline Inhibits Insulin Secretion R-HSA-400042
Glucagon-type Ligand Receptors R-HSA-420092
Chemokinesis (Belliveau et. al., 2023)
Chemotaxis (Belliveau et. al., 2023)
3D Amoeboidal (Belliveau et. al., 2023)
g
RHO GTPase Cycle
Signaling By Rho GTPases
Actin Polymerization Or Depolymerization
Actin Filament Network Formation
UP_DW
DOWN
0
1
2
3
$-\log_{10}$ (Adjusted P-value)


### Figure S5

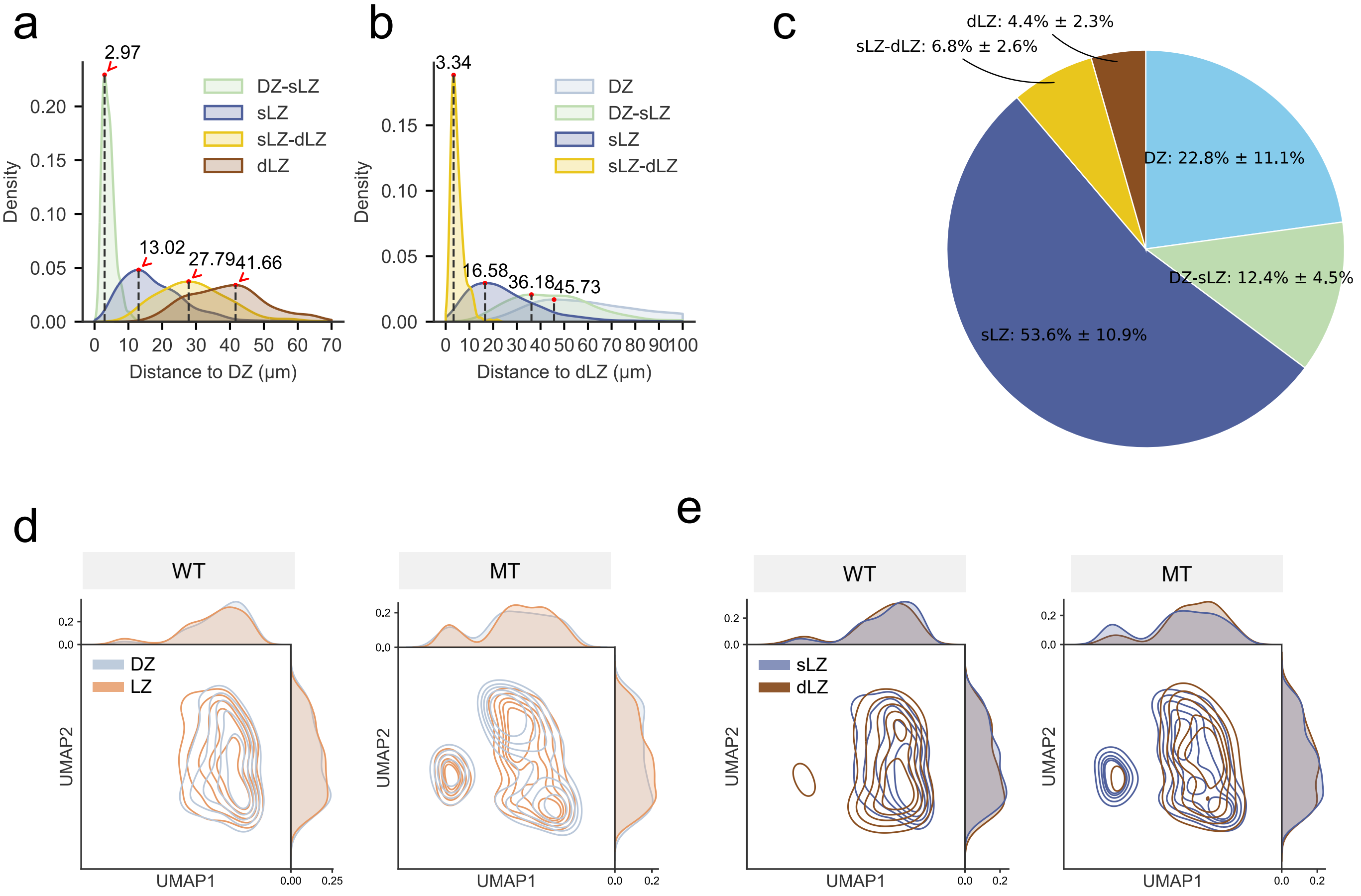

a
2.97
13.02
27.79
41.66
DZ-sLZ
sLZ
sLZ-dLZ
dLZ
Density
Distance to DZ (μm)
b
3.34
16.58
36.18
45.73
DZ
DZ-sLZ
sLZ
sLZ-dLZ
Density
Distance to dLZ (μm)
c
dLZ: 4.4% ± 2.3%
sLZ-dLZ: 6.8% ± 2.6%
DZ: 22.8% ± 11.1%
DZ-sLZ: 12.4% ± 4.5%
sLZ: 53.6% ± 10.9%
d
WT
MT
DZ
LZ
UMAP1
UMAP2
e
WT
MT
sLZ
dLZ
UMAP1
UMAP2


### Figure S6

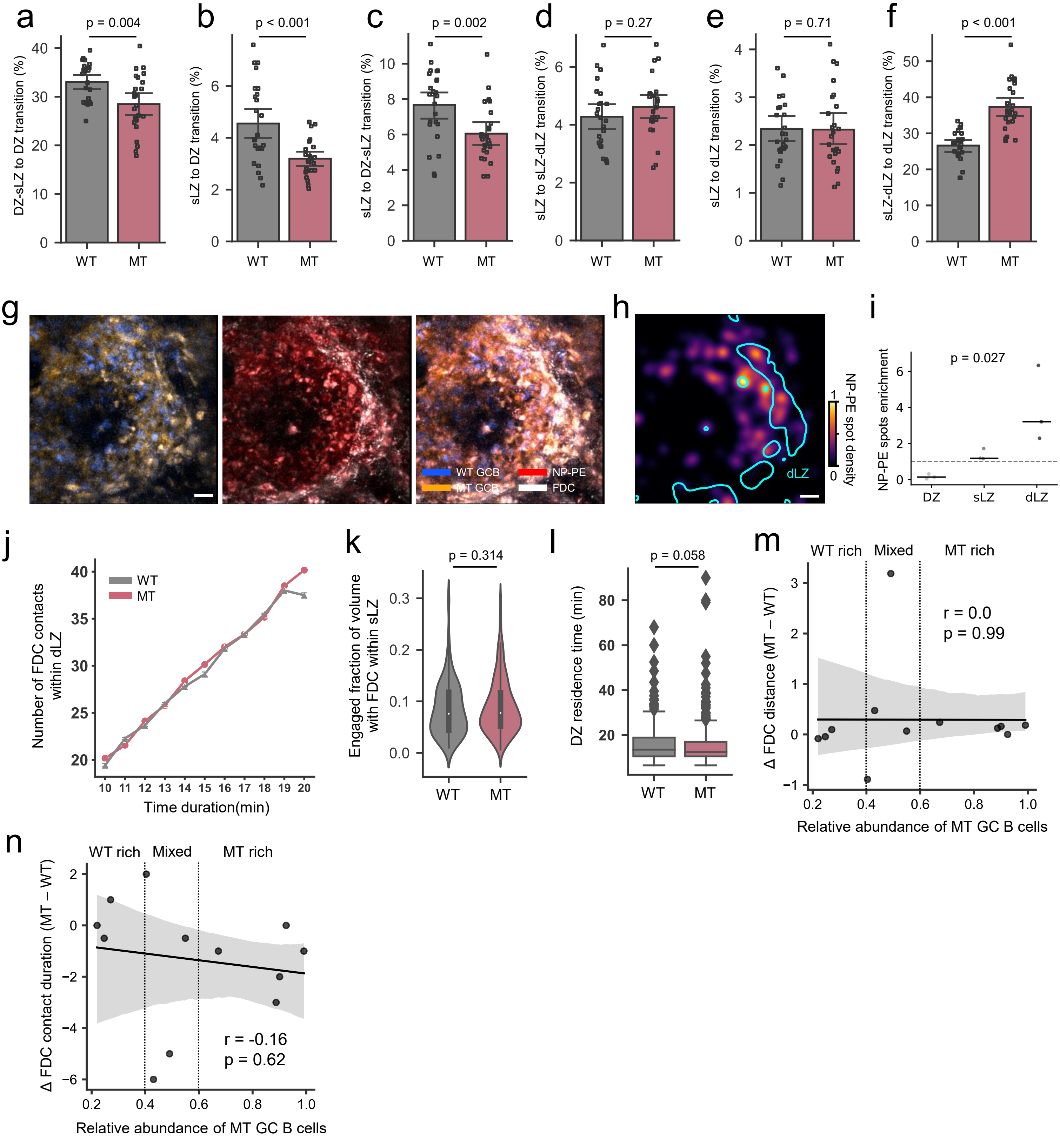

a
p = 0.004
DZ-sLZ to DZ transition (%)
WT
MT
b
p < 0.001
sLZ to DZ transition (%)
WT
MT
c
p = 0.002
sLZ to DZ-sLZ transition (%)
WT
MT
d
p = 0.27
sLZ to sLZ-dLZ transition (%)
WT
MT
e
p = 0.71
sLZ to dLZ transition (%)
WT
MT
f
p < 0.001
sLZ-dLZ to dLZ transition (%)
WT
MT
g
WT GCB
MT GCB
NP-PE
FDC
h
dLZ
NP-PE spot density
i
p = 0.027
NP-PE spots enrichment
DZ
sLZ
dLZ
j
WT
MT
Number of FDC contacts within dLZ
Time duration(min)
k
p = 0.314
Engaged fraction of volume with FDC within sLZ
WT
MT
l
p = 0.058
DZ residence time (min)
WT
MT
m
WT rich
Mixed
MT rich
r = 0.0
p = 0.99
Δ FDC distance (MT − WT)
Relative abundance of MT GC B cells
n
WT rich
Mixed
MT rich
r = -0.16
p = 0.62
Δ FDC contact duration (MT − WT)
Relative abundance of MT GC B cells


**Figure S7**

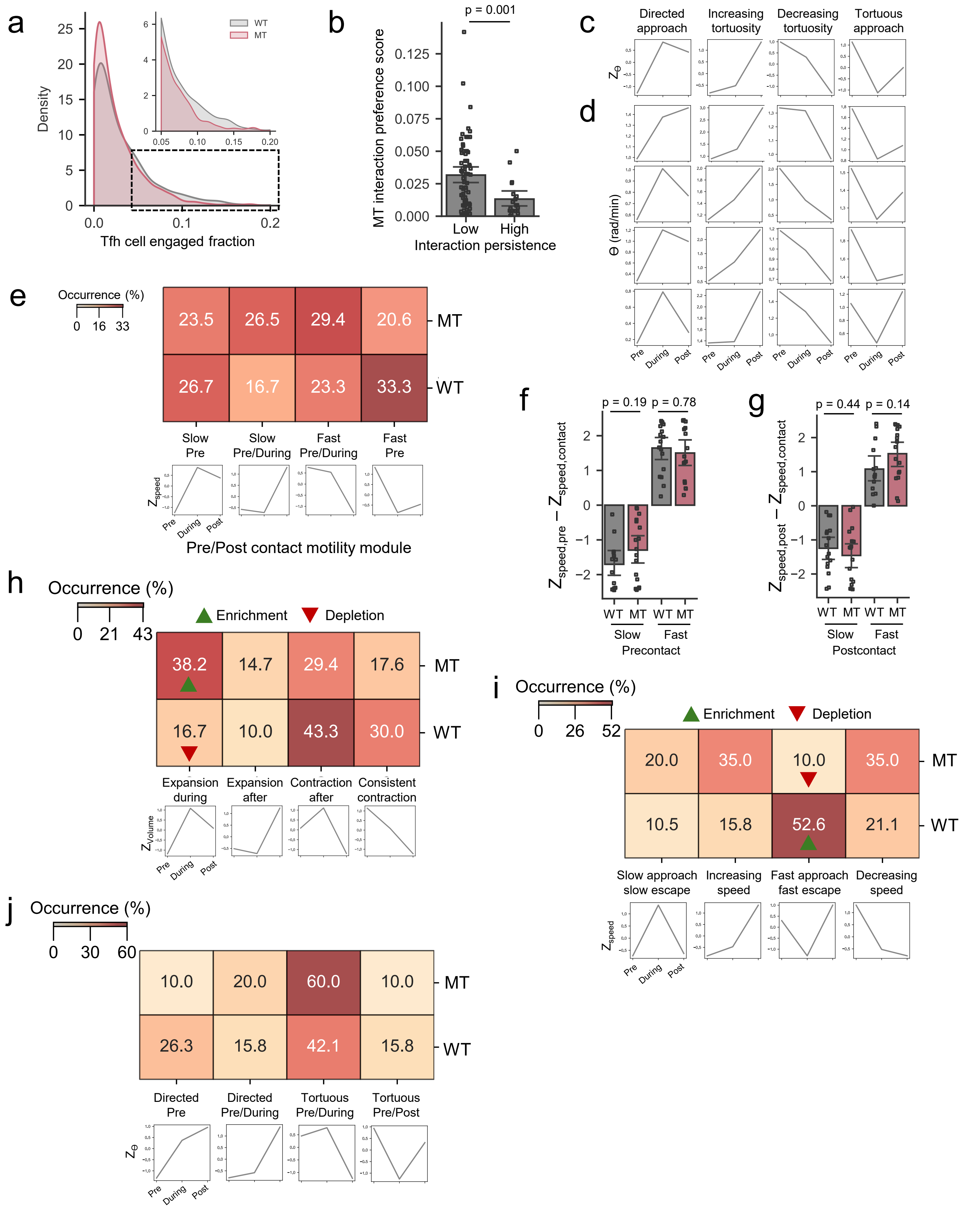
a
Density
Tfh cell engaged fraction
WT
MT
b
p = 0.001
MT interaction preference score
Low
High
Interaction persistence
c
Directed approach
Increasing tortuosity
Decreasing tortuosity
Tortuous approach
d
θ (rad/min)
e
Occurrence (%)
Slow Pre
Slow Pre/During
Fast Pre/During
Fast Pre
Pre/Post contact motility module
f
p = 0.19
p = 0.78
Precontact
g
p = 0.44
p = 0.14
Postcontact
h
Occurrence (%)
Enrichment
Depletion
Expansion during
Expansion after
Contraction after
Consistent contraction
i
Occurrence (%)
Enrichment
Depletion
Slow approach slow escape
Increasing speed
Fast approach fast escape
Decreasing speed
j
Occurrence (%)
Directed Pre
Directed Pre/During
Tortuous Pre/During
Tortuous Pre/Post

### Figure S8

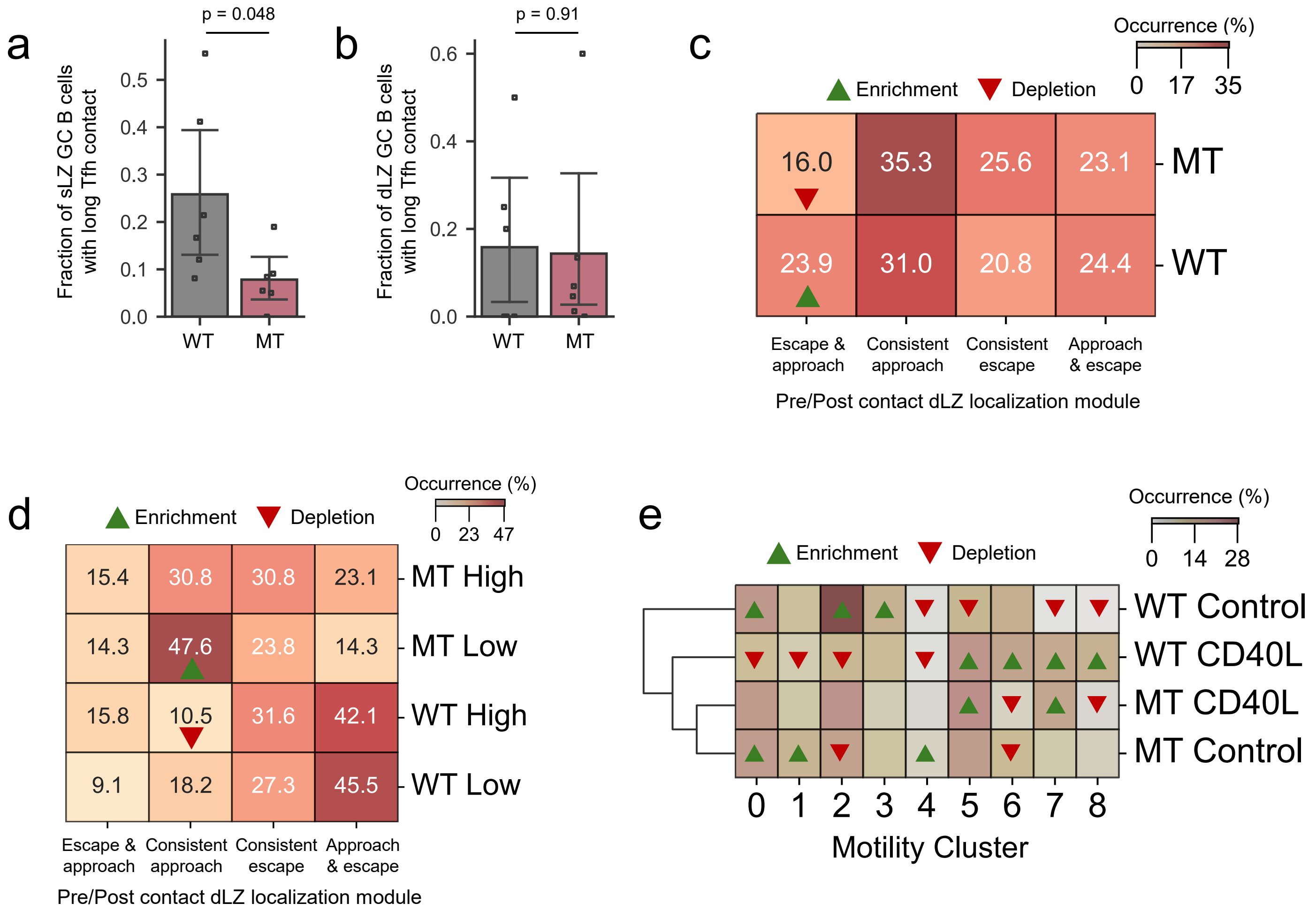
a
p = 0.048
Fraction of sLZ GC B cells with long Tfh contact
0.0
0.1
0.2
0.3
0.4
0.5
WT
MT
b
p = 0.91
Fraction of dLZ GC B cells with long Tfh contact
0.0
0.2
0.4
0.6
WT
MT
c
Occurrence (%)
0 17 35
Enrichment
Depletion
16.0
35.3
25.6
23.1
MT
23.9
31.0
20.8
24.4
WT
Escape & approach
Consistent approach
Consistent escape
Approach & escape
Pre/Post contact dLZ localization module
d
Enrichment
Depletion
Occurrence (%)
0 23 47
15.4
30.8
30.8
23.1
MT High
14.3
47.6
23.8
14.3
MT Low
15.8
10.5
31.6
42.1
WT High
9.1
18.2
27.3
45.5
WT Low
Escape & approach
Consistent approach
Consistent escape
Approach & escape
Pre/Post contact dLZ localization module
e
Occurrence (%)
0 14 28
Enrichment
Depletion
WT Control
WT CD40L
MT CD40L
MT Control
0 1 2 3 4 5 6 7 8
Motility Cluster

### Figure S9

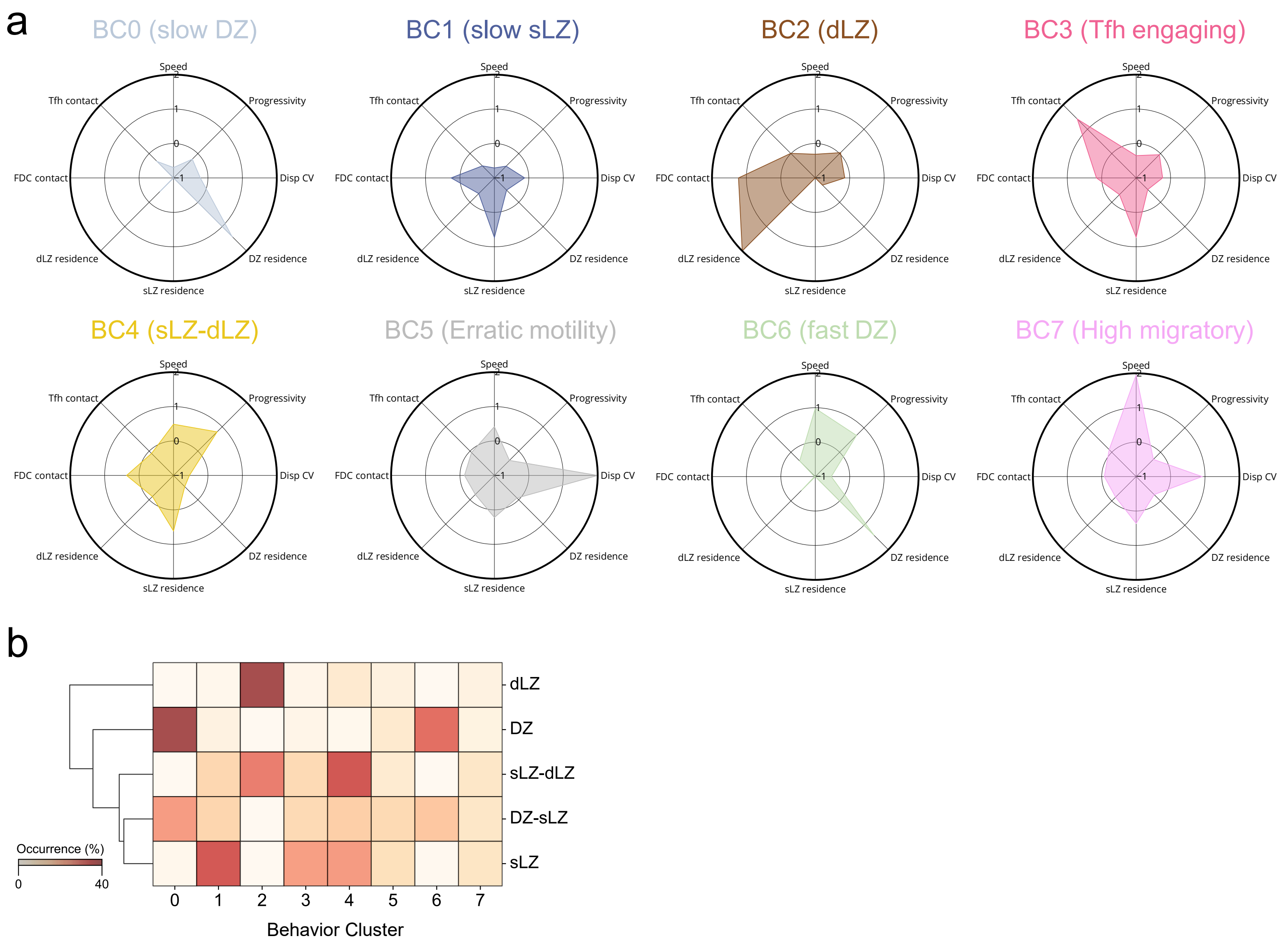
a
BC0 (slow DZ)
BC1 (slow sLZ)
BC2 (dLZ)
BC3 (Tfh engaging)
BC4 (sLZ-dLZ)
BC5 (Erratic motility)
BC6 (fast DZ)
BC7 (High migratory)
Speed
Progressivity
Disp CV
DZ residence
sLZ residence
dLZ residence
FDC contact
Tfh contact
b
dLZ
DZ
sLZ-dLZ
DZ-sLZ
sLZ
Occurrence (%)
0
40
0 1 2 3 4 5 6 7
Behavior Cluster

**Figure S10**

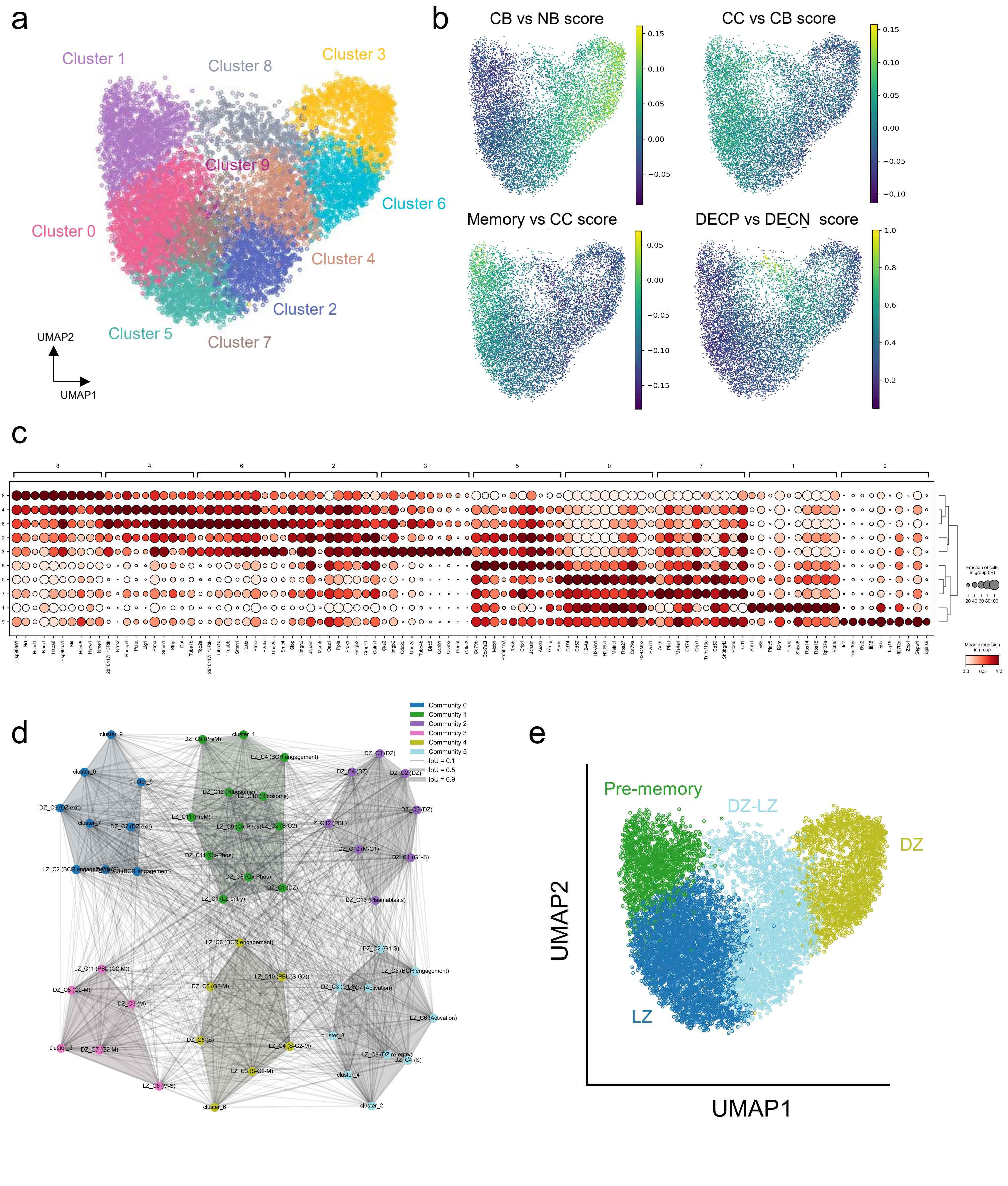
a
Cluster 1
Cluster 8
Cluster 3
Cluster 9
Cluster 6
Cluster 0
Cluster 4
Cluster 2
Cluster 5
Cluster 7
UMAP2
UMAP1
b
CB vs NB score
CC vs CB score
Memory vs CC score
DECP vs DECN score
c
Fraction of cells in group (%)
Mean expression in group
d
Community 0
Community 1
Community 2
Community 3
Community 4
Community 5
IoU = 0.1
IoU = 0.5
IoU = 0.9
e
Pre-memory
DZ-LZ
DZ
LZ
UMAP2
UMAP1

### Figure S11

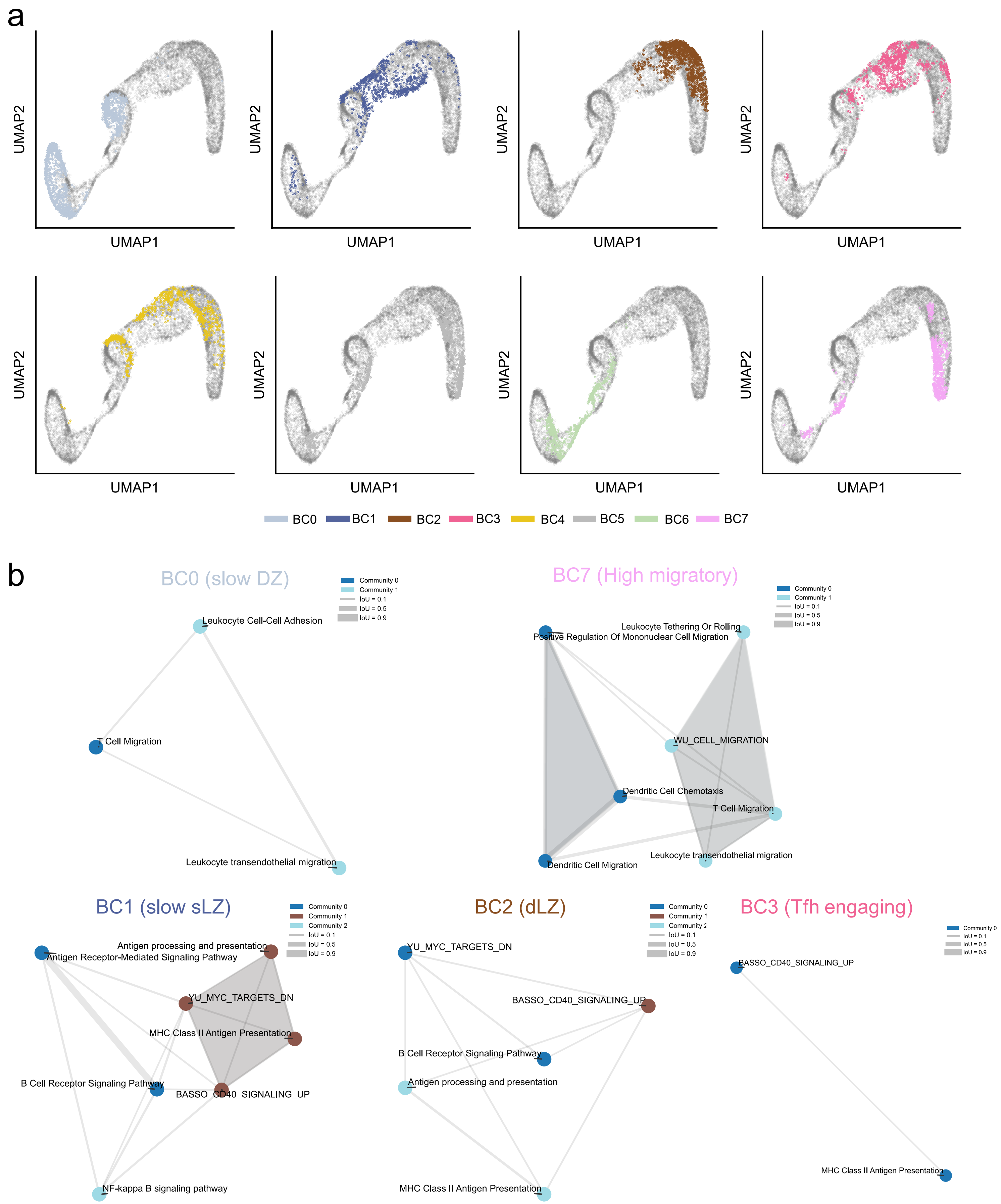
a
UMAP2
UMAP1
BC0
BC1
BC2
BC3
BC4
BC5
BC6
BC7
b
BC0 (slow DZ)
Community 0
Community 1
IoU = 0.1
IoU = 0.5
IoU = 0.9
Leukocyte Cell-Cell Adhesion
T Cell Migration
Leukocyte transendothelial migration
BC7 (High migratory)
Leukocyte Tethering Or Rolling
Positive Regulation Of Mononuclear Cell Migration
WU_CELL_MIGRATION
Dendritic Cell Chemotaxis
T Cell Migration
Leukocyte transendothelial migration
Dendritic Cell Migration
BC1 (slow sLZ)
Community 2
Antigen processing and presentation
Antigen Receptor-Mediated Signaling Pathway
YU_MYC_TARGETS_DN
MHC Class II Antigen Presentation
B Cell Receptor Signaling Pathway
BASSO_CD40_SIGNALING_UP
NF-kappa B signaling pathway
BC2 (dLZ)
YU_MYC_TARGETS_DN
BASSO_CD40_SIGNALING_UP
B Cell Receptor Signaling Pathway
Antigen processing and presentation
MHC Class II Antigen Presentation
BC3 (Tfh engaging)
BASSO_CD40_SIGNALING_UP
MHC Class II Antigen Presentation

**Figure S12**

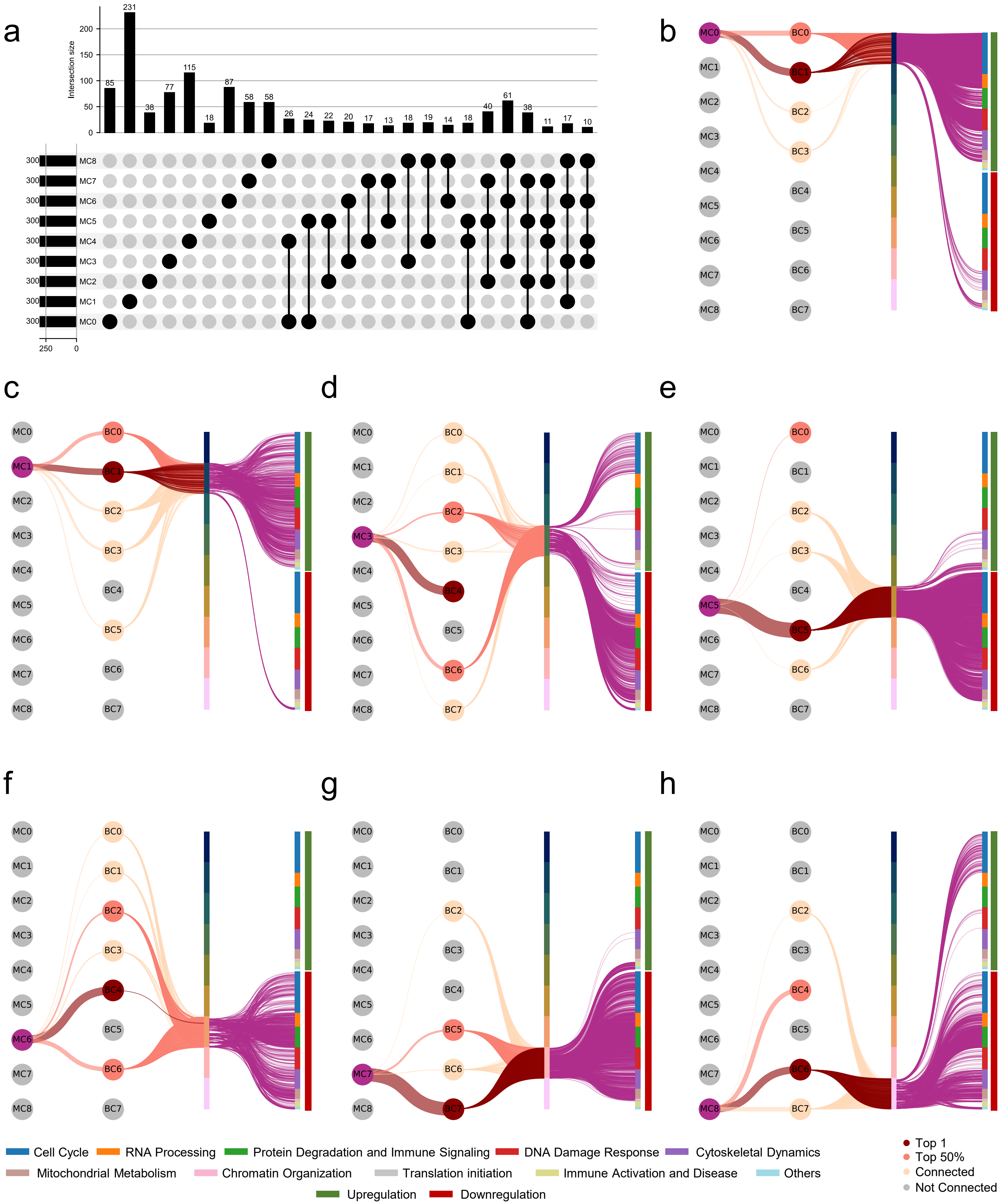
a
Intersection size
MC0
MC1
MC2
MC3
MC4
MC5
MC6
MC7
MC8
b
c
d
e
f
g
h
BC0
BC1
BC2
BC3
BC4
BC5
BC6
BC7
Cell Cycle
RNA Processing
Protein Degradation and Immune Signaling
DNA Damage Response
Cytoskeletal Dynamics
Mitochondrial Metabolism
Chromatin Organization
Translation initiation
Immune Activation and Disease
Others
Upregulation
Downregulation
Top 1
Top 50%
Connected
Not Connected

**Figure S13**
